# Genomic deletion and duplication effects on cognitive ability lie at the opposing ends of the cortical sensorimotor–association hierarchy

**DOI:** 10.1101/2025.01.06.631492

**Authors:** Kuldeep Kumar, Sayeh Kazem, Jane Yang, Florian Benitiere, Mame Seynabou Diop, Benjamin Clark, Guillaume Huguet, Worrawat Engchuan, Jakub Kopal, Thomas Renne, Omar Shanta, Bhooma Thiruvahindrapuram, Jeffrey R. MacDonald, Josephine Mollon, Laura M. Schultz, Emma E.M. Knowles, David Porteous, Gail Davies, Paul Redmond, Sarah E. Harris, Simon R. Cox, Gunter Schumann, Zdenka Pausova, Celia M. T. Greenwood, Tomas Paus, Stephen W. Scherer, Laura Almasy, Jonathan Sebat, David C Glahn, Guillaume Dumas, Sebastien Jacquemont

**Affiliations:** Centre de recherche CHU Sainte-Justine and University of Montreal, Montreal, QC, Canada; The Hospital for Sick Children, Genetics and Genome Biology, Toronto, ON, Canada; The Hospital for Sick Children, The Centre for Applied Genomics, Toronto, ON, Canada; Centre for Precision Psychiatry, Division of Mental Health and Addiction, Oslo University Hospital & Institute of Clinical Medicine, University of Oslo, Oslo, Norway; University of California, San Diego, Department of Psychiatry, Department of Cellular & Molecular Medicine, Beyster Center of Psychiatric Genomics, San Diego, CA, USA; Department of Psychiatry, Boston Children’s Hospital, Harvard Medical School, Boston, MA, USA; Department of Biomedical and Health Informatics, The Children’s Hospital of Philadelphia, Philadelphia, PA, USA; Lothian Birth Cohorts, Department of Psychology, School of Philosophy, Psychology and Language Sciences, The University of Edinburgh, Edinburgh, UK; Medical Genetics Section, Centre for Genomic & Experimental Medicine, MRC Institute of Genetics & Molecular Medicine, University of Edinburgh, Western General Hospital, Edinburgh, UK; Generation Scotland, Centre for Genomic and Experimental Medicine, University of Edinburgh, Edinburgh, UK; Institute of Psychiatry, Psychology, and Neuroscience, King’s College London, London, UK; Department of Pediatrics, University of Montreal, Montreal, Canada; ECOGENE-21, Chicoutimi, QC, Canada; Lady Davis Institute for Medical Research, Jewish General Hospital, Montreal, QC, Canada; Departments of Oncology, Epidemiology, Biostatistics, and Occupational Health, and Division of Cancer Epidemiology, McGill University, Montreal, QC, Canada; Department of Psychiatry and Addictology, Department of Neuroscience, Faculty of Medicine, University of Montreal, Montreal, QC, Canada; McLaughlin Centre and Department of Molecular Genetics, University of Toronto, Toronto, ON, Canada; Lifespan Brain Institute, Children’s Hospital of Philadelphia, and Penn Medicine, PA, USA; Boston Children’s Hospital, Tommy Fuss Center for Neuropsychiatric Disease Research, Boston, MA, USA; Mila – Quebec Artificial Intelligence Institute, Montreal, QC, Canada; Barcelonabeta Brain Research Centre, Pasqual Maragall Foundation, Barcelona, Spain

**Author notes:** authors contributed equally (Kuldeep Kumar, Sayeh Kazem). Shared senior authorship (Guillaume Dumas, Sebastien Jacquemont).

## Abstract

Cognitive deficits are common across many neurodevelopmental and psychiatric conditions, including those studied in the current set of PGC-CNV papers. How changes in regional gene expression across the cerebral cortex influence cognitive ability remains unknown. Population variation in gene dosage, which significantly impacts gene expression, represents a unique paradigm to address this question. We developed a cerebral-cortex gene-set burden analysis (CC-GSBA) to associate a trait with genomic deletions and duplications that disrupt genes with similar expression profiles across 180 cortical regions. We performed CC-GSBA across 180 cortical regions to test associations with cognitive ability in 258,292 individuals from general population cohorts. Most cortical gene sets were associated with a decrease in cognitive ability when deleted or duplicated, and this approach revealed opposing cortical patterns for the effect sizes of deletions and duplications. These cortical patterns of effect sizes followed the cortical hierarchy previously characterized at the molecular, cellular, and functional levels. We show that genes with preferential expression in sensorimotor regions demonstrated the largest effect on cognition when deleted. At the opposing end of the cortical hierarchy, genes with preferential expression in multimodal association regions affected cognition the most when duplicated. This cortical pattern of gene dosage response could not be explained by particular cell types, developmental epochs, or genetic constraints, highlighting the fact that the macroscopic network organization of the cerebral cortex is key to understanding the effects of gene dosage on cognitive traits.

## BACKGROUND

Neurodevelopmental and psychiatric conditions, including those studied in the current set of PGC-CNV papers^1,2^, are frequently associated with cognitive deficits^3^. Cognitive ability arises from large-scale brain networks^4–7^, which integrate a diverse range of microscale functions influenced by genetic variations. Recent studies have linked spatial patterns of gene expression in healthy individuals^8^ to structural and functional macroscale brain networks. However, despite significant progress, the role of variations in gene expression across the cerebral cortex in underpinning cognitive ability remains unknown. Specifically, in the absence of large transcriptomic datasets of brain tissue with corresponding phenotypes, a direct association between variation in transcription levels and cognition remains out of reach.

To circumvent this issue, we take advantage of two opportunities. First, over 10%^9,10^ of the general population carries a genomic deletion or duplication fully encompassing one or more coding genes. These deletions and duplications are associated with large decreases and increases ( > 1.5 to 2.5 standard deviations)^11^, respectively, of the expression of genes they encompass^12^. Second, functional genomic resources such as the Allen Human Brain Atlas (AHBA), BrainSpan^13^, and Human Protein Atlas^14,15^ (HPA) provide information on the spatial distribution of gene expression, allowing us to identify brain regions where genes are preferentially expressed.

Previous studies have estimated that a large proportion of the coding genome is associated with lower cognitive abilities when deleted or duplicated^9,10,16,17^. Most of our current knowledge of gene functions involved in cognitive abilities stems from studies investigating ontologies at the cellular, subcellular, and molecular level, i.e., genes associated with intellectual disability are enriched in chromatin remodelling, gene regulation, and neuronal communication processes^18,19^. We postulate that information on spatial profiles of gene expression across the human cerebral cortex will provide novel insights into the effect sizes on cognitive ability of genomic deletions and duplications. To test this hypothesis, we use cerebral-cortex gene-set burden analysis (CC-GSBA) to associate cognitive ability with genomic deletions and duplications that disrupt genes with similar expression profiles across 180 cortical regions. We implemented CC-GSBA in 258,292 individuals from six general population cohorts with measures of cognitive ability. Results demonstrated that most cortical gene-sets were associated with cognitive ability. Deletion and duplication effect sizes showed differential localization along the sensorimotor–association cortical hierarchy, which could not be explained by particular cell types, developmental epochs, or genetic constraint.

### Gene dosage effects on cognitive ability across the cerebral cortex

To understand the relationship between where genes are expressed in cortical networks and their impact on cognitive ability, we partitioned the coding genome into 180 gene-sets preferentially expressed in 180 regions of the human cerebral cortex based on the Glasser atlas^20^. To do so, we used normalized spatial profiles of gene expression from the Allen Human Brain Atlas^21^. A total of 13,678 genes with detectable expression in the cerebral cortex were assigned to cortical regions if their normalized value was higher than 0.5 z-score. As a result, we obtained 180 gene sets corresponding to 180 cortical regions (**Fig. 1**).

**Fig. 1:**
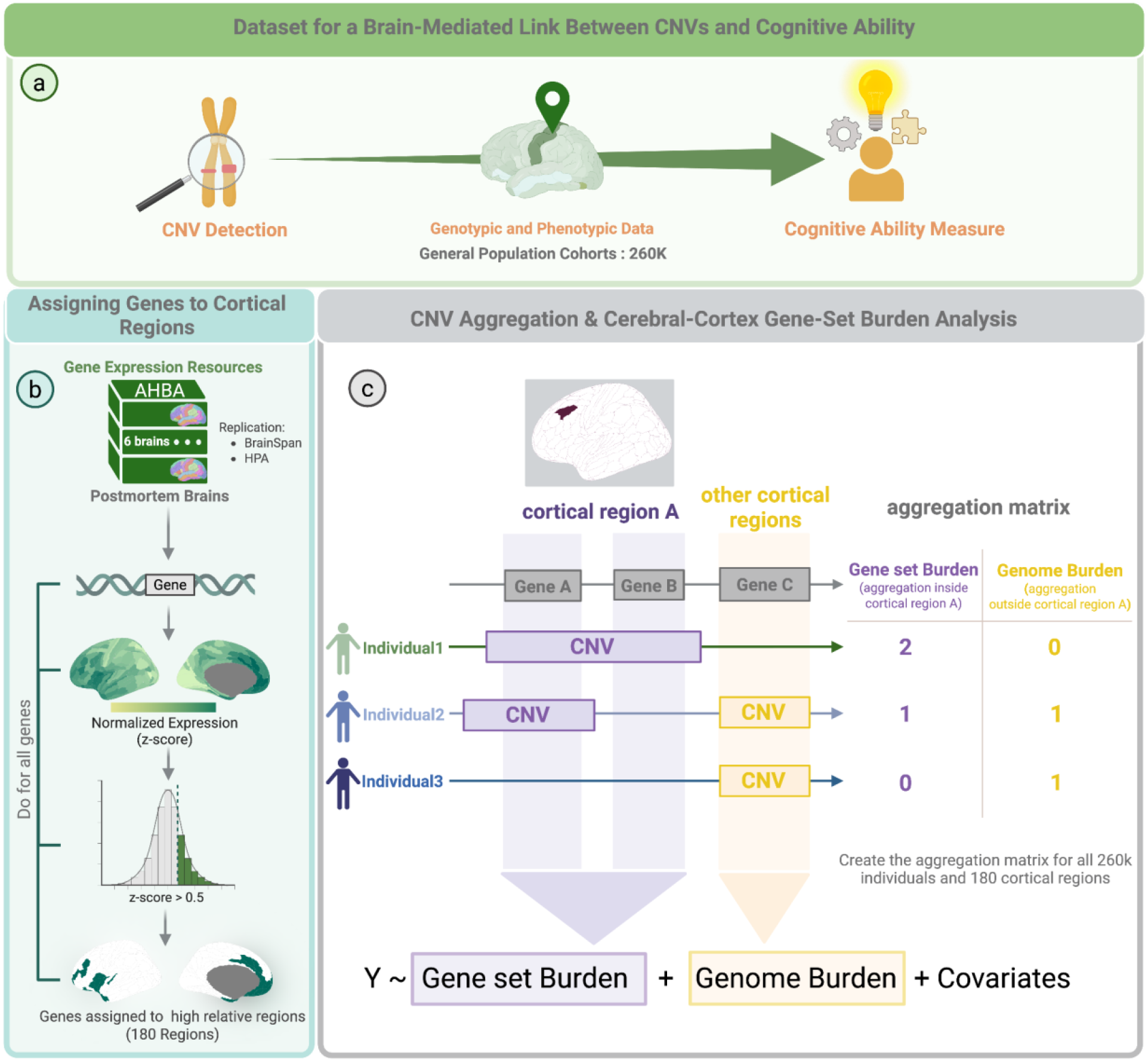
Investigation of the impact of gene expression variation on cognitive ability using Cerebral Cortical Gene Set Burden Analysis (CC-GSBA) **a)** Outline of our methodology for associating CNVs with cognitive ability, leveraging information on where genes are expressed in cortical regions. The analysis integrates genotypic and phenotypic data from six general population cohorts and gene expression data from an independent post-mortem resource, Allen Human Brain Atlas^8^ (AHBA). **b**) Using gene expression data from the AHBA, each gene is initially mapped to cortical regions based on its expression profile and subsequently assigned to regions with preferential expression (Z-score > 0.5). For example, the SST gene shows high expression (Z-score > 0.5) across multiple cortical regions. This process is repeated for all genes (13,678) we have and resulting in 180 gene sets. Replication was assessed using two independent gene expression resources: BrainSpan adult^13^ and Human Protein Atlas^14^ (HPA). **c)** Then, for each individual in the cohort, CNVs are aggregated both inside and outside the regional gene sets. This allows for the calculation of two distinct burden metrics for each cortical region: Gene-Set Burden is the number of protein-coding genes fully encompassed by an individual’s CNVs that belong to the regional gene set, and Genome Burden is the number of remaining protein-coding genes fully encompassed by those CNVs outside the tested gene set. Finally, a regression model is applied to test for a region-specific association between CNV burden and cognitive ability. *Created in BioRender. Kazem, S. (2026)* https://BioRender.com/dxrblrk. CNVs: Copy Number Variants; AHBA: Allen Human Brain Atlas.

We then identified all CNVs > 50 kilobases in 258,292 individuals from six general populations with assessments of general cognitive ability (**Supplementary Table 1**). 13% of individuals (34,169 of 258,292) carried at least one deletion/duplication fully encompassing one or more genes assigned to cortical regions (**Supplementary Fig. 1c; Supplementary Table 2)**. Among the 13,678 genes, 4,405 (32.2%) were deleted and 8,601 (62.9%) duplicated at least once (**Supplementary Fig. 1a, Supplementary Tables 2,3**). Given that deletion/duplication may occur multiple times, we observed a total of 28,108 and 77,695 deleted and duplicated genes, respectively (**Supplementary Fig. 1b, Supplementary Table 3**). We performed a cerebral-cortex gene-set burden analysis (CC-GSBA) by i) aggregating all CNVs affecting gene-sets assigned to cortical regions and ii) conducting 180 linear models to estimate the burden effect on cognitive ability of gene-sets assigned to each of the 180 cortical regions for deletions and duplications separately (**Fig. 1**). Most cortical regions showed (Bonferroni) significant negative associations (**Supplementary Table 4)** with cognitive ability for deletions (n=128) and duplications (n=112, **Fig. 2a**). None of the positive effects were significant. For deletions and duplications, the largest effect sizes were observed in sensorimotor (Cohen’s d = -0.12) and association (Cohen’s d = -0.06) regions, respectively (**Supplementary Tables 4,5**). Regions with the largest significant effects in deletions were those with the smallest (non-significant) effects in duplications and vice versa (**Fig. 2a-b**). The group of regions (41%) significantly associated with both deletions and duplications included those clustered around the dorsolateral prefrontal cortex (DLPFC). These patterns of effect sizes across the cerebral cortex were robust, and correlated between discovery, replication, and full datasets, as well as between the UK Biobank and the other general population cohorts (all two-sided p-spin <0.05; **Supplementary Note 1; Supplementary Fig. 2, Supplementary Table 6**).

**Fig. 2:**
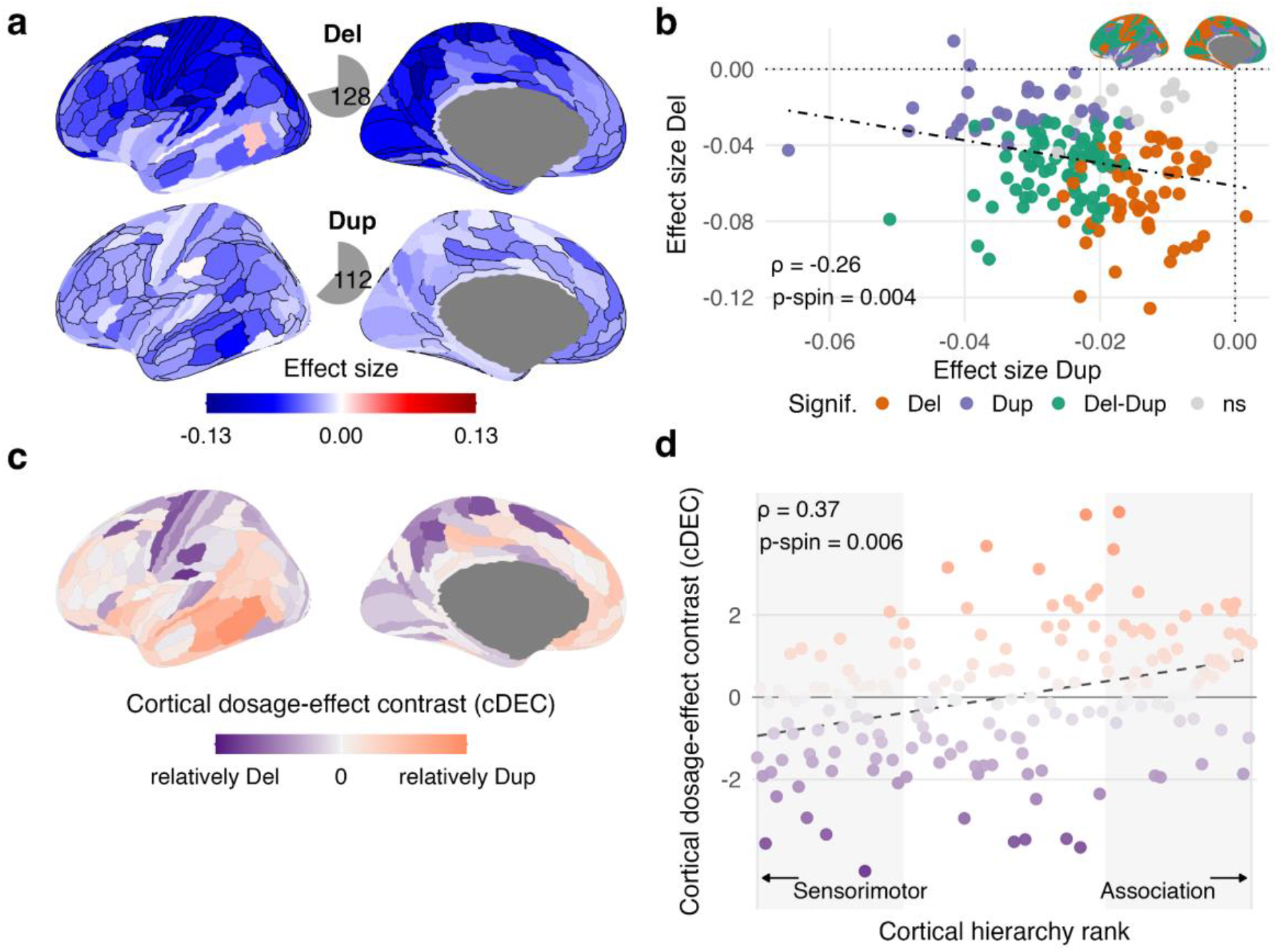
Cortical localization of deletion and duplication effects on cognitive ability. **a)** Cortical maps of deletion and duplication effect sizes across 180 Glasser regions. We performed CC-GSBA for each cortical region, running 180 separate linear models. Effect sizes represent the average effect of genes with high expression in each region on Z-scored cognitive ability, mapped onto the cortex for both deletions and duplications. Regions significant after Bonferroni correction across 360 deletion and duplication tests are outlined in black. Pie charts indicate the number of significant regions (Del, 128; Dup, 112). **b)** Spatial relationship between the complete deletion and duplication effect-size profiles across all 180 cortical regions. Each point represents one region and is colored according to whether the deletion effect, duplication effect or both were significant; non-significant regions are shown in grey. The inset shows their cortical distribution. Selected cortical regions are labelled. Spearman’s ρ and the two-sided spin-permutation p-value are shown. **c)** Cortical dosage-effect contrast (cDEC), defined as the difference between the separately standardized deletion and duplication effect-size profiles, cDEC = z(Del) − z(Dup). Negative values indicate relatively stronger deletion effects, whereas positive values indicate relatively stronger duplication effects. **d)** Association between cDEC and cortical hierarchy rank. Cortical regions were ordered from sensorimotor to association cortex using the T1w/T2w-derived anatomical hierarchy. Shaded regions indicate the bottom and top 30% of the hierarchy (54 regions each), corresponding to the sensorimotor and association poles, respectively. Spearman’s ρ and the two-sided spin-permutation p-value are shown. Spin-permutation tests used 10,000 null rotations and were two-sided. Source data are provided with this paper. Del, deletion; Dup, duplication; cDEC, cortical dosage-effect contrast; p-spin, spin-permutation p-value; ρ, Spearman correlation; Signif., significance.

### Deletion and duplication effects show a cortical dosage-effect contrast

We systematically compared deletion and duplication effect sizes across the complete cortical profiles. Deletion and duplication effect sizes were negatively correlated (all 180 regions: Spearman rho = -0.26; two-sided spin-permutation p-value = 4E-3; selecting regions significant for either CNV type: Spearman rho = -0.40; two-sided spin-permutation p-value = 1E-4), demonstrating that regions with larger deletion effects tend to show smaller duplication effects and vice versa (**Fig. 2b; Supplementary Table 7; Supplementary Note 2)**. Importantly, this spatial relationship reflects differences in the regional magnitude of deletion and duplication effects rather than opposite phenotypic directions, as both CNV classes were predominantly associated with lower cognitive ability. To map this gene dosage response across cortex, we calculated a cortical dosage-effect contrast (cDEC) as the difference between the standardized deletion and duplication effect-size profiles **(Supplementary Note 2)**. Each profile was standardized separately because a raw difference would largely reflect the approximately twofold difference in overall effect magnitude. Negative cDEC values indicate relative deletion weighting, whereas positive values indicate relative duplication weighting. The cDEC profile showed relative deletion weighting toward sensorimotor cortex and relative duplication weighting toward association cortex, with regions showing more similar deletion and duplication effects distributed closer to zero (**Fig. 2c)**. The broad localization of deletion and duplication effects was also observed using two independent gene-expression resources, the Human Protein Atlas and BrainSpan **(Supplementary Figs. 3,4; Supplementary Note 3)**, although their lower spatial resolution limited direct comparison. The patterns were broadly consistent across alternative parcellations (Desikan^22^; and Schaefer^23^) and definitions of cortical gene sets^24^ (**Supplementary Note 3, Supplementary Figs. 5-7; Supplementary Tables 6, 7**).

### Deletion and duplication effects align asymmetrically with the cortical sensorimotor–association hierarchy

We next asked whether the cortical dosage-effect contrast followed the established sensorimotor–association hierarchy, represented by H = −T1w/T2w, increasing toward association cortex (**Methods; Supplementary Fig. 8**). cDEC was positively correlated with H (Spearman ρ = 0.37, two-sided p-spin = 6.4E-3; **Fig. 2d, Supplementary Table 7**), supporting asymmetric alignment of deletion and duplication effects along the hierarchy, with relative deletion weighting toward sensorimotor cortex and duplication weighting toward association cortex.

To clarify how each dosage profile contributed to this alignment, we examined their correlations with H. Because cognitive effects are predominantly negative, a positive correlation indicates larger adverse effects toward the sensorimotor cortex. Deletion effects tended to follow this pattern (ρ = 0.31, p-spin = 0.11), whereas duplication effects were larger toward association cortex (ρ = −0.27, p-spin = 0.008; **Supplementary Table 7**). We then asked whether these hierarchy correlations differed directly. Their difference, the topographic contrast T, was −0.58 (two-sided p-spin = 7.1E-3), supporting the asymmetric alignment captured by cDEC (**Methods; Supplementary Note 2**).

### Phenotypic specificity and cross-cohort generalization

We asked whether the pattern of gene dosage response across the cortex observed above may reflect a general property of CNV burden unrelated to cognitive ability. To do so, we applied CC-GSBA to three groups of UK Biobank phenotypes: cognition-related traits (educational attainment, household income, based on genetic correlations with cognitive ability^25^), blood biochemistry measures (negative control; calcium, LDL and total cholesterol), and birthplace coordinates to assess residual population stratification. Deletion profiles for educational attainment, and household income correlated with the cognitive-ability profile (Spearman ρ = 0.88, and 0.83). However, unlike cognitive ability, the gene dosage response (measured by cDEC) did not follow the cortical hierarchy (**Supplementary Fig. 9; Supplementary Tables 6-8**). Despite approximately twice the sample size, negative controls (Blood biochemistry measures) and birthplace coordinates showed few regional burden associations after multiple-comparisons correction (significant regions out of 180, deletion/duplication: calcium, 2/0; LDL cholesterol, 3/1; total cholesterol, 10/3; birthplace north, 5/0; birthplace east, 0/0). Birthplace profiles did not reproduce the positive cDEC-hierarchy association observed for cognitive ability (**Supplementary Fig. 9; Supplementary Tables 7,8; Supplementary Note 4**).

We then tested cross-cohort and technology generalization using related phenotypes and whole-genome sequencing (WGS) CNVs calls in All of Us^26^. Income profiles were correlated with UK Biobank for deletions and duplications (ρ = 0.54 and 0.44), as were educational-attainment profiles (ρ = 0.64 and 0.26; all two-sided p-spin <0.005; **Supplementary Fig. 10; Supplementary Note 5; Supplementary Tables 6,8**). To assess sensitivity to CNV calling, we repeated the income analysis using GATK-SV whole-genome sequencing CNV calls at ≥50 kb, which also yielded profiles correlated with UK Biobank (ρ = 0.66 for deletions and 0.56 for duplications; both two-sided p-spin <0.005). Including smaller CNVs, 10–40 kb did not change these results (**Supplementary Fig. 11; Supplementary Note 5, Supplementary Table 6**).

### Asymmetric gene dosage response across the cortical organization

To provide biological interpretations of these gene-dosage effect-size maps, we asked whether the gene dosage response across the cortex (cDEC) aligned with known representations of cortical organization. We compared our gene dosage response to 20 normative cortical maps^27^ characterizing the cortical spatial distribution of microstructure, functional-connectivity, energy metabolism, cortical expansion, haemodynamics, allometric scaling and electrophysiology (**Fig. 3**; **Supplementary Fig. 12; Supplementary Table 9**). cDEC was correlated with multiple maps, in particular: microstructural, transcriptomic, electrophysiological, metabolic, and evolutionary expansion; and deletion and duplication profiles showed asymmetric relationships with these maps (**Fig. 3a**):

**Fig. 3:**
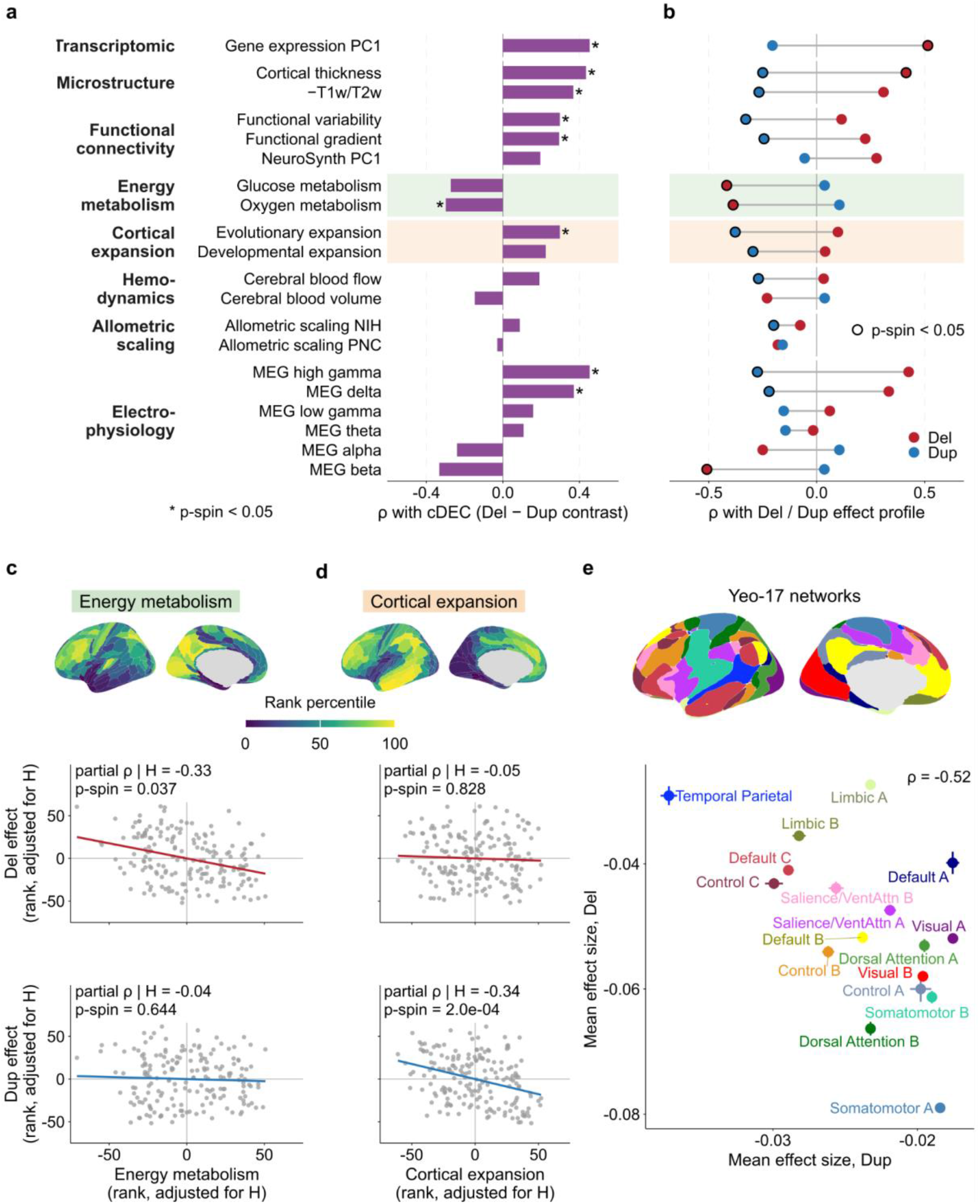
Spatial decoding of deletion and duplication effects and their association with energy metabolism and cortical expansion. **a**) Spatial correlation between the cortical dosage-effect contrast (cDEC) and 20 normative cortical maps^5,27^. The maps are grouped by domain: hierarchy and microstructure, energy metabolism, cortical expansion, hemodynamics, allometric scaling, and magnetoencephalography (MEG). Bars show Spearman’s ρ. Asterisks mark two-sided spin-permutation p < 0.05. Positive ρ indicates that the map is high where duplication effects are relatively stronger than deletion effects. The −T1w/T2w map is the cortical hierarchy H used in Fig. 2. **b**) Spatial Spearman correlation of each map with the deletion (red) and duplication (blue) effect-size profiles. The grey line joining the two points shows their difference, the topographic contrast, T = ρ(Dup,map) - ρ(Del,map). Rows are in the same order as in **a**. Because effect sizes are negative, a negative ρ indicates larger effects where the map is high. A dark outline marks two-sided spin-permutation p < 0.05. Shaded rows mark the two dimensions analysed in **c** and **d**. **c,d)** Deletion (red) and duplication (blue) effects versus energy metabolism (c; oxygen and glucose metabolism) and cortical expansion (d; evolutionary and developmental expansion), adjusted for H. Partial Spearman ρ and two-sided p-spin (composite and H rotated jointly). Unadjusted results in **Supplementary** Fig. 13. **e)** Mean deletion versus duplication effect size across the 17 Yeo resting-state networks (± s.e.m.); colours match the inset network map. Spearman ρ and two-sided p-spin. Spin-permutation tests used 10,000 null rotations and were two-sided. Source data are provided with this paper. Del, deletion; Dup, duplication; cDEC, cortical dosage-effect contrast; p-spin, spin-permutation p-value; ρ, Spearman correlation.

The metabolic maps were preferentially associated with the deletion profile, whereas developmental and evolutionary expansion maps were preferentially associated with the duplication profile (**Fig. 3b-d; Supplementary Fig. 13; Supplementary Note 6; Supplementary Table 9**). This remained true even after adjusting for the previously reported association with cortical hierarchy map (H = -T1w/T2w, **Fig. 3c,d**). This suggests that these different functional maps provide independent sources of information on the effects of gene dosage responses. Overall, this indicates larger adverse effects of deletions and duplications in regions with higher metabolism or expansion, respectively. Resting-state contextualization using 17 functional connectivity networks^28^ showed the largest deletion effects in somatomotor and visual networks and the largest duplication effects in temporoparietal and default-mode networks (**Fig. 3e; Supplementary Note 6)**

### Gene dosage effects on cognitive ability remain similar across levels of regional specificity

We reasoned that genes with the most specific regional expression were those driving the spatial patterns of gene dosage effect sizes across the cortical hierarchy. We therefore compared results from cortical burden tests at four levels of increasing specificity of regional expression. To do so, we thresholded normalized regional expression profiles at four levels of specificity: 0.2, 0.5, 0.8, and 1.0 z-scores (thus creating smaller gene sets with higher regional specificity).

The asymmetric alignment of deletion and duplication effects remained evident across increasing levels of regional specificity. Deletion effects were consistently larger in sensorimotor than association regions, whereas duplication effects were consistently larger in association than sensorimotor regions (two-sided Kolmogorov–Smirnov tests, FDR q<0.05; **Fig. 4a-d, Supplementary Tables 10,11**). Correspondingly, the gene dosage response across the cortex (cDEC) remained associated with the cortical hierarchy across all four regional-specificity thresholds (FDR-adjusted two-sided spin-permutation q<0.05; **Fig. 4e**; **Supplementary Table 7**).

**Fig. 4:**
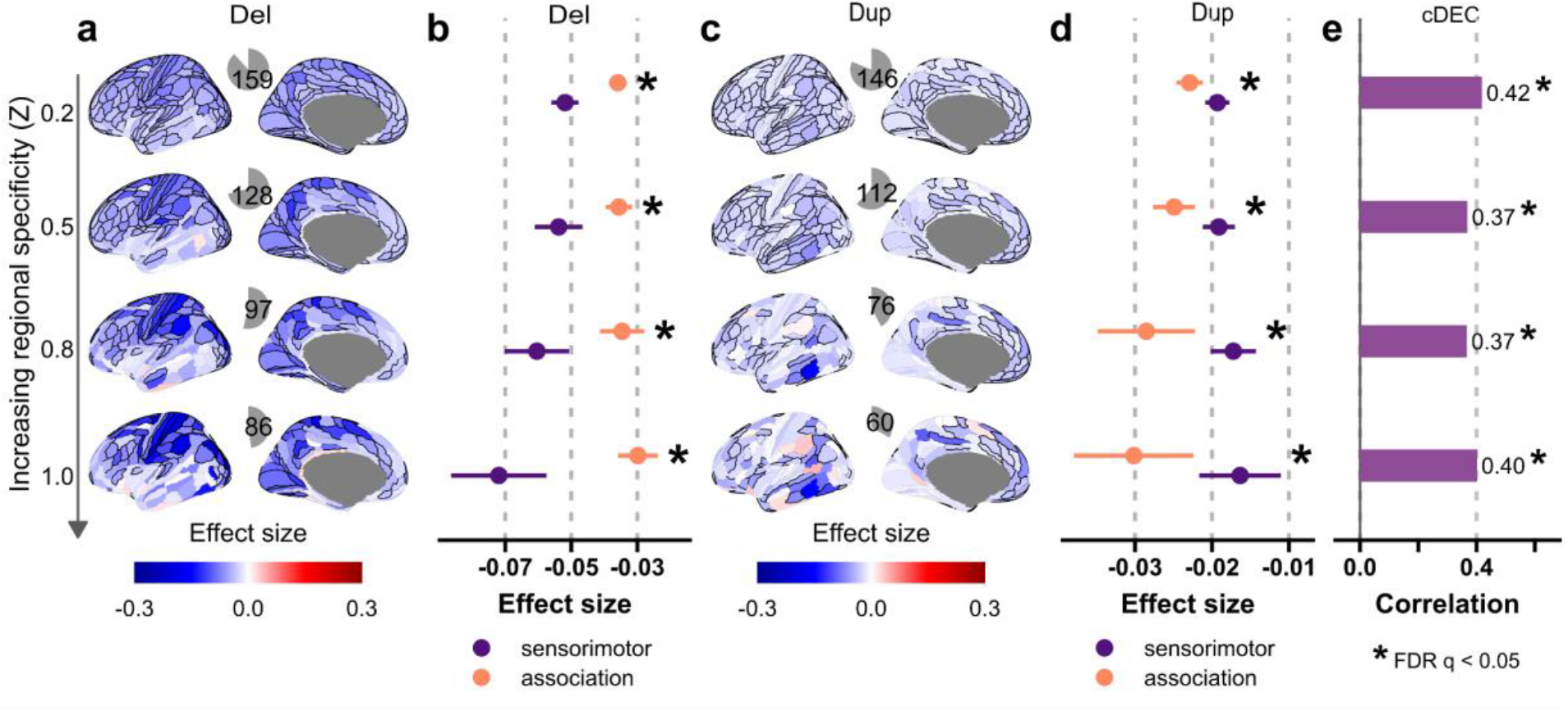
Gene dosage effects on cognitive ability across levels of regional specificity. **a,c**) Cortical effect sizes maps for **a**) deletions and **c**) duplications across four levels of regional specificity (Z = 0.2, 0.5, 0.8 and 1.0). Increasing the threshold progressively restricts each regional gene set to genes with greater regional expression specificity. Regions significant after Bonferroni correction across 180 deletion and 180 duplication regional tests within each threshold are outlined in black. Pie charts indicate the number of significant cortical regions. **b,d**) Mean **b**) deletion and **d**) duplication effect sizes in sensorimotor and association cortical regions across regional-specificity thresholds. Sensorimotor and association regions comprise the bottom and top 30%^75^ of the T1w/T2w-derived cortical anatomical hierarchy^5,27^ (54 regions each). Points show means and horizontal ranges show 95% confidence intervals. Differences between the sensorimotor and association distributions were tested using two-sided Kolmogorov–Smirnov tests; asterisks indicate significance after FDR correction(q<0.05). **e**) Association between the cortical dosage-effect contrast (cDEC) and cortical hierarchy across regional-specificity thresholds. Bars show Spearman’s ρ between cDEC and cortical hierarchy, with correlation coefficients shown beside the bars. Asterisks indicate FDR-adjusted two-sided spin-permutation q<0.05. Spin-permutation tests used 10,000 null rotations. Source data are provided with this paper. Del, deletion; Dup, duplication; cDEC, cortical dosage-effect contrast.

A related analysis showed that sensorimotor–association pole differences and cDEC–hierarchy associations were retained for both brain-elevated and other-tissue-elevated genes, but were weaker for genes with low tissue specificity (**Supplementary Figs. 14, 15**). Additional sensitivity analyses supported broadly consistent cortical patterns across demographic strata (sex, age, and ancestry) and gene-expression stability filters, and after excluding frequently affected genes, recurrent genomic-disorder loci or more frequent CNVs (**Supplementary Figs. 7, 16 and 17; Supplementary Notes 1,3; Supplementary Tables 6, 7, 12, 13**).

### Cortical patterns of gene dosage effect sizes are unrelated to genetic constraint, cell types, and developmental periods

We asked if the spatial distribution of gene dosage effects on cognitive ability was explained by genetic constraint measured by LOEUF^29^ values (loss-of-function observed/expected upper bound fraction). This was not the case. We first showed that genetic constraint was not correlated with the cortical hierarchy (LOEUF, Spearman ρ = 0.08; **Supplementary Fig. 18**). In other words, genes with a spatial pattern of expression following the hierarchy are not under higher constraint. We then stratified genes into 3 groups (**Figs. 5a-b**): intolerant (LOEUF<0.35), mildly intolerant (0.35<LOEUF<1.0), and tolerant (LOEUF>1.0). As expected, increasing intolerance to loss of function was associated with larger effect sizes on cognitive ability (**Figs. 5c-d, Supplementary Tables 3, 5**). However, the regional localization of deletion and duplication effects was retained across constraint strata, although its strength varied between groups (**Fig. 5d, Supplementary Table 7**). The cortical dosage-effect contrast remained significantly associated with the cortical hierarchy in the highly constrained and more tolerant strata, whereas this relationship was weaker in the intermediate LOEUF group (**Fig. 5e, Supplementary Table 7**). Thus, genetic constraint strongly influences the magnitude of gene-dosage effects but does not by itself account for their cortical localization **(Supplementary Note 3)**. Similar results were observed using alternative dosage sensitivity scores^30^ (probability of haploinsufficiency (pHS), probability of triplosensitivity (pTS)) and constraint metrics (Missense Z-score^29^, **Supplementary Figs. 15, 19; Supplementary Table 7**). We investigated additional factors that could explain the gene-dosage cortical patterns described above. First, since bulk tissue expression data combine expression of underlying cell-constituents, we stratified genes based on 6 adult cell-types^14,15^, including astrocytes, oligodendrocyte precursor cells, oligodendrocytes, microglia, excitatory neurons, and inhibitory neurons. Second, the temporal sequence of cortical maturation tends to progress along the sensorimotor-association cortical hierarchy^5,31^. We therefore stratified genes based on their temporal expression trajectories^32^ (prenatal, postnatal, and non-transitional). Overall, the broad cortical organization of deletion and duplication effect sizes remained detectable across these stratifications (**Supplementary Figs. 14,15 and 19, Supplementary Tables 5, 7; Supplementary Note 3**). Sensitivity analysis further showed that the cortical distributions of molecular and sub-cellular biological functions, as detailed by GO-terms^33^, were not associated with the pattern of CNV effects across the cerebral cortex (**Supplementary Fig. 20, Supplementary Tables 14, 15**).

**Fig. 5:**
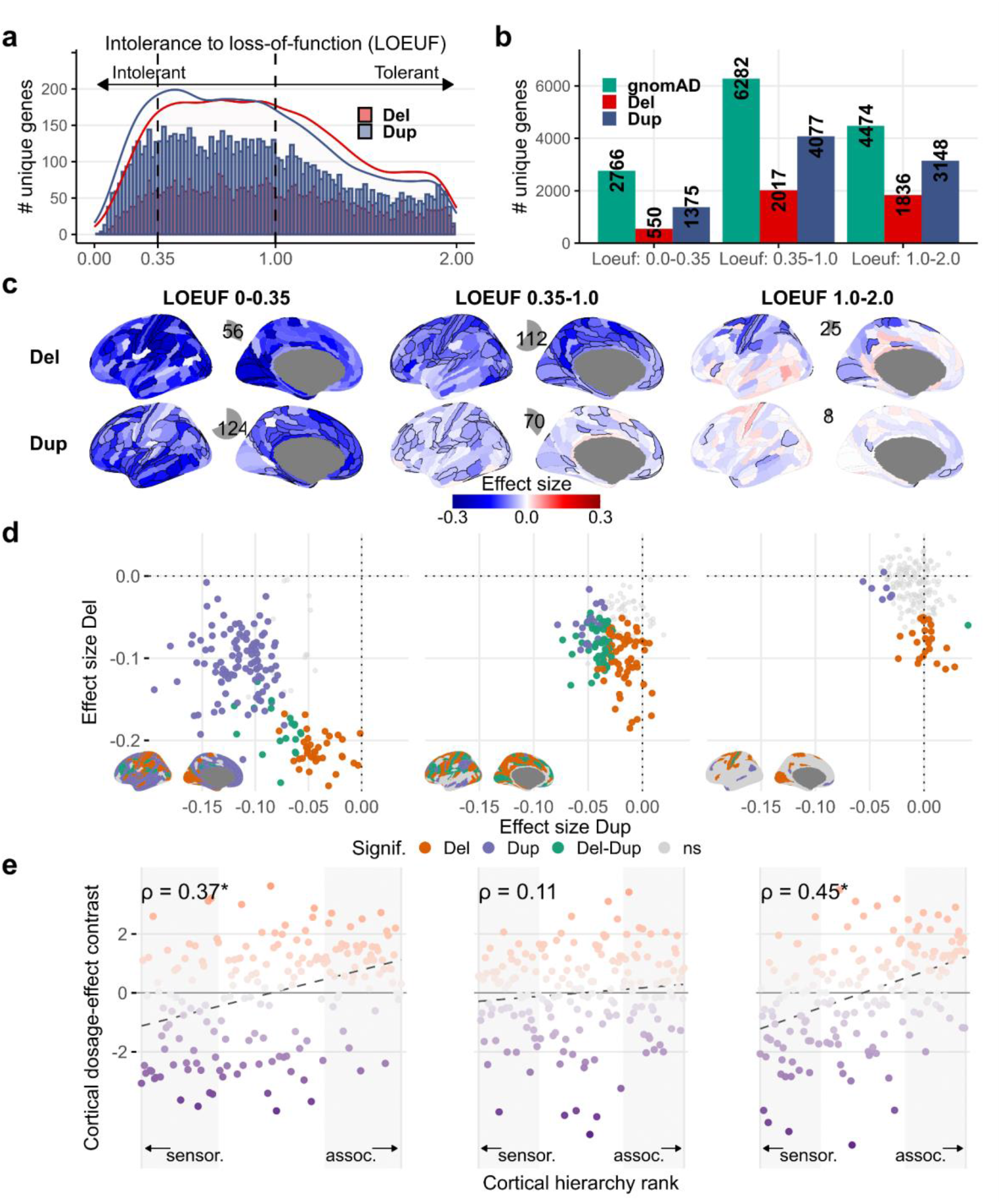
Cortical deletion and duplication effects across levels of genetic constraint. **a**) Distribution of the number of genes deleted or duplicated at least once in our dataset, with respect to their genetic constraint (intolerance to loss of function; LOEUF^29^ values). Histograms show the numbers of unique genes observed in deletions and duplications across LOEUF bins, with density curves indicating their corresponding distributions. Dashed lines denote the boundaries used to define the three LOEUF strata (0-0.35, 0.35-1.0 and 1.0-2.0); lower LOEUF values indicate greater intolerance to loss of function. **b)** Genes stratified into three LOEUF categories: i) highly intolerant (Loeuf1: 0-0.35); ii) intolerant (Loeuf2: 0.35-1.0); and iii) tolerant (Loeuf3: 1.0-2.0). Green bars show the reference number of genes in each LOEUF category, and red and blue bars show the numbers observed in deletions and duplications, respectively. **c**) Cortical deletion and duplication effect-size maps across the three LOEUF strata. Regions significant after Bonferroni correction across 180 cortical regions, two CNV types and three LOEUF strata (1,080 tests) are outlined in black, and pie charts indicate the number of significant cortical regions. A common effect-size scale is used across strata. **d**) Relationship between deletion and duplication effect sizes across all 180 cortical regions within each LOEUF stratum. Regions significant for deletions, duplications or both are colored, whereas non-significant regions are shown in light grey. Insets show the cortical distribution of these significance classes using the same colors. Common x- and y-axis scales are used across strata to show changes in effect-size magnitude with genetic constraint. **e**) Association between the cortical dosage-effect contrast (cDEC) and cortical hierarchy within each LOEUF stratum. Each point represents one cortical region and is colored by cDEC; negative and positive values indicate relative deletion and duplication weighting, respectively. Shaded regions indicate the bottom and top 30% of the cortical hierarchy (54 regions each), corresponding to the sensorimotor and association poles, respectively. Dashed lines indicate linear fits. Spearman’s ρ is shown in each panel; asterisks indicate significant two-sided spin-permutation tests (p<0.05). Spin-permutation tests used 10,000 null rotations. Source data are provided with this paper. Del, deletion; Dup, duplication; cDEC, cortical dosage-effect contrast; LOEUF, loss-of-function observed/expected upper-bound fraction.

## DISCUSSION

This cortical region genetic burden analysis demonstrates that the effect size on cognitive ability of deleted and duplicated genes is organized according to their spatial expression across the cerebral cortex. Genes with preferential expression in sensorimotor regions demonstrated the largest effect on cognition when deleted. At the opposing end of the cortical hierarchy, genes with preferential expression in multimodal association regions had the most significant impact on cognition when duplicated. These two opposing gene dosage cortical patterns could not be explained by particular cell types, developmental epochs, or genetic constraints.

Previous studies have shown that both deletions and duplications are associated with a decrease in cognitive ability^9,34,35^ (**Supplementary Table 3**). We did observe this inverted-U shape for a set of cortical regions (41%) clustered around the dorsolateral prefrontal cortex. However, half (51%) of the cortical regions were either sensitive to duplication or deletions^2,36^, with the largest effects on cognitive ability for deletions being observed in regions that showed smaller or near-zero effects for duplications, and vice versa. This asymmetric pattern suggests that opposing gene dosage alterations may lead to a decrease in cognitive ability through distinct cortical mechanisms.

Results demonstrate that a single cortical region, such as a motor region, can have either the largest or smallest effect on cognitive ability, depending on the class of genetic variant (deletion vs. duplication) disrupting genes preferentially expressed in that region. In studies where multiple classes of genomic variants (leading to different molecular effects) are aggregated at the gene level (i.e., gene-level burden or MAGMA), such variant-specific effect size patterns would not be observed. This may also explain why brain-behaviour neuroimaging association studies have produced small effect sizes. In other words, if heterogeneous molecular mechanisms (with different cortical patterns) are driving inter-individual changes in cognitive ability, BWAS^37^ would therefore only capture the blurred average effect of these factors.

Previous studies have spatially correlated regional gene-expression patterns with MRI-derived spatial distribution of brain alterations associated with genetic disorders and disease^38–41^. These studies are based on the assumptions that two brain maps may result from two modalities, but their spatial pattern may be mechanistically related. We do not make such assumptions as our CC-GSBA approach does not rely on spatial correlations. Instead, we partition gene-dosage effects according to where affected genes are cortically expressed, providing direct effect size estimates across the cerebral cortex. Spatial correlation is used only to characterize the resulting profiles. This approach interrogates the association between a trait and molecular changes in neuroanatomical regions.

Our findings place gene-dosage effects on cognitive ability within the sensorimotor–association hierarchy, a major organizing dimension of the cerebral cortex^5,27^. Sensorimotor and association cortices are characterized by distinct and competing transcriptional programs, with regional gene-expression profiles becoming increasingly aligned with this cortical axis during fetal development^42^. Complementing this observation, opposing molecular hierarchys of primate cortical organization are present at birth and undergo postnatal refinement^43^. Because the genomic deletions and duplications studied here are generally present from conception, it is therefore plausible that gene dosage disruption across development would affect the sensorimotor and association axes and a resulting neurodevelopmental phenotype such as cognitive ability. Consistent with this interpretation, deletion effects were relatively stronger for genes preferentially expressed in sensorimotor regions, which mature earlier, whereas duplication effects were relatively stronger for genes preferentially expressed in association regions characterized by protracted development from late childhood to adulthood^5^. Spatial decoding further distinguished these profiles: deletion effects showed stronger relationships with metabolic cortical maps^44,45^, whereas duplication effects showed stronger relationships with maps of developmental and evolutionary cortical expansion^5,46^.

Our findings highlight primary sensorimotor regions as key contributors to general cognitive ability. We show that in the context of clear haploinsufficiency, these regions show the largest effects on cognitive ability. This aligns with recent research suggesting that the motor network plays a role not only in movement but also in cognitive functions like attention, working memory, and decision-making^47,48^. The observation of overconnectivity in somatomotor networks across psychiatric conditions^49^, cognitive ability, and CNVs is consistent with the fact that many rare loss-of-function variants associated with psychiatric risk are also linked to delayed gross motor milestones^50,51^ and developmental coordination disorders. The companion PGC-CNV study^2^, which analyzes the risk conferred by CNVs for multiple psychiatric conditions, using the same Gene Set Burden Analysis (CC-GSBA) pipeline, also demonstrates the largest effects for deleted genes expressed in sensory motor regions for Schizophrenia, Attention Deficit Hyperactivity Disorder (ADHD), Major Depressive Disorder (MDD), and Autism Spectrum Disorder (ASD). However, the two studies address complementary questions: while the companion study examines gene-set burden associated with psychiatric diagnostic risk, the present study investigates the cortical organization of gene-dosage effects on cognitive ability.

While studies have mapped common variants to genes preferentially expressed in the cerebral cortex^52^ using GWAS summary statistics, as well as run case-control transcriptomic differences^53^, they have primarily focused on the Dorsolateral prefrontal cortex. Our results show that this association is underpinned by complex spatial patterns across the cortex tied to two distinct classes of variants: deletions and duplications. We observed up to a 5-fold difference in effect sizes depending on where the genes are expressed across the cerebral cortex. This underscores the importance of variant-class consideration as well as moving beyond a DLPFC-centric view of transcriptomic studies to investigate the entire cerebral cortex and aligns with a recent transcriptomic study for ASD, albeit across 11 cortical regions^54^.

Finally, we show that the spatial patterns of gene dosage effects on cognitive ability cannot be explained by known contributors such as intolerance to loss of function. We demonstrate that constraint and cortical regional effects on cognition are orthogonal. Similarly, the asymmetric effects across the sensorimotor-association axis were not limited to brain-specific genes, suggesting that this “spatial organizing principle” regroups genes with multiple functions across the entire body. Stratification by cell type also showed the asymmetric effect across six major cell types.

Several limitations in our study are worth noting. The dataset’s large proportion of older adults may affect the interpretation of the results, making it difficult to distinguish between developmental and age-related effects and cognitive decline. The use of a spatial gene expression resource like the AHBA, which has sparse cortical sampling, required spatial interpolation, although each gene was assigned to several contiguous brain regions. However, the patterns of effect sizes were replicated using two independent RNA-seq resources covering the cerebral cortex. CC-GSBA estimates aggregate gene-set burden effects and does not identify individual causal genes or directly measure molecular changes in CNV carriers. However, aggregating gene-dosage events allows us to move beyond known genomic disorders and investigate broader biological functions and cortical organization. Recent proteomic analyses further support the expected molecular dosage effects of deletions and duplications^55^. The predominance of UK Biobank participants and potential residual population structure or participation biases may limit generalizability. However, ancestry and geographic sensitivity analyses, together with cross-cohort comparisons in All of Us using different CNV-calling approaches and size thresholds, supported the broad consistency of the cortical burden profiles. We also acknowledge that while our findings were robust across data subsets and were replicated in independent datasets, the smaller sample size for duplications, in particular, affects the reliability of some estimates.

Overall, this framework highlights the value of mapping genetic association results to the cerebral cortex using RNA-seq resources and concepts from structural, functional, and electrophysiological neuroimaging. Our gene dosage cortical patterns could not be explained by particular cell types, developmental epochs, or genetic constraints, highlighting the fact that the macroscopic network organization of the cerebral cortex is key to understanding the effects of gene dosage on cognitive traits.

## METHODS

### Cohorts and genetic data

#### Ethics

The study protocols were approved by the institutional review board for each of the cohorts involved with recruitment. All procedures contributing to this work comply with the ethical standards of the relevant national and institutional committees on human experimentation and with the Helsinki Declaration of 1975, as revised in 2008. Informed consent and permission to share the data were obtained from all individuals, in compliance with the guidelines specified by the institutional review boards of the recruiting centres. This study was approved by the ethics board of CHU Sainte Justine Research Center, Montreal, Canada.

### Participants and phenotype

The primary analysis included a pooled sample of 258,292 participants from six general-population cohorts: UK Biobank (UKBB^56^, N = 238,176), CARTaGENE^57^ (N = 2,589), Generation Scotland^58^ (N = 13,715), IMAGEN^59^ (N = 1,744), the Lothian Birth Cohort 1936 (LBC^60^; N = 503), and the Saguenay Youth Study (SYS^61^; N = 1,565). UK Biobank therefore contributed 92.2% of the total sample. Overall, 54.6% of participants were female and 95.1% were classified as having European genetic ancestry; age distributions differed substantially across cohorts (**Supplementary Table 1**). The primary analyses used all participants, irrespective of ancestry. To assess robustness to cohort composition, analyses were repeated separately in UK Biobank and in the five other cohorts combined, and in discovery and replication halves from a random split of the pooled sample (Supplementary Fig. 2; Supplementary Methods 1.8). Analyses restricted to participants of European genetic ancestry served as an ancestry sensitivity analysis.

Cognitive ability was harmonized at the level of standardized general cognitive performance rather than by pooling raw scores obtained using different instruments. The primary cognitive phenotype was a general cognitive factor (g-factor) in CARTaGENE and Generation Scotland, WISC-IV in IMAGEN, the Moray House Test in LBC, and WISC-III or g-factor in SYS children and parents, respectively. The g-factor was defined as the first unrotated principal component obtained from principal-component analysis of multiple standardized cognitive measures.

UK Biobank participants could have more than one eligible cognitive assessment. As previously described^9,62^, one cognitive phenotype was selected per participant according to the following hierarchy: in-person g-factor, online g-factor, in-person fluid intelligence, and online fluid intelligence. Each phenotype was standardized separately within its corresponding assessment group before the pooled analysis. As an assessment-mode sensitivity analysis, CC-GSBA was also performed separately using the UK Biobank in-person and online fluid-intelligence subsets. Detailed cohort-specific cognitive assessments, phenotype construction and sample characteristics are provided in the **Supplementary Methods** and **Supplementary Table 1**.

### Sample Processing and Genotyping

UK Biobank DNA samples were genotyped on the UK BiLEVE or UK Biobank Axiom arrays, which have approximately 95% probe overlap; approximately 750,000 common markers were used for CNV calling. The five additional cohorts were genotyped using Illumina SNP arrays: GSA and Omni2.5 in CARTaGENE, 610Kq in Generation Scotland, 610Kq and 660Wq in IMAGEN, 610Kq in LBC, and 610Kq or HOE-12V in SYS. Probe coordinates were harmonized to the hg19 reference genome where required.

Genotype data were processed within a common quality-control framework across cohorts using the same established processing framework^9,62^. For genotype quality control and ancestry estimation, analyses were restricted to autosomal variants with minor-allele frequency >5%, Hardy–Weinberg equilibrium P ≥ 1E-6 and genotype call rate >90%. Duplicate samples, discordance between reported and genetically inferred sex, and relatedness were assessed within each cohort. Genetic ancestry and ancestry principal components were estimated from common autosomal variants using KING and 1000 Genomes reference populations. Platform-specific genotyping details and cohort characteristics are provided in the **Supplementary Methods** and **Supplementary Table 1**.

### CNV calling and quality controls

CNVs were called from SNP-array data using PennCNV^63^ and QuantiSNP^64^ following the previously established pipelines^9,10^. Initial calls required at least three consecutive probes, a minimum size of 1 kb and a caller confidence score ≥15. Calls from the two algorithms were integrated using CNVision, and adjacent CNVs of the same type separated by ≤150 kb were concatenated according to the established pipeline. Array-level quality control required a genotype call rate ≥95%, log R ratio standard deviation <0.35, B-allele frequency standard deviation <0.08 and absolute waviness factor <0.05. The primary analysis was restricted to rare autosomal deletions and duplications ≥50 kb with a confidence score ≥30 for at least one caller and <50% overlap with segmental duplications, the HLA region or centromeric regions. Recurrent CNVs were manually reviewed using log R ratio and B-allele frequency plots by at least one CNV expert. Additional cross-platform and frequency-based filters are described in the **Supplementary Methods**.

### Annotation of CNVs

We annotated the CNVs using Gencode V19 annotation (the reference release for hg19 Human genome release) with ENSEMBL gene name (https://grch37.ensembl.org/index.html). Briefly, we used the bedtools suite (https://bedtools.readthedocs.io/en/latest/) to compare the different elements of the genes that overlap CNVs (UTRs, start and stop codons, exons, and introns). Thus, for each CNV, we obtained its proportion of overlap with each gene as well as with each gene component (UTR, Coding exons, start and stop codons, introns).

### Cortical gene sets

#### Brain parcellations

We used the Glasser brain parcellation^20^ for our main analysis. Glasser parcellation was obtained using multi-modal magnetic resonance images from the Human Connectome Project (HCP)^65^, by delineating 180 areas per hemisphere bounded by sharp changes in cortical architecture, function, connectivity, and/or topography. In addition, we also investigated a functional cortical parcellation: Schaefer^23^, as well as the Desikan^22^ parcellation with 68 parcels.

### Gene expression data

#### Allen Human Brain Atlas (AHBA) data

The AHBA gene data matrix^8^ downloaded from (https://human.brain-map.org/static/download) was further processed using the abagen toolbox (version 0.1.3; https://github.com/rmarkello/abagen)^66^ based on recommendations from previous studies^67,68^. The following text was generated directly from abagen, describing the exact methodologies employed during preprocessing:

First, microarray probes were reannotated using data provided by^67^; probes not matched to a valid Entrez ID were discarded. Next, probes were filtered based on their expression intensity relative to background noise^21^, such that probes with intensity less than the background in >=50% of samples across donors were discarded. When multiple probes indexed the expression of the same gene, we selected and used the probe with the most consistent pattern of regional variation across donors (i.e., differential stability; ^21^. The MNI coordinates of tissue samples were updated to those generated via non-linear registration using the Advanced Normalization Tools (ANTs; https://github.com/chrisfilo/alleninf). Inter-subject variation was addressed by normalizing tissue sample expression values across genes using a robust sigmoid function^69^. Normalized expression values were then rescaled to the unit interval. Finally, gene expression values were normalized across tissue samples using an identical procedure. Samples assigned to the same brain region were averaged separately for each donor and then across donors, yielding a regional expression matrix. After processing, a final 180 cortical regions x 15,633 gene matrix was used for subsequent analysis. We restricted to genes that were detected in the brain (list from the Human Protein Atlas^14^), leading to a final matrix of 180 cortical regions x 13678 genes.

#### BrainSpan-adult data (Independent replication resource)

processed using ABData package in R^70^. Briefly, RNA-seq data were grouped into 5 stages (fetal to adults)^13^, and the mean expression for 11 cortical structures was computed. We used the BrainSpan-adult (stage 5) data as an independent gene expression resource for replication.

#### Human Protein Atlas (HPA) data (Independent replication resource)

RNA-seq data from healthy adults processed using the pipeline proposed in the Human Protein Atlas^14^ (HPA v22). We obtained 73 regions x 16,000 genes matrix of normalized expression, and restricted it to the genes overlapping with our main analysis for subsequent analysis.

### Assigning genes to cortical brain regions

For each gene, normalized expression (abagen output) was Z-scored across the 180 cortical regions (mean 0, s.d. 1). A gene was assigned to a region if its Z-score exceeded 0.5, i.e., its expression in that region was at least 0.5 s.d. above its own cortical mean (Fig. 1; Supplementary Methods 3.1). These genes are hereinafter referred to as preferentially-expressed-genes. On average, a gene was assigned to 52 of 180 regions.

### Stratified cortical gene sets

#### Assigning genes to developmental epochs

To investigate if our findings were dependent on the specific development stage (age) of gene-expression data, we stratified the adult gene-expression data into three non-overlapping gene sets with peak expression in different peak epochs. Briefly, genes were assigned to one of the three developmental epochs, based on BrainVar resource^32^: i) Epoch 1: ranges from 10 to 19 post-conception weeks (pcw); ii) Epoch 2: ranges between 19pcw and 6 postnatal months, and covers the late-fetal transition period, which has been hypothesised to be a critical window for many disorders^32^; and iii) Epoch 3: starts around 6 month and reaches all the way to adulthood.

#### Assigning genes to temporal trajectories

Genes were also stratified based on their temporal trajectories, based on the BrainVar^32^ resource, into: i) falling genes; ii) non-transitional genes; and iii) rising genes.

#### Brain and other tissue-specific gene sets

To investigate if our findings are driven by brain-elevated genes or can be reproduced irrespective of the tissue specificity of gene expression, we used the tissue-specific gene lists from the Human Protein Atlas (HPA, v22)^14^. The HPA performed a normalized transcriptome analysis of protein-coding genes across all major organs and tissue types in the human body, and compared mRNA levels. A gene is considered to have elevated expression based on either of the three conditions: i) Tissue enriched: At least four-fold higher mRNA level in a particular tissue compared to any other tissue; ii) Group enriched: At least four-fold higher average mRNA level in a group of 2-5 tissues compared to any other tissue; and iii) Tissue enhanced: At least four-fold higher mRNA level in a particular tissue compared to the average level in all other tissues. We looked into genes that were brain-elevated (2685 genes), other-tissue elevated, and those that showed low-tissue specificity (https://www.proteinatlas.org/humanproteome/brain)^14^. Of note, we excluded the genes that were not detected in the brain as established by the HPA.

#### Adult cell-type marker gene sets

Since bulk tissue expression data is an ensemble of underlying cell constituents, we looked into deconvolving our findings using cell-type marker gene sets defined in the HPA for major brain cell types^14^. The HPA used single-cell RNA sequencing (scRNAseq) data to cluster genes by their expression across major cell types. In this study, we focused on 6 major cell types in the brain: Astrocytes, Oligodendrocyte precursor cells (OPCs), Oligodendrocytes, Microglial, Excitatory neurons, and Inhibitory neurons. A genome-wide classification of all the protein-coding genes with regard to specificity was performed as follows: (i) cell type enriched genes with at least four fold higher expression levels in one cell type as compared with any other analyzed cell type; (ii) group enriched genes with enriched expression in a small number of cell types (2 to 10); and (iii) cell type enhanced genes with only moderately elevated expression. We considered all genes that met any of these criteria for this study. The HPA confirmed the scRNA-seq profiles and cell type specificity using immunohistochemistry (IHC) on tissue microarrays.

#### Genetic constraint-based gene sets

Previous studies have shown that measures of evolutionary/genetic constraint (such as LOEUF score^29^) explain a significant proportion of the effects of coding CNVs on cognitive and behavioral traits^9,16,17,71^. We then stratified genes into 3 non-overlapping groups based on genetic constraint: i) intolerant (LOEUF<0.35), ii) mildly intolerant (0.35<LOEUF<1.0), and iii) tolerant (LOEUF>1.0), and created the Glasser 180 cortical gene sets for these 3 categories. We applied this same strategy of stratifying into 3 non-overlapping groups for alternative dosage sensitivity scores^30^ (probability of haploinsufficiency (pHS or pHaplo), probability of triplosensitivity (pTS or pTriplo)) and constraint metrics (Missense Z-score^29^).

### Statistical Analysis

#### Cortex-wide copy number variant gene-set burden analysis (CC-GSBA)

We tested the association between cognitive ability and CNV gene burden for each cortical gene set using linear regression. Deletions and duplications were analysed separately. For cortical region *r*, the model was:

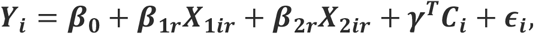

where *Y_i_* is the standardized cognitive score for participant *i*, *X*_1*ir*_ is the number of protein-coding genes encompassed by CNVs that belong to cortical gene set *r* (**Supplementary Fig. 21**), and *X*_2_*_ir_* is the number of remaining protein-coding genes encompassed by CNVs outside that gene set and *C_i_* are covariates (age, sex and genetic ancestry principal components 1–10; **Supplementary Methods**). *β*_1_*_r_*estimates the average effect on cognitive ability of deleting or duplicating one gene preferentially expressed in region *r*, conditional on *X*_2*ir*_. *β*_2*r*_ captures the effect of all other deleted or duplicated genes, including those carried on the same multigenic CNV, so that their contribution is not attributed to the regional gene set (**Supplementary Methods; Supplementary Fig. 22**). Alternative specifications of the Genome Burden term, omitting *X*_2_*_ir_* or replacing it with the total gene count, yielded concordant cortical profiles (**Supplementary Note 7; Supplementary Fig. 23; Supplementary Tables 6, 16)**. CC-GSBA does not introduce a new association test; it applies established gene-set burden regression to spatially defined cortical gene sets.

#### Multiple comparisons correction

Bonferroni correction was applied across the 180 deletion and 180 duplication tests (P < 0.05/360 = 1.39E-4). The same threshold was used for analyses that varied a parameter of the primary analysis (regional-specificity threshold, data subset, gene-expression filter, model specification and phenotype), and was adjusted to the number of regions for other parcellations and expression resources (for example, 0.05/2*34 for Desikan). For analyses that stratified genes into groups (genetic constraint, dosage sensitivity, missense constraint, tissue specificity, cell type, developmental epoch and trajectory), the correction was also applied across strata (P < 0.05/1,080 = 4.63E-5 for three strata; P < 0.05/2,160 = 2.31E-5 for six cell types). Regional significance was used only to identify individual associated gene sets, not to select regions for the cortical profile analyses.

#### Spin permutation testing

Spatial correlations between cortical profiles, and between profiles and reference maps, were computed using Spearman correlation. Their statistical significance was assessed using spin permutation^68,72^ testing, which accounts for spatial autocorrelation between brain regions. Since neighboring cortical regions are expected to be similar, this approach avoids inflated significance values and generates plausible null brain maps. This may account for the overlap between the gene sets for neighboring cortical regions, as the cortical gene set overlap follows the spatial autocorrelation between brain regions. We generated 10,000 spherical rotations for each parcellation. For correlations between a profile and a reference map, the reference map was rotated; for correlations between two profiles, each profile was rotated in turn and the two p-values were averaged; for partial correlations controlling for the hierarchy, the reference map and the hierarchy were rotated jointly. P-spin values were two-sided and are reported without correction, unless stated (**Supplementary Methods**).

#### Correlation of deletion and duplication effect size maps

As a descriptive analysis, we computed Spearman correlation between the complete deletion and duplication effect-size profiles across all cortical regions, irrespective of the statistical significance of individual regional associations. As a sensitivity analysis, we re-computed the correlation after restricting the analysis to cortical regions that were significantly associated with either deletions or duplications after Bonferroni correction. This analysis was performed to assess whether the spatial relationship observed across the complete cortical profiles was dependent on the inclusion of regions without individually significant CC-GSBA associations. Because selection based on statistical significance can alter the observed correlation between the two profiles, this restricted analysis was considered secondary and was not used for the primary inference.

#### Cortical dosage-effect contrast (cDEC)

Spatial analyses used the complete deletion and duplication effect-size profiles across all 180 regions, irrespective of the significance of individual CC-GSBA associations. We first assessed regional heterogeneity: whether variation in effect sizes across cortical regions exceeded that expected from estimation uncertainty. We summarized this descriptively using the dispersion ratio, SD(β)/mean(SE), and I² ^73^. We tentatively used I² <25% to identify profiles with low estimated regional heterogeneity, adapting the conventional interpretation of Higgins et al^73^. These profiles were flagged in summary figures, and their topographic statistics were interpreted cautiously. Because regional estimates share genes and participants, neither measure nor the threshold was treated as a formal test distinguishing regional variation from estimation noise **(Supplementary Methods)**. Similarity between the two profiles was quantified using Spearman correlation.

To characterize their relative cortical localization, we computed a cortical dosage-effect contrast (cDEC):

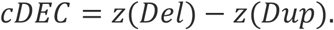

where z(Del) and z(Dup) are the deletion and duplication effect-size profiles, each standardized across all cortical regions using its own mean and standard deviation. Because each profile is standardized separately, cDEC captures their relative cortical localization rather than differences in absolute effect magnitude; negative and positive values indicate relative deletion and duplication weighting, respectively, with respect to the full profiles (**Supplementary Methods**).

#### Localization along the sensorimotor–association hierarchy

We tested whether deletion and duplication effects localize differently along the sensorimotor–association hierarchy (H), defined by the T1w/T2w ratio^5,27,74^). First, the spatial correlation between cDEC and H was tested (cDEC–hierarchy ρ = ρ(cDEC, H)). Second, Spearman correlations were computed between H and Del and Dup cortical profiles (ρ(Del, H) and ρ(Dup H)). Third, their difference was quantified as the topographic contrast:

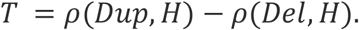

Significance was assessed using two-sided spin-permutation tests (10,000 rotations; **Supplementary Methods**).

#### Sensorimotor and association cortical regions

To quantify the preference for the sensorimotor or association axes, we compared the effect sizes between the most sensorimotor or association regions using a previously established stratification approach^75^ (top 30% and bottom 30% regions of cortical hierarchy, **Supplementary Fig. 8**). We used the anatomical organization hierarchy (T1w/T2w ratio) as the cortical hierarchy.

### Annotation and generalization analyses

#### Spatial decoding using normative cortical maps

We used the previously published cortical organization hierarchies^5^, covering reference maps for molecular, microstructural, electrophysiological, developmental, and functional characterization of the human cortex. Deletion and duplication effect size maps of cognitive ability were annotated using 20 neural maps^5,27^, that included two microstructural (cortical thickness, T1w/T2w), four metabolic (cerebral blood flow, cerebral blood volume, glucose metabolism, oxygen metabolism), three functional (functional gradient, PC1 NeuroSynth, intersubject functional variability), four expansion (evolutionary and developmental expansion, and allometric scaling NIH/PNC), six band-specific electrophysiological signal power (alpha, beta, delta, low gamma, high gamma, theta), and one spatial transcriptomic map (PC1 gene expression). To compute a spatial correlation between deletion/duplication effect sizes maps of cognitive ability and the 20 neural maps, we obtained the 20 neural maps in fsaverage5 space from the neuromaps package and mapped them to 180 cortical regions of Glasser parcellation by averaging values across all the vertices within a parcel. Finally, a spin permutation^27^ method was employed to assess the significance of the spatial correlation. Map acquisition, processing and individual map definitions are provided in the **Supplementary Methods** and **Supplementary Table 9**.

#### Resting state contextualization

We used the BrainStat toolbox (https://brainstat.readthedocs.io/en/master/) for resting state contextualization. We mapped the effect sizes from the 180 cortical regions to 17 resting state brain networks from Yeo 17 parcellation^28^ by first mapping the 180 effect sizes to fsaverage5 vertices, and then averaging the effect sizes for 17 brain networks.

#### Enrichment in gene ontologies across cortical gene sets

The enrichment analyses encompassed a total of 2,519 gene ontology (GO) terms^33^ from three categories: Gene Ontology biological process (GO-BP) with 1938 terms, Gene Ontology cellular component (GO-CC) with 272 terms, and Gene Ontology molecular function (GO-MF) with 309 terms. We used GO-term gene-sets from the Molecular Signatures Database^33^ (MSigDB, the msigdbr package in R, version 7.5.1), with gene-set size >50 and <700. The GO-term analyses encompassed three main steps. Firstly, GO enrichments were computed for each of the 180 brain regions and their corresponding gene sets using the ranked gene set enrichment method (the fgsea^76^ package in R). Secondly, enrichment scores were projected onto cortical maps to generate spatial profiles for each GO-term. Third, these profiles were correlated with the complete deletion and duplication effect-size profiles for cognitive ability across all 180 regions. Spatial correlation analyses were restricted to nine GO terms showing significant enrichment in more than five cortical regions after false discovery rate correction across all 2,519 terms × 180 regions (FDR q<0.05). Correlations were computed using Spearman’s ρ, with significance assessed using two-sided spin-permutation tests with 10,000 rotations. FDR correction was applied across the nine profiles separately for deletions and duplications (q<0.05). Correlations used the signed burden coefficients; a negative correlation therefore indicates greater GO enrichment in regions with larger adverse effects on cognitive ability (Supplementary Tables 14,15).

#### Phenotype-specificity analyses

To assess whether the cortical deletion and duplication profiles observed for cognitive ability were phenotype specific, we applied the same CC-GSBA framework to additional UK Biobank phenotypes. These included educational attainment, household income, three blood biochemistry measures (calcium, LDL and total cholesterol) and birthplace coordinates. For each phenotype, deletion and duplication effects were estimated independently across the same cortical gene sets.

We assessed two complementary relationships: (i) similarity of the same dosage profile across phenotypes (Del–Del and Dup–Dup correlations), and (ii) the relationship between deletion and duplication profiles within each phenotype. Phenotype definitions, coding, sample sizes and processing are provided in the **Supplementary Methods** and **Supplementary Tables 1, 8**.

#### All of Us analyses

We used All of Us^26^ Controlled Tier data release v8 to assess cross-cohort generalization and sensitivity to CNV call set and minimum CNV size. The Array∩WGS resource comprised 402,817 participants and retained CNVs supported by both genotyping arrays and our internally generated WGS calls. Array calls were generated using PennCNV^63^ and QuantiSNP^64^, following a pipeline similar to that used for UK Biobank^36^. Internal WGS calls were generated using gVCF2CNV^77^, which extracts log R ratio and B-allele frequency signals from genomic VCF files for CNV calling with PennCNV alone. We then intersected the retained array and internal WGS call sets to construct Array∩WGS. Separately, we analysed programme-provided GATK-SV structural-variant calls from the same release in 95,405 participants (out of 97,061 samples in All of Us Genomic Quality Report, v8).

Cross-cohort comparisons examined household income and educational attainment using Array∩WGS CNVs ≥50 kb. GATK-SV sensitivity analyses focused on household income and examined minimum CNV sizes of 10, 20, 30, 40 and 50 kb. Deletions and duplications were analysed separately. Cortical profiles were compared across cohorts, call sets and size thresholds using Spearman correlations and sign concordance. Phenotype availability and analysis-specific exclusions further determined the sample used in each analysis. Detailed CNV processing, sample selection and quality-control procedures are provided in **Supplementary Methods 9**.

#### Brain-maps using ggseg

All brain maps shown in this study were generated using the ggseg package^78^ in R.

## SUPPLEMENTARY MATERIALS

Supplementary Figs. 1-23, Supplementary Tables 1-17.

## ADDITIONAL INFORMATION

### MATERIALS & CORRESPONDENCE

Requests for further information and resources should be directed to and will be fulfilled by the lead PI, Sebastien Jacquemont.

### DATA AND MATERIALS AVAILABILITY

The individual-level UK Biobank data analysed in this study are available under restricted access and cannot be shared directly by the authors owing to participant-confidentiality, consent and data-protection requirements governing UK Biobank. Access can be requested by bona fide researchers through the UK Biobank Access Management System (https://www.ukbiobank.ac.uk/enable-your-research/apply-for-access). Applications are reviewed by UK Biobank, and approved researchers obtain access under a Material Transfer Agreement that defines the permitted research use, data-security requirements and duration of access. The data used here were accessed under UK Biobank application 40980.

UK Biobank CNVs were called using the pipeline developed in the Jacquemont Lab (https://github.com/labjacquemont/MIND-GENESPARALLELCNV). The final CNV calls have been returned to UK Biobank as a returned dataset under Return ID 3104 (https://biobank.ndph.ox.ac.uk/ukb/dset.cgi?id=3104). This public catalogue page identifies the returned dataset but does not itself provide direct download access. Researchers who wish to access these CNV calls must have an approved UK Biobank application and request access to Return ID 3104 through the UK Biobank Showcase/Access Management process; once approved, the data can be made available within an approved UK Biobank Research Analysis Platform project.

Individual-level data from the five additional general-population cohorts (CARTaGENE^57^, Generation Scotland^58^, IMAGEN^59^, Lothian Birth Cohort 1936^60^ and Saguenay Youth Study^61^) are available under restricted access through each cohort’s data access procedure; reference papers and data links are listed in **Supplementary Table 1**.

The individual-level All of Us data (Controlled Tier, release v8) are available under restricted access through the All of Us Researcher Workbench (https://www.researchallofus.org).

Access requires researcher registration, completion of required ethics training, institutional authorization as applicable and acceptance of the All of Us Data Use Agreement. Controlled-tier genomic data, including the CNV calls analysed here, are accessible only to approved researchers within the Workbench environment and cannot be exported or shared directly by the authors. Results are reported in accordance with the All of Us dissemination policy.

Publicly available resources used in this study include: gene-expression data from the Allen Human Brain Atlas (https://human.brain-map.org), BrainSpan (https://www.brainspan.org), the Human Protein Atlas, (v22; https://www.proteinatlas.org) and BrainVar; gene-level constraint metrics from gnomAD v2.1 (https://gnomad.broadinstitute.org) and dosage-sensitivity scores (pHaplo, pTriplo); Gene Ontology gene sets from MSigDB (C5 collection; msigdbr 7.5.1); normative cortical maps from neuromaps; the Glasser, Desikan, Schaefer and Yeo cortical parcellations; and PGC-CNV psychiatric gene-set burden summary statistics². The cortical gene sets, gene-level scores and strata, Gene Ontology enrichment results and the CC-GSBA pipeline with a simulated toy example are available on Zenodo (https://doi.org/10.5281/zenodo.23776818). All summary statistics generated in this study are provided in Supplementary Tables 1–17. Source data are provided with this paper.

### CODE AVAILABILITY

The code for generating all the Figures reported in the main analysis and supplement material can be found at the following GitHub link: https://github.com/kkumar-iitkgp/DelDup_Asymmetry_Cortex. Additional data (gene-sets) and code are archived on Zenodo: https://doi.org/10.5281/zenodo.23776818.

### AUTHOR CONTRIBUTIONS

K.K., S.K., G.Du. and S.J. designed the study. K.K. and S.K. performed the analyses and drafted the manuscript. G.Du. and S.J. jointly supervised the work. J.Y., F.B., B.C. and M.S.D. performed the All of Us CNV calling, quality control and analyses. All other authors contributed cohort data, CNV calling or interpretation of the results. All authors reviewed and edited the manuscript.

## ACKNOWLEDGEMENTS

This research was supported by Calcul Quebec (http://www.calculquebec.ca) and Compute Canada (http://www.computecanada.ca), the Brain Canada Multi-Investigator initiative, NIH U01 grant for CAMP (1U01MH119690-01), the Canadian Institutes of Health Research, CIHR_400528, The Institute of Data Valorization (IVADO) through the Canada First Research Excellence Fund, Healthy Brains for Healthy Lives through the Canada First Research Excellence Fund. Dr Jacquemont is a recipient of a Canada Research Chair in neurodevelopmental disorders and a chair from the Jeanne et Jean Louis Levesque Foundation. The Canadian Institutes of Health Research and the Heart and Stroke Foundation of Canada fund the Saguenay Youth Study (SYS). SYS was funded by the Canadian Institutes of Health Research (T.P., Z.P.) and the Heart and Stroke Foundation of Canada (Z.P.). Funding for the project was provided by the Wellcome Trust. This work was also supported by NIH award U01 MH119690 granted to L.A., S.J., and D.C.G. and U01 MH119739. LBC1936 is supported by the Biotechnology and Biological Sciences Research Council and the Economic and Social Research Council (BB/W008793/1; which supports S.E.H.), Age UK (Disconnected Mind project), the Milton Damerel Trust, and the University of Edinburgh. Generation Scotland received core support from the Chief Scientist Office of the Scottish Government Health Directorates (CZD/16/6) and the Scottish Funding Council (HR03006) and is currently supported by the Wellcome Trust (216767/Z/19/Z). Genotyping of the GS:SFHS samples was carried out by the Genetics Core Laboratory at the Edinburgh Clinical Research Facility, University of Edinburgh, Scotland, and was funded by the Medical Research Council UK and the Wellcome Trust (Wellcome Trust Strategic Award “Stratifying Resilience and Depression Longitudinally” (STRADL) (reference 104036/Z/14/Z). IMAGEN received support from the following sources: the European Union-funded FP6 Integrated Project IMAGEN (Reinforcement-related behaviour in normal brain function and psychopathology) (LSHM-CT-2007-037286); the Horizon 2020-funded ERC Advanced Grant “STRATIFY” (Brain network based stratification of reinforcement-related disorders) (695313); the Human Brain Project (HBP SGA 2, 785907, and HBP SGA 3, 945539); the Medical Research Council Grant “c-VEDA” (Consortium on Vulnerability to Externalizing Disorders and Addictions) (MR/N000390/1); the NIH (R01DA049238, A decentralized macro and micro gene-by-environment interaction analysis of substance use behavior and its brain biomarkers); the National Institute for Health Research (NIHR) Biomedical Research Centre at South London and Maudsley NHS Foundation Trust and King’s College London; the Bundesministerium für Bildung und Forschung (BMBF grants 01GS08152 and 01EV0711; Forschungsnetz AERIAL 01EE1406A and 01EE1406B; Forschungsnetz IMAC-Mind 01GL1745B); the Deutsche Forschungsgemeinschaft (DFG grants SM 80/7-2, SFB 940, TRR 265, and NE 1383/14-1); the Medical Research Foundation and Medical Research Council (grants MR/R00465X/1 and MR/S020306/1); and the NIH-funded ENIGMA (grants 5U54EB020403-05 and 1R56AG058854-01). The funder had no role in the design and conduct of the study or preparation of the manuscript. We thank the participants for their time and contributions to these databases.

