## Supplementary Material for "Genomic deletion and duplication effects on cognitive ability lie at the opposing ends of the cortical sensorimotor–association hierarchy"

**Supplementary Information**

**Supplementary Table legend**

- **Supplementary Table 1** Data demographics
- **Supplementary Table 2** Summary statistics on gene count and number of CNV carriers
- **Supplementary Table 3** Overall burden analysis results
- **Supplementary Table 4** Cortical gene set burden analysis results
- **Supplementary Table 5** Summary statistics on cortical gene set effect sizes
- **Supplementary Table 6** Deletion and duplication profile stability: concordance (Spearman ρ) between effect-size profiles (del–del and dup–dup)
- **Supplementary Table 7** Profile statistics for every analysis: dispersion (SD/SE), $I^2$, cDEC–hierarchy ρ, correlations of the deletion and duplication profiles with the hierarchy (H), and deletion–duplication correlation
- **Supplementary Table 8** Cortical gene set burden analysis results for other phenotypes in UK Biobank and for income and educational attainment in All of Us
- **Supplementary Table 9** Correlations between deletion and duplication effect-size profiles and 20 neuromaps
- **Supplementary Table 10** Fold change between regional specificity thresholds of Z>0.2 and Z>1.0 for sensorimotor and association cortical axes
- **Supplementary Table 11** Summary statistics on effect sizes for sensorimotor and association cortical axes
- **Supplementary Table 12** Cortical gene set burden analysis results for CNV-level robustness analyses
- **Supplementary Table 13** Genomic disorder catalogue: 41 ClinGen recurrent CNV

regions and the 16 major loci

- **Supplementary Table 14** GO-term enrichments in Glasser 180 cortical gene sets
- **Supplementary Table 15** Correlations between GO-term enrichment profiles and deletion/duplication effect-size profiles
- **Supplementary Table 16** Cortical gene set burden analysis results under three Genome Burden specifications (G0, G1, G3)
- **Supplementary Table 17** Harmonized All of Us GATK-SV call-set counts at minimum CNV sizes of 10, 20, 30, 40 and 50 kb, separately for deletions and duplications

**Supplementary Methods**

**1. Study cohorts and cognitive phenotypes**

The primary analysis included 258,292 participants from six general-population cohorts: UK Biobank (UKBB[1], N = 238,176), CARTaGENE[2] (N = 2,589), Generation Scotland[3] (N = 13,715), IMAGEN[4] (N = 1,744), the Lothian Birth Cohort 1936 (LBC[5]; N = 503), and the Saguenay Youth Study (SYS[6]; N = 1,565). This is the same harmonized cognitive dataset described in our previous studies[7,8]. The pooled sample was used for the primary analyses in the present study. UK Biobank and the five non-UKBB cohorts combined were additionally analysed separately to assess robustness to cohort composition.

The primary analysis retained participants across genetically inferred ancestry groups. Of the total sample, 245,562 participants (95.1%) were classified as having European genetic ancestry and 12,730 (4.9%) as other ancestries. The sample included 141,142 females (54.6%) and 117,150 males (45.4%). Because age distributions differed markedly across cohorts, and SYS included both adolescents and their parents, age is reported separately by cohort and SYS generation in **Supplementary Table 1** rather than summarized primarily by a pooled mean. Analyses restricted to participants of European genetic ancestry were performed as a sensitivity analysis.

Cognitive measures differed across cohorts. We therefore harmonized cognition at the level of standardized general cognitive performance rather than pooling raw test scores across instruments. Each phenotype was standardized within its corresponding cohort or assessment group. Where a g-factor was used, cognitive measures were standardized before principal-component analysis, and the first unrotated principal component was retained as the measure of general cognitive ability.

**1.1 UK Biobank**

After phenotypic and genotypic quality control, 238,176 UK Biobank[1] participants contributed to the pooled cognitive analysis. Some participants had more than one eligible cognitive assessment. For these individuals, one phenotype was selected according to the following priority order: (1) in-person g-factor (N = 73,882), (2) online g-factor (N = 62,080), (3) in-person fluid intelligence (N = 88,441), and (4) online fluid intelligence (N = 13,773).

The g-factor was computed separately for the in-person and online assessment groups using cognitive tasks assessing executive function, processing speed, verbal declarative memory and crystallized ability, including trail-making, symbol-digit substitution, paired-associate learning and picture vocabulary. The first unrotated principal component was retained and standardized separately within each assessment group.

Fluid intelligence was assessed either at an assessment-centre visit (UK Biobank field 20016) or through the online assessment (field 20191). The test comprised 13 verbal and numerical reasoning items, with participants completing as many items as possible within two minutes. Because score distributions differed between assessment modes, fluid-intelligence scores were standardized separately within the in-person and online groups using their respective distributions (in-person mean = 6.07, s.d. = 2.15; online mean = 6.61, s.d. = 1.98).

The final UK Biobank sample comprised 129,725 females and 108,451 males, with a mean age of 59.03 years (s.d. = 8.59). A total of 225,733 participants were classified as having European genetic ancestry and 12,443 as other ancestries.

**1.2 CARTaGENE**

The CARTaGENE analysis included 2,589 participants after genotype and phenotype quality control, comprising 2,074 participants genotyped using the GSA platform and 515 using Omni2.5. General cognitive ability was represented by a g-factor derived from verbal and numerical reasoning, paired-associate learning, and two-choice reaction time. The first unrotated principal component was retained and standardized within the cohort before inclusion in the pooled analysis.

The sample comprised 1,375 females and 1,214 males, with a mean age of 53.94 years (s.d. = 7.84). Of these participants, 2,472 were classified as having European genetic ancestry and 117 as other ancestries.

**1.3 Generation Scotland**

The Generation Scotland analysis included 13,715 participants genotyped using the 610Kq array. General cognitive ability was represented by a g-factor derived from four measures: Logical Memory, Digit Symbol, verbal fluency and Mill Hill Vocabulary, capturing verbal declarative memory, processing speed, executive function and vocabulary. The first unrotated principal component was retained and standardized within the cohort.

The sample comprised 8,081 females and 5,634 males, with a mean age of 46.73 years (s.d. = 15.00). A total of 13,672 participants were classified as having European genetic ancestry and 43 as other ancestries.

**1.4 IMAGEN**

The IMAGEN analysis included 1,744 adolescents genotyped using the 610Kq or 660Wq arrays. Cognitive ability was assessed using the Wechsler Intelligence Scale for Children, Fourth Edition (WISC-IV), and the cognitive score was standardized within the cohort before inclusion in the pooled analysis.

The sample comprised 891 females and 853 males, with a mean age of 14.45 years (s.d. = 0.37). A total of 1,624 participants were classified as having European genetic ancestry and 120 as other ancestries.

#### 1.5 Lothian Birth Cohort 1936

The LBC analysis included 503 participants genotyped using the 610Kq array. Cognitive ability was represented by the Moray House Test obtained at the later-life assessment, when participants were approximately 70 years of age. The cognitive score was standardized within the cohort before analysis.

The sample comprised 246 females and 257 males, with a mean age of approximately 70.0 years (s.d. = 0.83). A total of 500 participants were classified as having European genetic ancestry and three as having other ancestries.

#### 1.6 Saguenay Youth Study - children

The SYS child sample included 967 adolescents. Cognitive ability was assessed using the Wechsler Intelligence Scale for Children, Third Edition (WISC-III), and standardized within the analysis group. Participants were genotyped using either the 610Kq (N = 559) or HOE-12V (N = 408) array.

The sample comprised 505 females and 462 males, with a mean age of 14.99 years (s.d. = 1.84). A total of 965 participants were classified as having European genetic ancestry and two as other ancestries.

#### 1.7 Saguenay Youth Study - parents

The SYS parent sample included 598 participants genotyped using the HOE-12V array. General cognitive ability was represented by a g-factor derived from 12 tasks administered through the Cambridge Brain Sciences platform: colour-word remapping, spatial planning, self-ordered search, paired-associate learning, digit span, spatial span, visuospatial working memory, interlocking polygons, feature matching, odd-one-out, grammatical reasoning and spatial rotation. The first unrotated principal component was retained and standardized within the parental sample.

The sample comprised 319 females and 279 males, with a mean age of 49.50 years (s.d. = 4.87). A total of 596 participants were classified as having European genetic ancestry and two as other ancestries.

**1.8 Cohort and assessment sensitivity analyses**

The pooled six-cohort dataset was used for the primary analyses. To test whether the cortical profiles depended on the predominance of UK Biobank, CC-GSBA was repeated separately in UK Biobank and in the five other cohorts combined. These two non-overlapping datasets were compared with each other and with the pooled analysis. To assess internal replicability, the pooled sample was randomly split into two non-overlapping halves (discovery and replication). CC-GSBA was performed in each half, and the two profiles were compared with each other and with the pooled analysis. Profiles were compared separately for deletions and duplications using Spearman correlation across the 180 regions, with spin-permutation testing (Supplementary Fig. 2; Supplementary Table 6).

To assess sensitivity to cognitive assessment mode, CC-GSBA was also performed separately among UK Biobank participants assessed for fluid intelligence at an assessment-centre visit and those assessed using the online fluid-intelligence test. The two phenotypes were standardized independently using their assessment-specific distributions.

To assess sensitivity to genetic ancestry, analyses were repeated after restriction to participants classified as having European genetic ancestry. The all-ancestry pooled analysis remained the primary analysis.

### 2. Genotyping, CNV calling and quality control

#### 2.1 Genotyping and genetic ancestry

DNA was extracted from blood. UK Biobank participants were genotyped using the UK BiLEVE or UK Biobank Axiom arrays. These arrays have approximately 95% probe overlap, and approximately 750,000 common markers were used for CNV detection. The non-UKBB cohorts were genotyped using Illumina SNP arrays: GSA and Omni2.5 in CARTaGENE, 610Kq in Generation Scotland, 610Kq and 660Wq in IMAGEN, 610Kq in LBC1936, and 610Kq or HOE-12V in SYS. Where required, probe coordinates were converted to the hg19 reference genome before harmonization across datasets.

Genotype processing and quality control were performed using PLINK v1.9. For ancestry estimation and genotype quality control, analyses were restricted to autosomal markers with minor-allele frequency >5%, Hardy–Weinberg equilibrium P ≥ 1E−6 and genotype call rate >90%. Duplicate samples, relatedness and discordance between reported and genetically inferred sex were evaluated within each cohort.

Genetic ancestry and ancestry principal components were derived from common autosomal variants using KING and 1000 Genomes reference populations following the previously established procedure[7,8]. The primary analysis retained participants across ancestry groups.

Analyses restricted to participants classified as having European genetic ancestry were used as sensitivity analyses for population structure.

**2.2 CNV calling**

CNVs were called independently from SNP-array intensity data using PennCNV and QuantiSNP. For both algorithms, initial CNV detection required at least three consecutive probes, an event size ≥1 kb, and a caller confidence or likelihood score ≥15. Calls from the two algorithms were integrated using CNVision. Events with copy number <2 were classified as deletions and events with copy number >2 as duplications.

Following integration of the caller outputs, adjacent CNVs of the same dosage class were concatenated when separated by ≤150 kb, provided that the events satisfied the minimum size and probe-number criteria.

**2.3 Array-level quality control**

Arrays were required to have a genotype call rate ≥95%, log-R-ratio standard deviation <0.35, B-allele-frequency standard deviation <0.08, and absolute waviness factor <0.05. Samples showing unusually large numbers of CNV calls were excluded, using thresholds of 50 calls for arrays with fewer than one million probes and 200 calls for arrays with at least one million probes. Duplicate samples and samples with discordance between reported and genetically inferred sex were also excluded.

**2.4 CNV-level filtering and cross-platform harmonization**

Analyses were restricted to autosomal CNVs. Retained events were required to have a confidence score ≥30 for at least one caller, a size ≥50 kb, an unambiguous deletion or duplication classification, and <50% overlap with segmental duplications, the HLA region or centromeric regions. CNVs ≥10 Mb were excluded.

Because CNV detection sensitivity varies with array design and probe density, additional filters were applied before pooling cohorts. First, a CNV was required to be represented by at least ten probes on each array technology included in the pooled analysis. Second, CNVs occurring at a frequency ≥1% in any cohort were removed from all cohorts. Third, events showing extreme between-cohort variation in detection frequency were excluded using the upper 1% of the corresponding coefficient-of-variation distribution. For these frequency-based filters, CNVs were considered equivalent when they shared ≥50% reciprocal genomic overlap.

Recurrent CNVs were manually reviewed using log-R-ratio and B-allele-frequency intensity profiles. Residual likely artefacts were additionally filtered using the previously validated framework[7,8], which combines array-quality, genomic-context and call-level characteristics to distinguish high-confidence CNVs from technical artefacts.

#### 2.5 CNV annotation

CNVs were annotated on the hg19 reference genome using GENCODE v19 protein-coding gene annotations. CNV coordinates were intersected with gene coordinates using bedtools. For gene-burden analyses, a protein-coding gene contributed to the burden score when it was fully encompassed by the CNV. Deletions and duplications were annotated and analysed separately.

### 3. Construction of cortical gene sets

#### 3.1 Allen Human Brain Atlas

The primary cortical gene sets were derived from the Allen Human Brain Atlas (AHBA). AHBA microarray data were processed using the abagen toolbox (version 0.1.3) following established recommendations for imaging-transcriptomic analyses. Probes were reannotated, probes with low expression relative to background were removed, and expression samples were assigned to cortical regions. The primary analysis used the Glasser parcellation, with 180 cortical regions represented in the analysis.

For each protein-coding gene, regional expression was standardized across the 180 cortical regions (Z-score: the gene's regional expression minus its mean across regions, divided by its standard deviation across regions). Genes with relatively elevated expression in a region were assigned to that regional gene set. The primary definition used a regional expression threshold of $Z > 0.5$. A gene could therefore belong to more than one cortical gene set when it showed elevated expression in multiple regions. Sensitivity analyses used alternative regional-specificity thresholds of $Z > 0.2$, 0.8 and 1.0.

#### 3.2 Alternative parcellations and transcriptomic resources

To assess dependence on the cortical parcellation and expression resource, we repeated the regional analyses using alternative cortical definitions, including the Schaefer parcellation[9], Desikan parcellation[10,11], and independently processed transcriptomic resources. For, alternative gene-set definition, we used the 16 spatial modules[12] of cortical gene expression identified from Allen Human Brain atlas data. Adult BrainSpan RNA-seq[13–15] data were used as an independent lower-resolution cortical expression resource, covering 11 cortical regions. Human Protein Atlas[16] (HPA v22) RNA-seq data from healthy adults provided an additional anatomical expression resource spanning 73 regions. Because these resources differ substantially in spatial resolution and anatomical definition from the Glasser atlas[17], they were used to test broad regional localization rather than as exact parcel-wise replications of the primary analysis.

**4. Cortex-wide copy number variant gene-set burden analysis (CC-GSBA)**

The CC-GSBA model is specified in the Methods. This section describes how its terms were constructed, the rationale for conditioning on the remaining CNV burden, and model-specific sensitivity analyses. The outcome is the within-cohort standardized cognitive score described in **Supplementary Methods 1**.

**4.1 Burden variables**

Both burden variables count protein-coding genes fully encompassed by a participant's CNVs (**Supplementary Methods 2.5**).

- $X_{1ir}$ counts those genes that belong to cortical gene set r (**Supplementary Methods 3.1**).
- $X_{2ir}$ counts all other encompassed protein-coding genes, including genes assigned to other cortical gene sets and genes not assigned to any region.

For every region, $X_{1ir}$ + $X_{2ir}$ equals the participant's total number of deleted (or duplicated) protein-coding genes. Only the split between the two terms changes from region to region. Because a gene can belong to several overlapping regional gene sets, the same CNV can contribute to $X_1$ in several regional models.

The burden data are sparse: approximately 13% of participants carry a deletion or duplication fully encompassing at least one protein-coding gene, so $X_{1ir}$ = 0 for most participants. Among carriers, $X_{1ir}$ is a low count with a Poisson-like, right-skewed distribution: most carriers have one or a few regional genes, and a small number of large multigenic CNVs contribute higher values. The information for estimating $\beta_{1r}$ therefore comes mainly from carriers. All participants, including non-carriers ($X_{1ir} = X_{2ir} = 0$), were retained in every model so that covariate effects are estimated in the full sample.

**4.2 Covariates**

$C_i$ included age, sex and genetic ancestry PCs 1–10. Age was defined as age at the cognitive assessment and sex as self-reported sex, both taken from each cohort's phenotype data; samples whose reported sex did not match their genetically inferred sex were excluded (**Supplementary Methods 2.1 and 2.3**). Ancestry principal components were derived as described in **Supplementary Methods 2.1**.

**4.3 Rationale for conditioning on $X_2$**

The main reason for including $X_2$ is the multigenic structure of CNVs: a single CNV can encompass genes in a regional cortical gene set together with many genes outside it. Without $X_2$, the full effect of such a CNV would be attributed to the subset of its genes in the regional set, inflating $\beta_{1r}$. $X_2$ also adjusts for differences in overall CNV load between individuals.

Because many genes affect cognitive ability, conditioning on $X_2$ also absorbs some true effects of genes outside the regional set. $\beta_{1r}$ is therefore a conservative estimate of the additional per-gene effect of genes preferentially expressed in region r.

**4.4 Estimation and interpretation**

Models were fitted by ordinary least squares, separately for each of the 180 Glasser regions and for deletions and duplications. Because cognitive ability is standardized (mean 0, SD 1), $\beta_{1r}$ is the average change in cognitive ability, in standard deviations, for each additional gene of gene set r encompassed by a deletion (or duplication). For example, $\beta_{1r} = -0.10$ means that each additional deleted gene preferentially expressed in region r is associated with a 0.10 SD lower cognitive score, holding the remaining CNV burden and covariates constant.

The robustness analyses in **Supplementary Methods 3** (alternative parcellations, regional-specificity thresholds and expression resources) used the same model, covariates and CNV data. Only the definition of the regional gene sets, and therefore the split of each participant's genes between $X_1$ and $X_2$, changed.

#### 4.5 Sensitivity to conditioning on $X_2$

To assess the effect of conditioning on the remaining CNV burden, all regional models were re-fitted without $X_2$, keeping the same outcome, covariates and gene sets. The resulting deletion and duplication cortical profiles were compared with the primary profiles using Spearman correlation across the 180 regions, and the cortical profile analyses (deletion–duplication correlation, cDEC and hierarchy contrast; **Supplementary Methods 5**) were repeated on these profiles. Sensitivity analyses for cohort composition, assessment mode and genetic ancestry are described in **Supplementary Methods 1.8**.

### 5. Cortical profile and spatial analyses

#### 5.1 Complete cortical profiles

Analyses used the complete regional effect-size profiles, irrespective of the significance of individual burden associations. Before characterizing their spatial organization, we assessed regional heterogeneity, defined here as variation in regional effect sizes beyond estimation uncertainty. We calculated two descriptive summaries separately for deletions and duplications.

The dispersion ratio was the standard deviation of regional effect sizes divided by their mean standard error, SD(β)/mean(SE). Higher values indicate greater regional dispersion relative to typical estimation uncertainty. A value of 1 provides a reference, rather than an exact null expectation, because standard errors differ across regions and regional estimates are correlated.

$I^2$ summarizes heterogeneity relative to the uncertainty of the regional estimates[18]. Higher values indicate greater variation beyond that attributed to estimation error under the calculation's assumptions. Because cortical gene sets share genes and participants, we used $I^2$ descriptively rather than interpreting it as an exact percentage of biological regional variation.

$$w_r = SE_r^{-2}, \qquad \bar{\beta}_w = \frac{\sum_r w_r \beta_r}{\sum_r w_r},$$

$$Q = \sum_r w_r (\beta_r - \bar{\beta}_w)^2, \qquad I^2 = 100 \ \max\left(0, \frac{Q-(K-1)}{Q}\right),$$

Where K is the number of cortical regions.

We tentatively adopted $I^2$ <25% as a threshold for low estimated regional heterogeneity, informed by the conventional interpretation of Higgins et al [18]. This threshold was used to guide presentation and interpretation, rather than to establish the presence or absence of regional variation. Summary figures use gray shading when either CNV profile falls below this threshold. Statistics involving a flagged profile, including cDEC and deletion–duplication correlations, are shown descriptively and interpreted cautiously; the other CNV profile is considered separately. Low regional heterogeneity does not imply an absence of burden associations.

#### 5.2 Cortical dosage-effect contrast (cDEC): construction and interpretation

The cortical dosage-effect contrast (cDEC) is defined on the complete deletion and duplication effect-size profiles, $\beta_{Del}$ and $\beta_{Dup}$, i.e. the 180 regional $\beta_1$ estimates from CC-GSBA. Each profile was standardized using its own mean and standard deviation across all regions: $z(Del)$ = ($\beta_{Del}$ − mean($\beta_{Del}$)) / SD($\beta_{Del}$), and likewise $z(Dup)$ for duplications, $cDEC = z(Del) - z(Dup)$.

Because each profile is standardized with statistics computed over all regions, cDEC is a relative measure. It describes how the cortical pattern of deletion effects differs from that of duplication effects. It is not a region-specific estimate, and its values cannot be interpreted independently of the full profiles. By construction, cDEC has a mean of zero. Because almost all effects are negative, more negative values of z(Del) or z(Dup) indicate relatively larger effects within that profile. Negative and positive cDEC values therefore indicate relative deletion and duplication weighting, respectively. Negative values mark parts of the cortex where deletion effects are relatively stronger within the deletion profile than duplication effects are within the duplication profile, and positive values mark the reverse.

Standardizing each profile separately was necessary because deletion effects are on average about twice as large as duplication effects. A raw difference, $\beta_{Del} - \beta_{Dup}$, would mainly reflect this overall difference in scale and would be dominated by the deletion profile. cDEC removes this difference in scale, so it compares where each profile is relatively strongest rather than how large the effects are. Differences in absolute effect size between deletions and duplications are reported separately **(Results; Supplementary Tables 3 and 5**).

#### 5.3 Sensorimotor-association hierarchy: definition and interpretation

The sensorimotor–association hierarchy (H) was represented by T1w/T2w ratio map[19–21]. T1w/T2w is an MRI measure of cortical myelin content that is highest in primary sensory and motor cortex and lowest in association cortex[19], and it is widely used as an anatomical marker of this hierarchy[20,21]. H was oriented so that values increase from sensorimotor to association cortex. We chose T1w/T2w because it comes from structural MRI, a data source separate from the gene-expression data used to build the cortical gene sets, and because it captures the main axis of gene-expression variation across the cortex[19]. The functional connectivity gradient[22] was used as one of the reference maps in the spatial decoding analysis, not as the primary hierarchy. Because almost all regional effects are negative, a larger effect on cognitive ability corresponds to a more negative β. A positive $\rho(Del, H)$ therefore indicates that deletion effects are largest toward sensorimotor cortex, and a negative $\rho(Dup, H)$ that duplication effects are largest toward association cortex. The contrast $T = \rho(Dup, H) - \rho(Del, H)$ tests directly whether the two profiles relate differently to the same hierarchy (T = 0 if they do not), while the cDEC–H correlation summarizes how the relative weighting of deletion and duplication effects changes along the hierarchy. All hierarchy analyses were repeated using the Schaefer and Desikan parcellations, with H and the effect-size profiles recomputed for each (**Supplementary Methods 3.2**).

#### 5.4 Spin-permutation testing

Spin permutations were used to account for spatial autocorrelation between neighbouring cortical regions. The deletion and duplication profiles are both estimated from the same 180 regional gene sets. Neighbouring regions have similar gene-expression profiles, so their gene sets overlap. Within each profile, the same genes, and therefore the same CNV carriers, contribute to the estimates of several nearby regions. As a result, both the deletion and the duplication profiles vary smoothly across the cortex, with neighbouring regions having similar effect sizes. When the two profiles are correlated, the 180 regions are therefore not independent observations, and a standard correlation test would overstate the significance of the deletion–duplication correlation.

The empirical statistic was compared with a null distribution generated by spherical rotations of the cortical data while preserving the spatial organization of the parcellation[23,24]. To create each null map, the centre of each region was placed on a sphere representing the cortical surface. The sphere was then randomly rotated, with the same rotation mirrored between the two hemispheres, and each region took the value of the region nearest to its new position. Applied to the duplication profile, each rotation keeps its smooth spatial pattern but places it at a random position relative to the deletion profile. We generated 10,000 rotated duplication profiles and recalculated the deletion-duplication correlation for each, giving the correlation expected from the two profiles' spatial smoothness alone.

Two-sided permutation P values were used for the primary spatial analyses, calculated as the proportion of null correlations as large as or larger than the observed correlation in absolute value. A significant result means that the deletion and duplication profiles are more strongly correlated than expected from their spatial smoothness alone. The same procedure was used for all other spatial correlations and for the alternative parcellations.

**6. Spatial decoding using normative cortical maps**

To contextualize the deletion, duplication and cDEC profiles, we compared them with 20 normative cortical maps spanning molecular, metabolic, microstructural, developmental, evolutionary and functional properties of the cortex. Each reference map was mapped to the primary cortical parcellation and correlated with the complete CC-GSBA profiles using Spearman correlation. Spatial significance was assessed using two-sided spin permutations.

The normative maps were used for spatial decoding of the CC-GSBA profiles and were not used to define the primary sensorimotor-association hierarchy. Map sources, biological annotations, processing and multiple-testing correction are provided in **Supplementary Table 9.**

**7. Phenotype-specificity analyses**

To assess whether the cortical deletion and duplication profiles observed for cognitive ability were phenotype specific, we applied the same CC-GSBA framework to additional UK Biobank phenotypes. The final comparison set included cognitive and socioeconomic phenotypes, and selected physiological and biochemical comparison phenotypes.

For each phenotype, deletion and duplication effects were estimated independently across the same cortical gene sets. We assessed two complementary relationships. First, same-dosage correlations (Del-Del and Dup-Dup) were used to quantify similarity of cortical profiles across phenotypes. Second, the Del-Dup correlation was calculated within each phenotype to determine whether the relationship between the two dosage directions varied across traits.

Phenotypes were coded so that the direction of higher values was documented before cross-phenotype comparison. Sample sizes, field definitions, transformations, covariates and standardization are reported in **Supplementary Tables 1, 8**.

**8. Population-structure sensitivity analyses**

The primary burden models included ancestry principal components (10 PCs). We further assessed robustness to population structure using geographic birthplace coordinates (North birth coordinate; and East birth coordinate). These analyses were treated as complementary sensitivity tests rather than as evidence that all forms of residual population structure could be excluded.

Selected physiological and biochemical phenotypes were additionally analysed using the same cortical gene sets to assess whether similar spatial profiles were observed for traits without an expected direct relationship to cognitive performance.

**9. All of Us analyses**

We used All of Us[25] Controlled Tier data release v8 to assess cross-cohort generalization, sensitivity to CNV call set and sensitivity to minimum CNV size. We constructed an Array∩WGS call set by intersecting array-derived CNVs with internally generated WGS-derived CNVs, and separately analysed the programme-provided GATK-SV structural-variant call set. Deletions and duplications were analysed separately using the same cortical gene-set definitions.

Array-derived CNVs were called using PennCNV[26] and QuantiSNP[27], following a pipeline similar to that used for UK Biobank. Calls were retained when supported by both algorithms with at least 50% overlap. Internal WGS-derived CNVs were generated using gVCF2CNV[28], which extracts log R ratio and B-allele frequency signals from genomic VCF files. For the present analyses, these signals were used for CNV calling with PennCNV alone. We then intersected the retained array calls with these internal WGS calls to construct Array∩WGS. Thus, the two-caller overlap criterion applied to array calling, whereas Array∩WGS required support across the array and internal WGS call sets.

The Array∩WGS resource comprised 402,817 participants from v8. Household-income data were available for 329,406 participants. Cross-cohort generalization was assessed by comparing Array∩WGS profiles at a minimum CNV size of 50 kb with corresponding UK Biobank profiles for household income and educational attainment. Phenotype availability and analysis-specific exclusions determined the sample used in each analysis.

Separately, we analysed programme-provided GATK-SV calls from v8 in 95,405 participants, drawn from the 97,061 samples documented in the All of Us Genomic Quality Report (v8). These calls were generated using the GATK-SV ensemble structural-variant pipeline. Our harmonized call set retained PASS calls and predefined recurrent CNVs. Non-recurrent CNVs were required to overlap problematic regions by less than 50% and have a European-ancestry allele frequency <0.001. Gene-dosage events required complete coverage of a protein-coding transcript (Transcript_Overlap = 1). The source-call-set summary excluded four participants carrying very large CNVs. Counts of CNV events, unique CNVs, carriers, gene-dosage events and affected genes across minimum-size thresholds are provided in Supplementary Table 17. These counts describe the source call set before restriction to participants with household-income data.

GATK-SV sensitivity analyses focused on household income. We repeated CC-GSBA at minimum CNV sizes of 10, 20, 30, 40 and 50 kb and compared each lower-threshold profile with the ≥50-kb profile. The ≥50-kb GATK-SV profiles were also compared with UK Biobank profiles. Burden analyses used autosomal gene-dosage events and excluded participants with overlapping gene-level CNV representations. For call-set sensitivity, GATK-SV and Array∩WGS profiles were compared in matched participants with both call sets and the required phenotype and covariate data.

Profile agreement was summarized using Spearman correlation, Pearson correlation where reported, and sign concordance, defined as the proportion of cortical regions with effect estimates of the same sign. Spatial significance was assessed using the spin-permutation procedures described in Supplementary Methods 5. These analyses assessed cross-cohort generalization, CNV call-set sensitivity and minimum CNV size; they did not constitute a matched array-versus-WGS comparison within UK Biobank (Supplementary Figs. 10,11; Supplementary Note 5; Supplementary Tables 6,8,17).

### Supplementary Notes

#### Supplementary Note 1. Robustness to sample, cognitive measure, genes, loci, CNV size and frequency

We asked whether the profiles depended on the composition of the sample or on how cognition was measured. We repeated CC-GSBA separately in UK Biobank and in the five other cohorts combined, in the discovery and replication halves of the pooled sample, in 100 random split-halves, in participants of European genetic ancestry, in males and females, in participants aged ≤60 and >60 years, and for the in-person and online UK Biobank fluid-intelligence assessments. Each profile was compared with the primary profile of the same CNV type. Deletion profiles were concordant across subsets ($\rho$ = 0.44 to 0.995), as were duplication profiles ($\rho$ = 0.38 to 0.994; **Supplementary Table 6; Supplementary Figs. 2, 7, 16**). Concordance was lowest in the smallest subsets, the five other cohorts ($\rho$ = 0.44 and 0.38) and the online fluid-intelligence assessment for duplications ($\rho$ = 0.53), consistent with reduced precision. These results indicate that the profiles reflect neither the predominance of UK Biobank nor ancestry, sex, age or assessment mode.

We then asked whether the regional estimates were driven by a few genes, a few loci, large multigenic CNVs or recurrent CNVs, and whether they held for rare CNVs. Gene-wise effects cannot be estimated reliably from burden counts at the available carrier numbers. We therefore described how burden is distributed across genes, which does not depend on the outcome, and re-estimated the profiles after removing specific genes, loci or classes of CNV (**Supplementary Table 12; Supplementary Fig. 17**). Carriers of removed CNVs remained in the model with zero burden.

Burden was distributed across many genes: a median of 1,302 genes (deletions) and 2,562 genes (duplications) were affected per regional gene set, and the most frequently affected gene carried a median of 15% and 4% of regional burden. Concentration was unrelated to the hierarchy and to effect size ($|\rho| \leq 0.14$). Removing the 1 to 25 most frequently affected genes per set (up to 54% of deletion burden) lowered concordance gradually, from 0.86 to 0.62 for deletions and from 0.97 to 0.74 for duplications, the behaviour of a distributed signal rather than one carried by a few genes.

No single chromosome drove the profiles: leaving out each autosome in turn gave a median concordance of 0.997 for deletions and 0.995 for duplications, with a minimum of 0.77 for chromosome 15 in deletions (about 10% of deletion burden removed). No single genomic-disorder locus drove them either: removing each of 16 major recurrent loci in turn kept $\rho \geq 0.92$ for deletions and $\geq 0.90$ for duplications. Removing all 16 loci together (6,959 carriers) retained concordance of 0.76 and 0.80, and removing all 41 recurrent regions gave 0.81 and 0.81 **(Supplementary Tables 6, 13)**.

Excluding CNVs encompassing ≥5 to ≥10 genes preserved the deletion profile ($\rho$ = 0.89 to 0.97). Small duplications showed no spatial variation above noise (dispersion ratio ≈ 1), so the duplication profile was carried by larger duplications ($\rho$ = 0.94 for CNVs with ≥5 genes). Both profiles were reproduced among CNVs with a frequency ≤1 in 10,000 ($\rho$ = 0.54 and 0.73); at ≤1 in 25,000, the deletion profile fell to noise level while the duplication profile was retained (ρ = 0.66). The cDEC–hierarchy correlation remained positive in all 58 analyses (ρ = 0.11 to 0.52; **Supplementary Table 7**). These results indicate that the profiles reflect effects distributed across many genes and loci. Genomic-disorder CNVs contribute to the deletion profile as a group, as expected for high-penetrance multigenic deletions, but no individual locus or chromosome accounts for it.

**Supplementary Note 2. Statistics of the cortical effect-size profiles**

CC-GSBA yields two cortical profiles: the per-gene effects of deletions and of duplications on cognitive ability across 180 regions. We asked four questions: whether effects varied across regions, whether each profile followed the sensorimotor–association hierarchy, whether the two profiles localized differently along that hierarchy, and how deletion and duplication effects corresponded across regions.

We first asked whether the effect-size profiles showed variation beyond estimation uncertainty, which we refer to as regional heterogeneity. We assessed this with two descriptive summaries: the dispersion ratio, SD(β)/mean(SE), and $I^2$ (**Supplementary Methods 5.1; Supplementary Table 7; Supplementary Fig. 7a**). The dispersion ratio compares regional variation with typical estimation uncertainty, whereas $I^2$ summarizes heterogeneity relative to the uncertainty of individual regional estimates[18]. Neither measure identifies which regions differ.

The dispersion ratio was 2.7 for deletions and 1.9 for duplications, and $I^2$ was 85% and 70%, respectively. Together, these summaries supported appreciable regional heterogeneity in both cognitive profiles. In subsequent comparisons, we tentatively flagged profiles with $I^2$ <25% as having low estimated regional heterogeneity and interpreted their spatial organization cautiously[18]. This descriptive convention does not establish that a flagged profile contains only noise.

We then asked whether each profile followed the sensorimotor–association hierarchy (H = −T1w/T2w, increasing from sensorimotor to association cortex). Because effects are negative, a positive ρ(β, H) means larger effects toward sensorimotor cortex. Duplication effects increased toward association cortex (Spearman ρ = −0.27, two-sided p-spin = 0.008). Deletion effects tended to increase toward sensorimotor cortex, but this was not significant on its own (ρ = 0.31, p-spin = 0.11). The two profiles related differently to the hierarchy: the topographic contrast T = ρ(Dup, H) − ρ(Del, H) was −0.58 (p-spin = 0.007).

To describe how this difference is distributed across the cortex, we used the cortical dosage-effect contrast, cDEC = z(Del) − z(Dup). Each profile was standardized separately across all regions, so cDEC compares where each CNV type is relatively strongest, independent of the twofold difference in overall magnitude. Negative and positive values indicate relative deletion and duplication weighting, respectively (**Supplementary Methods 5.2**). cDEC is a relative contrast defined on the complete profiles, not a region-specific or absolute deletion-minus-duplication difference; absolute magnitudes are retained in the regional effect-size maps. cDEC increased from sensorimotor to association cortex (ρ = 0.37, p-spin = 0.006). T and the cDEC–H correlation are the primary tests of differential localization. T tests whether the two profiles differ in their association with H, and the cDEC–H correlation summarizes how their relative weighting changes along the hierarchy. Across the CNV-level sensitivity analyses in Supplementary Note 1, the cDEC–hierarchy correlation remained positive (ρ = 0.11 to 0.52; **Supplementary Table 7**).

We report the deletion–duplication correlation descriptively. The two profiles were negatively correlated (ρ = −0.26, p-spin = 0.004). This correlation is sensitive to any component the two profiles share (**Supplementary Note 1**). it becomes stronger when regions are first selected on significance (ρ = −0.40 among the 166 regions significant for either CNV type), because such selection induces a negative relationship between the two statistics.

**Supplementary Note 3. Robustness to gene-set definition and gene properties**

We asked whether the localization depended on how regional gene sets were defined. We varied the regional-expression threshold (Z > 0.2, 0.5, 0.8 and 1.0), the AHBA differential-stability filter (10th to 40th centile), the parcellation (Schaefer-400, left hemisphere; Desikan-34) and the expression resource (spatial co-expression modules; BrainSpan; Human Protein Atlas). The cDEC–hierarchy correlation was retained across the four thresholds (ρ = 0.37 to 0.42, p-spin ≤ 0.007), the differential-stability filters (ρ = 0.25 to 0.33) and the Schaefer-400 (ρ = 0.34, p-spin = 0.008) and Desikan-34 parcellations (ρ = 0.39, p-spin = 0.12, 34 regions; **Supplementary Table 7; Supplementary Figs. 5, 7**). With Desikan poles defined in the same way as for the Glasser parcellation (10 regions each), deletion effects differed between sensorimotor and association regions (exact Kolmogorov–Smirnov D = 0.80, p = 0.002) and duplication effects did not (D = 0.30, p = 0.79).

The 16 spatial co-expression modules and the two RNA-sequencing resources cannot be mapped onto the Glasser hierarchy, so we used them to test the deletion–duplication relationship and broad sensorimotor-versus-association localization rather than as parcel-wise replications (Supplementary Figs. 3,4 and 6). Across regions significant for either

CNV type, deletion and duplication effects were negatively related in all three (Spearman ρ = −0.61 for modules, −0.55 for the Human Protein Atlas and −0.55 for BrainSpan; descriptive). For the Human Protein Atlas, we compared anatomical sensorimotor (10 regions) and temporoparietal association (9 regions) cores defined from sample labels before testing; for BrainSpan, we compared primary sensory and motor regions (4) with association regions (5). These comparisons were not significant (exact Kolmogorov–Smirnov p = 0.23 and 0.44 for the Human Protein Atlas; p = 0.43 and 0.75 for BrainSpan, where the smallest attainable p is 0.016) and are reported as descriptive.

We then asked whether the localization was explained by properties of the genes involved. Constraint of individual genes was only weakly related to how closely their cortical expression followed the hierarchy (LOEUF, ρ = +0.08; **Supplementary Fig. 18**). Genes whose expression follows the hierarchy are therefore not, as a group, more or less constrained. We stratified genes into non-overlapping bins of loss-of-function constraint (LOEUF), haploinsufficiency (pHaplo), triplosensitivity (pTriplo) and missense constraint, rebuilt the 180 regional gene sets within each bin, and repeated CC-GSBA. Constraint strongly modified effect magnitude: deletions and duplications of the most intolerant genes had the largest effects **(Supplementary Tables 3, 5)**. The cDEC–hierarchy correlation was retained within the most and least constrained LOEUF strata (ρ = 0.37 and 0.45) but was weak in the intermediate stratum (ρ = 0.11; **Fig. 5e; Supplementary Table 7**).

Among the other strata, the cDEC–hierarchy association was retained for five of six cell types (ρ = 0.29 to 0.44; astrocytes ρ = 0.13), for brain-elevated and other-tissue-elevated genes (ρ = 0.35 and 0.43), for the second and third developmental epochs (ρ = 0.28 and 0.38) and for rising and non-transitional trajectories (ρ = 0.47 and 0.21). It was weak or absent for genes with low tissue specificity, the first epoch, falling trajectories and the most haploinsufficiency-sensitive and missense-constrained strata (ρ = −0.10 to 0.11; **Supplementary Figs. 14, 15 and 19; Supplementary Table 7**). These results indicate that regional constraint covaries with the hierarchy and accounts for part of the deletion profile's relationship to it, but that the differential localization of deletion and duplication effects is retained across most gene properties.

**Supplementary Note 4. Phenotype specificity and population structure**

We asked whether this organization is a general property of CNV burden in these gene sets or specific to cognition-related traits. We applied the same CC-GSBA to educational attainment and household income, and to three blood biochemistry measures (calcium, LDL and total cholesterol; **Supplementary Fig. 9; Supplementary Table 8**). We compared same-dosage profiles across traits and the deletion–duplication relationship within each trait.

Cognition-related traits shared the cognitive profiles (Spearman ρ with the cognitive-ability profile = 0.88 and 0.83 for deletions and 0.25 and 0.38 for duplications, for educational attainment and household income, respectively; **Supplementary Table 6**). Despite approximately twice the sample size, negative controls (Blood biochemistry measures) and birthplace coordinates showed few regional burden associations after multiple-comparisons correction (significant regions out of 180, deletion/duplication: calcium, 2/0; LDL cholesterol, 3/1; total cholesterol, 10/3; birthplace north, 5/0; birthplace east, 0/0). The biochemistry measures showed no regional variation above sampling noise (dispersion ratio 1.00 to 1.09), so their hierarchy statistics were not interpreted. The deletion–duplication relationship differed within the cognition-related family: negative for cognitive ability (ρ = −0.26) and positive for educational attainment, and income (ρ = 0.39, and 0.51).

We then asked whether residual population structure could produce the pattern. The models adjust for ten ancestry principal components, and the profiles were unchanged in participants of European ancestry (ρ = 0.99 for both CNV types; Supplementary Note 1). Birth coordinates (UK Biobank fields 129 and 130) are geographically structured and therefore a sensitive negative control for residual stratification; we analysed them in the same way. They showed weak regional variation (dispersion ratio 1.03 to 1.52). Their cDEC–hierarchy correlations were of opposite sign to cognition (north ρ = −0.47, p-spin = 8E-4; east ρ = −0.21, p-spin = 0.020), so they did not reproduce the cognitive organization; we interpret them descriptively only. These results indicate that the organization is specific to cognition-related phenotypes and is not explained by residual geographic population structure, although no single analysis can exclude all forms of confounding.

**Supplementary Note 5. Generalization to All of Us: cohort, call set and CNV size**

We asked whether cortical burden profiles generalized to an independent cohort and were retained across CNV-calling approaches and minimum-size thresholds. Using All of Us v8, we compared household-income and educational-attainment profiles with the corresponding UK Biobank profiles. The primary All of Us call set was Array∩WGS (CNVs ≥50 kb), constructed by intersecting array-derived CNVs with internally generated WGS-derived CNVs. Array calls required support from PennCNV and QuantiSNP with at least 50% overlap, whereas internal WGS calls used gVCF2CNV-derived signals and PennCNV alone (Diop et al., 2026; Supplementary Methods 9).

Household-income profiles correlated between UK Biobank and All of Us for deletions (Spearman ρ = 0.54, two-sided p-spin = 2E-4; sign concordance 97%) and duplications (ρ = 0.44, p-spin = 2E-4; sign concordance 92%). Educational-attainment profiles also correlated across cohorts (ρ = 0.64 for deletions and 0.26 for duplications; p-spin = 1E-4 and 2E-3, respectively; Supplementary Fig. 10; Supplementary Tables 6,8).

We next assessed sensitivity to CNV calling using the programme-provided GATK-SV call set from v8, separately from our internal WGS calls. At a minimum CNV size of 50 kb, GATK-SV income profiles correlated with UK Biobank profiles for deletions (ρ = 0.66) and duplications (ρ = 0.56; both two-sided p-spin <5E-3). In matched participants, GATK-SV and Array∩WGS profiles correlated at ρ = 0.52 for deletions and 0.38 for duplications, with sign concordance of 95.0% and 65.6%, respectively.

We then lowered the minimum GATK-SV CNV size from 50 kb to 40, 30, 20 and 10 kb. In the harmonized source call set, lowering the threshold from 50 to 10 kb increased deletion carriers from 8,443 to 12,964 and duplication carriers from 32,306 to 37,360. Gene-dosage events increased from 28,538 to 34,745 for deletions and from 145,242 to 158,970 for duplications (Supplementary Table 17). These counts describe the source call set before restriction to participants with household-income data.

Income profiles remained broadly concordant across thresholds, although deletion-profile agreement decreased as smaller CNVs were included. Spearman correlations with the ≥50-kb profile were 0.98, 0.95, 0.91 and 0.74 for deletions and 0.99, 0.98, 0.97 and 0.97 for duplications at minimum sizes of 40, 30, 20 and 10 kb, respectively (Supplementary Fig. 11; Supplementary Table 6).

Together, these analyses support cross-cohort generalization of the income and educational-attainment profiles and broad consistency of income profiles across CNV-calling approaches and size thresholds. They provide complementary tests of robustness, rather than a matched array-versus-WGS comparison within UK Biobank.

**Supplementary Note 6. Energy metabolism and cortical expansion beyond the hierarchy**

Decoding with 20 normative maps suggested that deletion effects aligned with metabolic maps and duplication effects with maps of cortical expansion (**Fig. 3a,b; Supplementary Fig. 12; Supplementary Table 9**). Because most maps cover the hierarchy, we asked whether these associations held beyond H. We combined maps into two composites, energy metabolism (oxygen and glucose metabolism) and cortical expansion (evolutionary and developmental expansion), each the mean of the rank-based z-scores of its two maps. We computed partial Spearman correlations between each profile and each composite controlling for rank(H), with joint rotation of the composite and H (10,000 rotations).

After adjustment, deletion effects remained associated with energy metabolism (partial $\rho = -0.33$, p-spin = 0.037) and duplication effects with cortical expansion (partial $\rho = -0.34$, p-spin = 2E-4), whereas the cross-associations were small (duplications with energy metabolism, partial $\rho = -0.05$, p-spin = 0.64; deletions with cortical expansion, partial $\rho = -0.05$, p-spin = 0.83; **Supplementary Fig. 13; Source Data**). The interaction I_H = [ρ(Del) − ρ(Dup)]energy − [ρ(Del) − ρ(Dup)]expansion tests whether the two CNV types differ in their relationship with the two dimensions. I_H was −0.57 (p-spin = 0.014); without adjustment for H it was −0.99 (p-spin = 2E-4).

The interaction was reproduced in UK Biobank (I_H = −0.59, p-spin = 0.011) and in all eight internal data subsets (p-spin = 0.004 to 0.038). The duplication–expansion arm was the stronger of the two (p-spin ≤ 0.01 in every subset), whereas the deletion–energy arm was consistent in size (partial $\rho = -0.27$ to $-0.34$) but nominally significant in only five of nine subsets. In the five other cohorts combined, the interaction was not detected (I_H = −0.10, p-spin = 0.60). This sample was underpowered: its standard errors were 3.7 (deletions) and

3.1 (duplications) times those of UK Biobank, and the interaction expected after attenuation ($I\_H = -0.18$) lay inside the null interval (−0.31 to 0.34).

Mean effects across the 17 resting-state networks are shown in Fig. 3e (deletion–duplication $\rho = -0.52$, p-spin = 0.13). These results indicate that deletions and duplications relate to distinct features of cortical organization beyond the sensorimotor–association hierarchy.

**Supplementary Note 7. Specification of the Genome Burden term**

CNVs are frequently multigenic, so a CNV that affects a regional gene set usually also affects genes outside it. The Genome Burden term $X_2$ accounts for these co-carried genes, at the cost of absorbing some of their genuine effects. We asked how the choice of $X_2$ affects the profiles.

We compared three specifications (Supplementary Methods 4.5), that answer three distinct questions and are reported in **Supplementary Table 16 and Supplementary Fig. 23**: without $X_2$ (G0), $\beta_1$ is the total association of regional gene burden; with outside-set genes (G1, primary), $\beta_1$ is the regional effect conditional on co-carried genes; with the total gene count (G3), $\beta_1$ is the excess effect of a regional gene over the average gene.

The total gene count reproduced the primary profiles (Spearman $\rho = 0.997$ for deletions and 0.994 for duplications); it is nearly a reparameterisation of the primary model ($\beta_1 \approx \beta_{1,primary} - \beta_{2,primary}$; $r = 0.998$). Counting all protein-coding genes, rather than those with a constraint annotation, also reproduced them ($\rho = 0.96$ for both CNV types). Omitting $X_2$ increased effect sizes: median $\beta_1$ rose from −0.045 to −0.074 for deletions and from −0.022 to −0.038 for duplications, so conditioning attenuated $\beta_1$ by 39% and 44%. This attenuation equals the outside-set effect multiplied by the number of outside-set genes accompanying each regional gene.

That accompanying-gene count was nearly identical for deletions and duplications ($\rho = 0.85$ across regions) and unrelated to the hierarchy ($\rho \approx 0$). Omitting $X_2$ therefore adds a common component to both profiles, reflecting CNV structure rather than regional biology, and the profiles were moderately reordered ($\rho = 0.71$ and 0.52 with the primary profiles). The cDEC–hierarchy correlation was retained under all three specifications ($\rho = 0.32$, 0.37 and 0.38 for G0, G1 and G3). These results indicate that the primary model is the only specification that separates the effect of a region's genes from that of genes carried on the same CNVs, and that alternative definitions of genome burden give the same profiles.

**Supplementary Figures**

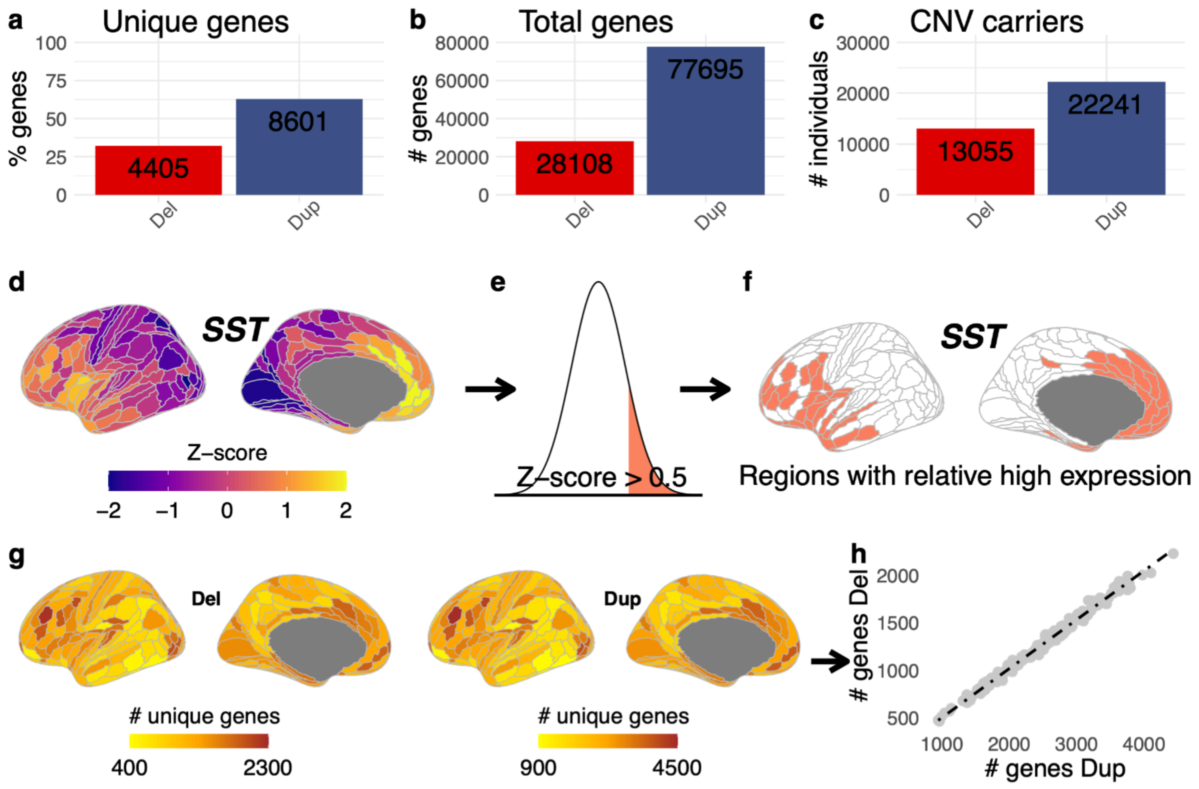

**Supplementary Fig. 1. Summary of CNV carriers and construction of cortical gene sets.**

Legend: **a-c)**, Percentage of unique genes, total number of genes (counting recurrence) and number of CNV carriers observed for deletions and duplications. Of the 13,678 genes with AHBA expression data that are detected in the brain (HPA), 32.2% were deleted and 62.9% duplicated at least once. **d-f)**, Identification of genes with relatively high expression across cortical regions. As an example, the expression of *SST* is z-scored across the 180 cortical regions (**d**), and thresholded at z > 0.5 (**e**), to identify the regions where it is relatively highly expressed (**f**). This procedure was applied across all 13,678 genes, resulting in 180 lists of relatively highly expressed genes corresponding to each cortical region of the Glasser parcellation. **g)**, The number of unique deleted and duplicated genes per cortical region. Separate colour bars show the different ranges of values for deletions and duplications. **h)**,

Number of unique genes per cortical region for deletions (y axis) versus duplications (x axis); each point is a region (n = 180).

AHBA: Allen Human Brain Atlas; HPA: Human Protein Atlas.

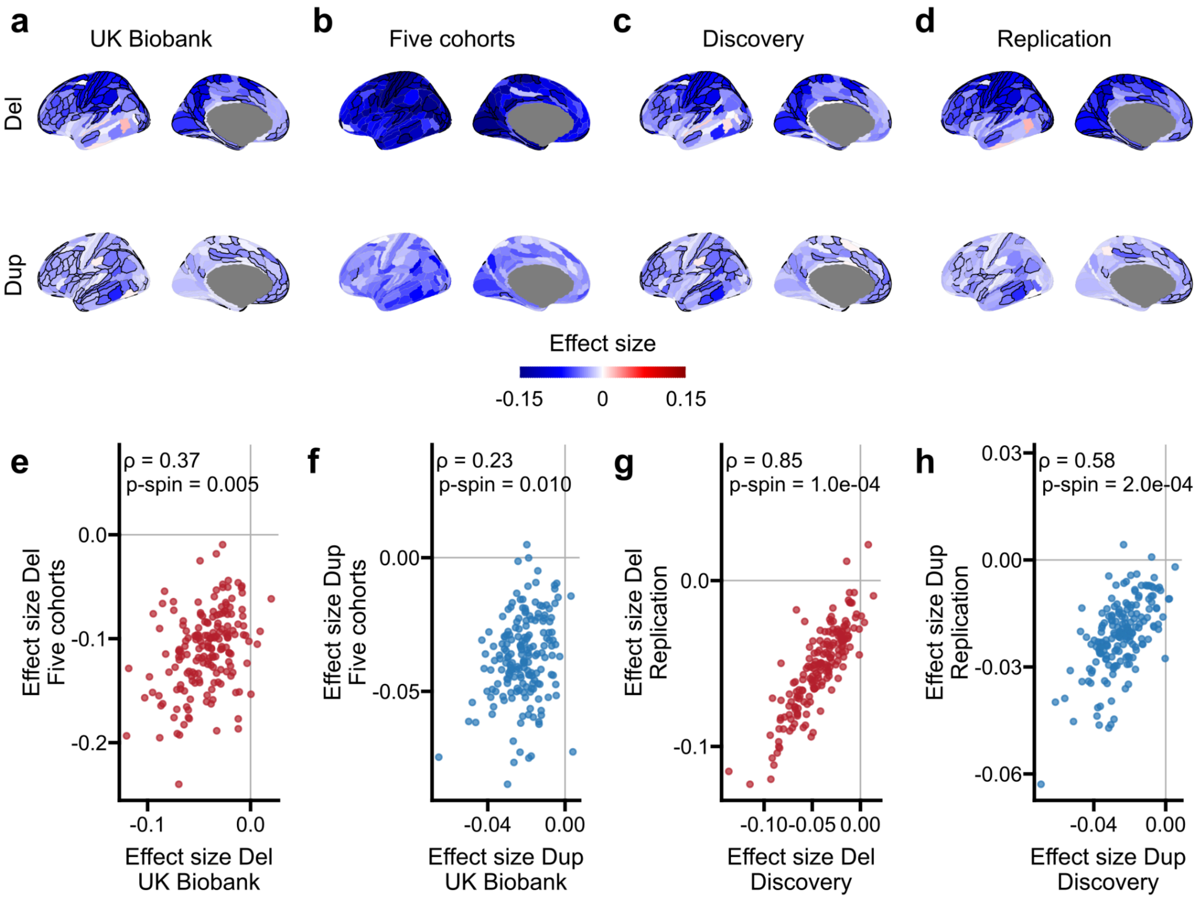

**Supplementary Fig. 2. Replication in data subsets.**

Legend: **a–d**) Deletion (top) and duplication (bottom) effect-size maps for UK Biobank and the five other cohorts combined (two non-overlapping datasets), and for the discovery and replication halves (a random split of the pooled sample). Black outline, region significant after Bonferroni correction (0.05/360). Shared colour bar, clipped at ±0.15. **e,f)** UK Biobank versus the five other cohorts combined, for deletions and duplications, respectively. **g,h)** Discovery versus replication halves of the pooled sample, for deletions and duplications, respectively. Each point represents one cortical region (n = 180). Spearman ρ and two-sided p-spin are shown; spin tests used 10,000 null rotations.Spearman ρ and two-sided p-spin

(10,000 null rotations) from **Supplementary Table 6**.

Del: deletion; Dup: duplication; p-spin: spin permutation p-value;

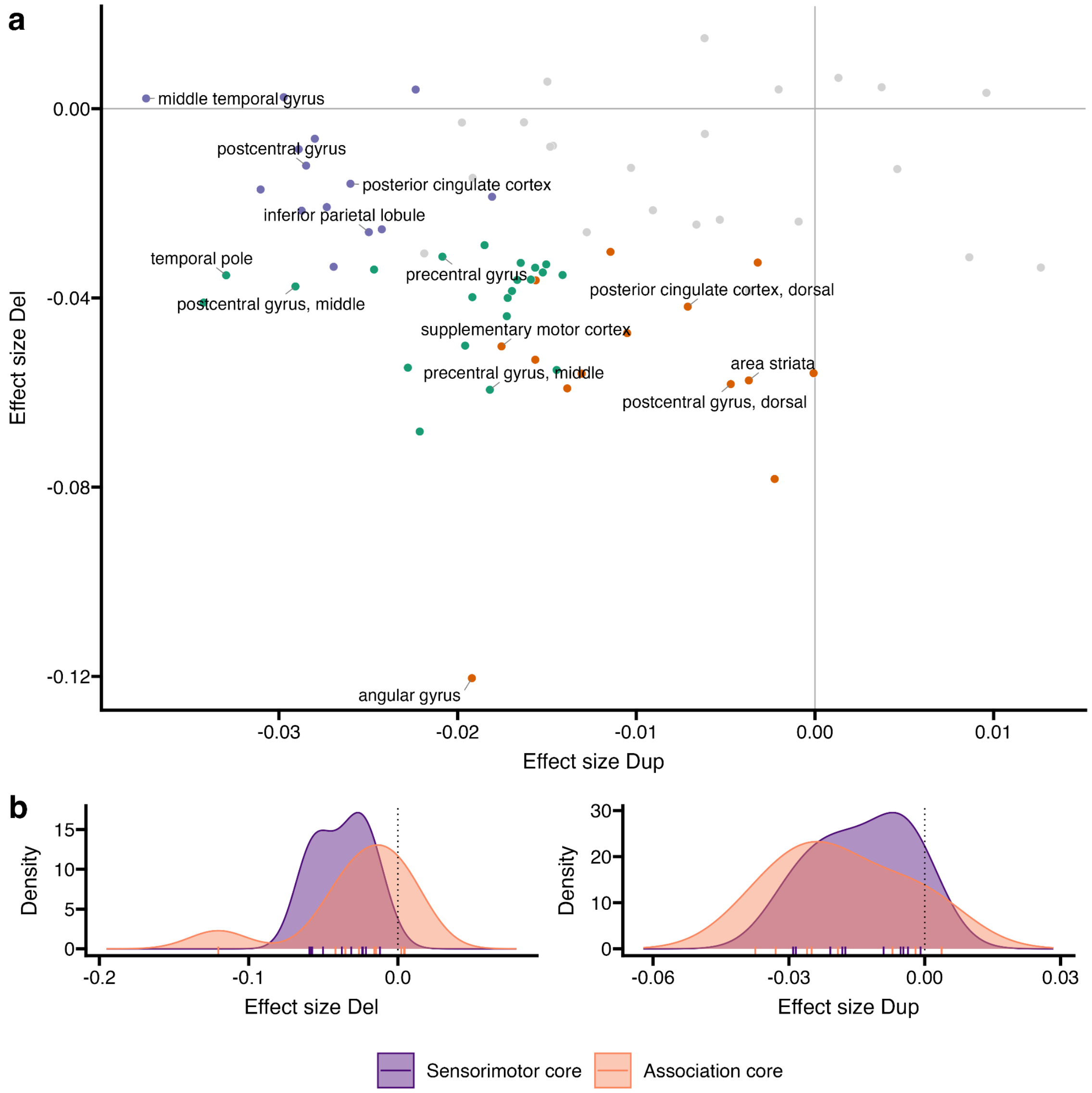

**Supplementary Fig. 3. Replication with Human Protein Atlas RNA-sequencing, 73 cortical regions.**

Legend: **a**) Deletion versus duplication effect sizes for the 73 Human Protein Atlas cortical regions, coloured by significance (Bonferroni 0.05/146): deletion-significant (orange), duplication-significant (purple), both (green), not significant (grey). Labels mark significant regions of the two sensorimotor and association axes cores. Deletion–duplication Spearman ρ = −0.01 across all regions and −0.55 across the 50 significant regions (descriptive). **b**) Effect-size distributions in the sensorimotor core and the temporoparietal association core; rugs show regions. The sensorimotor core has 10 regions: area striata; precentral and postcentral gyrus, whole, dorsal, middle and ventral; supplementary motor cortex. The association core has 9 regions: angular gyrus; inferior parietal lobule; middle and inferior temporal gyrus; temporal pole; precuneus; posterior cingulate cortex, whole, dorsal and ventral. Both cores were defined from the sample labels before testing. Exact two-sided Kolmogorov–Smirnov test (descriptive): deletions $D = 0.46$, $p = 0.23$, larger in the sensorimotor core; duplications $D = 0.37$, $p = 0.44$, larger in the association core. The Human Protein Atlas regions have no coordinates or surface parcellation, so no spin test was applied. See **Supplementary Note 3**.

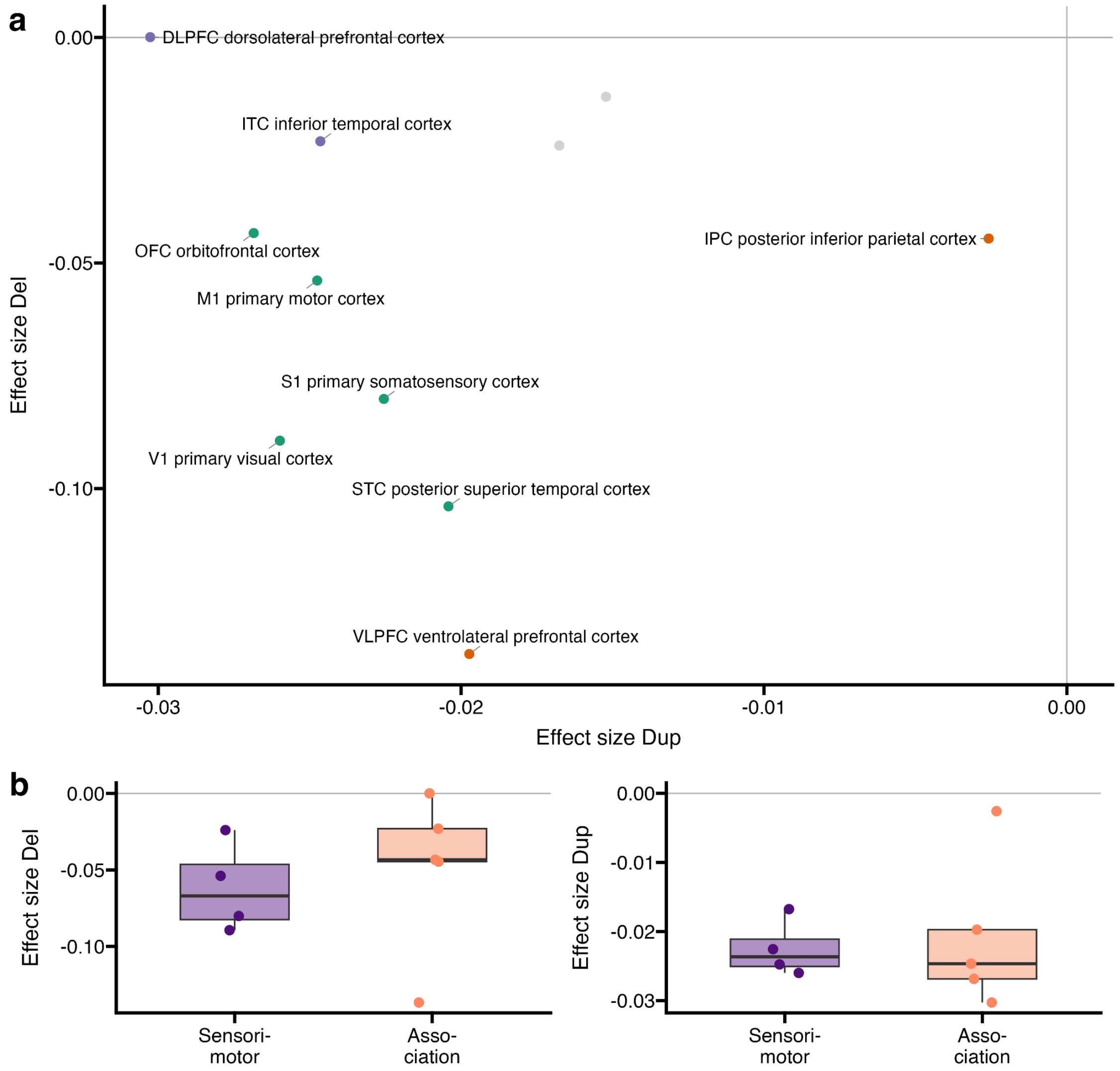

**Supplementary Fig. 4. Replication with BrainSpan adult RNA-sequencing, 11 cortical regions.**

Legend: **a**) Deletion versus duplication effect sizes for the 11 BrainSpan regions, coloured by significance class (Bonferroni 0.05/22); significant regions labelled. Deletion–duplication Spearman ρ = −0.12 across all regions and −0.55 across the 9 significant regions (descriptive). **b**) Regional effect sizes by pole: sensorimotor (V1, primary visual cortex; A1, primary auditory cortex; S1, primary somatosensory cortex; M1, primary motor cortex) and association (IPC, inferior parietal cortex; ITC, inferior temporal cortex; DLPFC, dorsolateral prefrontal cortex; VLPFC, ventrolateral prefrontal cortex; OFC, orbitofrontal cortex). ACC (anterior cingulate cortex) and STC (superior temporal cortex) are intermediate and not included in either pole. Points represent regions (sensorimotor, n = 4; association, n = 5). Boxes show the median and interquartile range; whiskers extend to the most extreme values within 1.5 × the interquartile range from the box edges (first and third quartiles). Exact two-sided Kolmogorov–Smirnov test: deletions D = 0.55, p = 0.43, with larger effects in sensorimotor regions; duplications D = 0.40, p = 0.75, with larger effects in association regions. With 4 versus 5 regions the smallest attainable p is 0.016, so the test is descriptive.

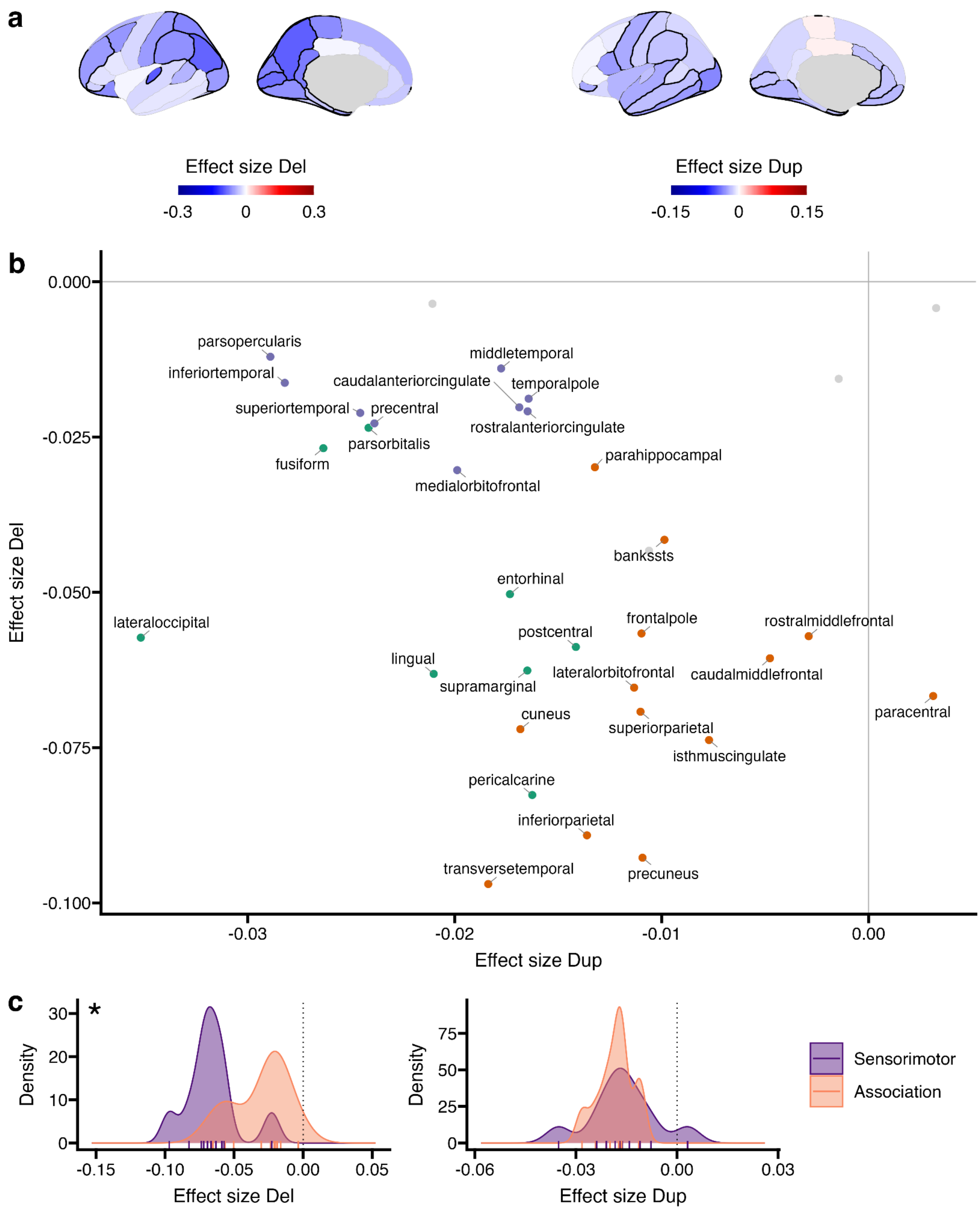

**Supplementary Fig. 5. Alternative AHBA gene sets: Desikan-34 parcellation.**

Legend: **a**) Deletion (±0.3) and duplication (±0.15) effect-size maps, left hemisphere; black outline, Bonferroni-significant region (0.05/68). **b**) Deletion versus duplication effect sizes for the 34 regions, coloured by significance class; significant regions labelled. Deletion–duplication Spearman ρ = −0.29 (p-spin = 0.17) across all regions and −0.50 (p-spin

= 0.015) across the 30 significant regions (descriptive; **Supplementary Table 7**). **c**) Effect-size distributions in sensorimotor and association regions (bottom and top 30% of H, 10 regions each). Exact two-sided Kolmogorov–Smirnov test (nominal, descriptive): deletions D = 0.80, p = 0.002, larger in sensorimotor regions; duplications D = 0.30, p = 0.79, larger in association regions.

Spearman correlations used two-sided spin-permutation tests with 10,000 null rotations. Sensorimotor–association Kolmogorov–Smirnov comparisons are descriptive because cortical regions are spatially dependent.

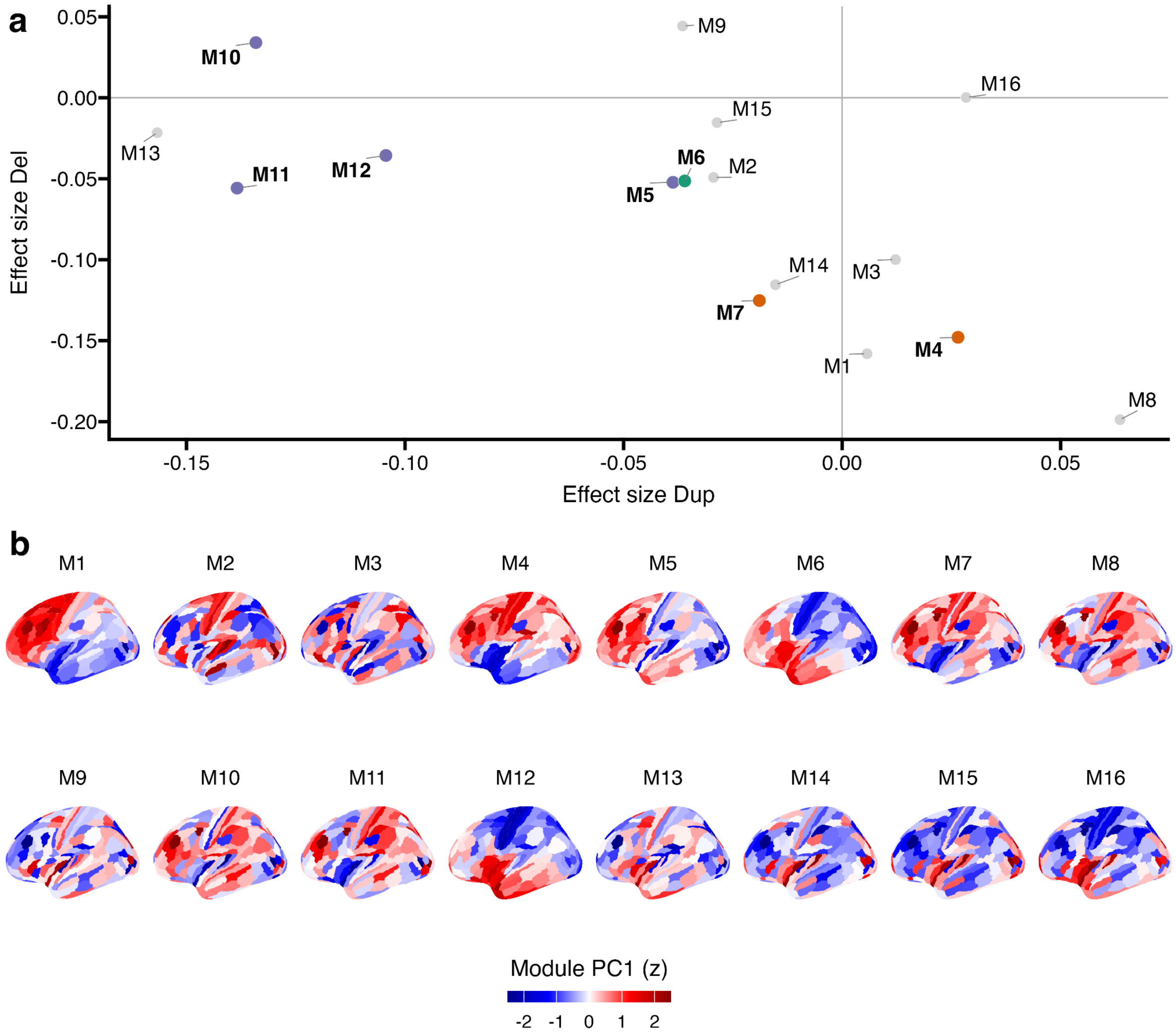

**Supplementary Fig. 6. Alternative AHBA gene sets: spatial co-expression modules.**

Legend: **a**) Deletion versus duplication effect sizes for the 16 AHBA co-expression modules, coloured by significance class (Bonferroni 0.05/32); all modules labelled, significant modules in bold. Deletion–duplication Spearman ρ = −0.53 across all modules (descriptive). Modules are not a spatial parcellation, so no spin test was applied. **b**) First principal component of each module (z-scored), Glasser-180, lateral view.

Spatial co-expression modules were obtained from Wagstyl et al. (2023).

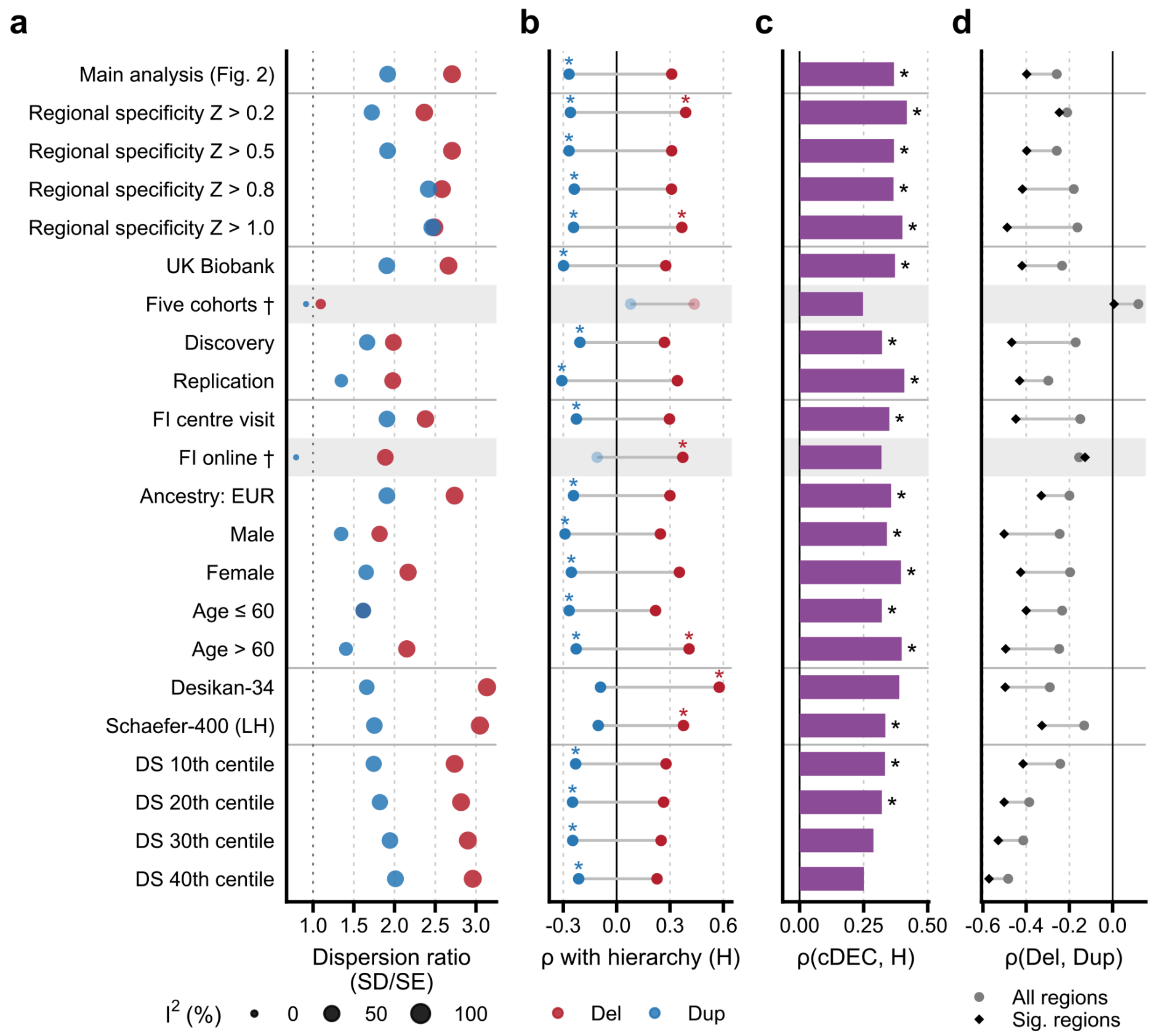

**Supplementary Fig. 7. Sensitivity analysis: cortical profile summary.**

Legend: **a**) Regional heterogeneity: dispersion ratio (SD of regional effect sizes / mean standard error; x-axis; dotted line, ratio = 1, the spread expected from estimation error alone) and $I^2$ (point size; percentage of variation across regions not due to estimation error; 0–100%). Both are descriptive; higher values indicate effect sizes that vary across cortex beyond noise. **b**) Spearman ρ of the deletion (red) and duplication (blue) profiles with the cortical hierarchy H (−T1w/T2w). c) ρ between the cortical dosage-effect contrast (cDEC) and H. d) Deletion–duplication correspondence across all regions (circle) and across regions significant for either CNV type (diamond). Rows: regional-specificity thresholds $Z > 0.2$ to 1.0; UK Biobank, five cohorts, discovery and replication; fluid-intelligence assessment (centre visit, online); ancestry (EUR), sex (male-only and female-only analyses) and age ($\leq$ 60, > 60); Desikan-34 and Schaefer-400 (left hemisphere) parcellations; AHBA differential-stability thresholds (10th to 40th centile). Asterisk, two-sided p-spin < 0.05 (nominal). Grey bands and daggers (†) identify analyses in which either the deletion or duplication profile showed limited regional heterogeneity ($I^2$ <25%). Spatial statistics involving the flagged profile, including cDEC–H and deletion–duplication correlations, are shown descriptively and interpreted cautiously (Supplementary Methods 5.1).

Spin tests used 10,000 null rotations. Values from **Supplementary Table 7**.

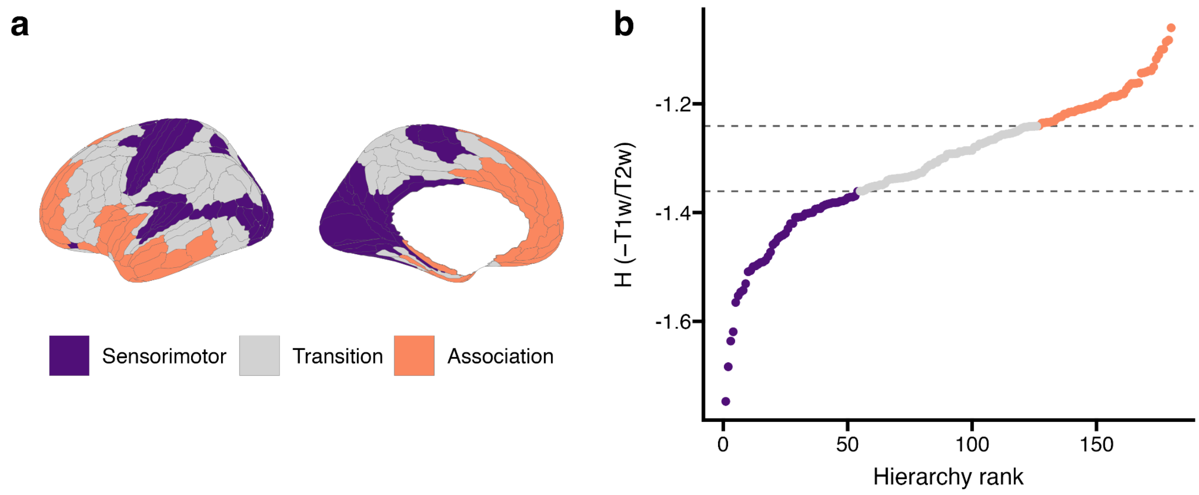

**Supplementary Fig. 8. Assigning cortical regions to the sensorimotor and association poles.**

Legend: **a**) Sensorimotor (≤ 30th percentile of H), transition and association (≥ 70th percentile) Glasser regions, 54 regions in each pole. **b**) H (−T1w/T2w) by rank; dashed lines, 30th and 70th percentiles. Pole membership in the **Source Data** and **Supplementary Table 11**.

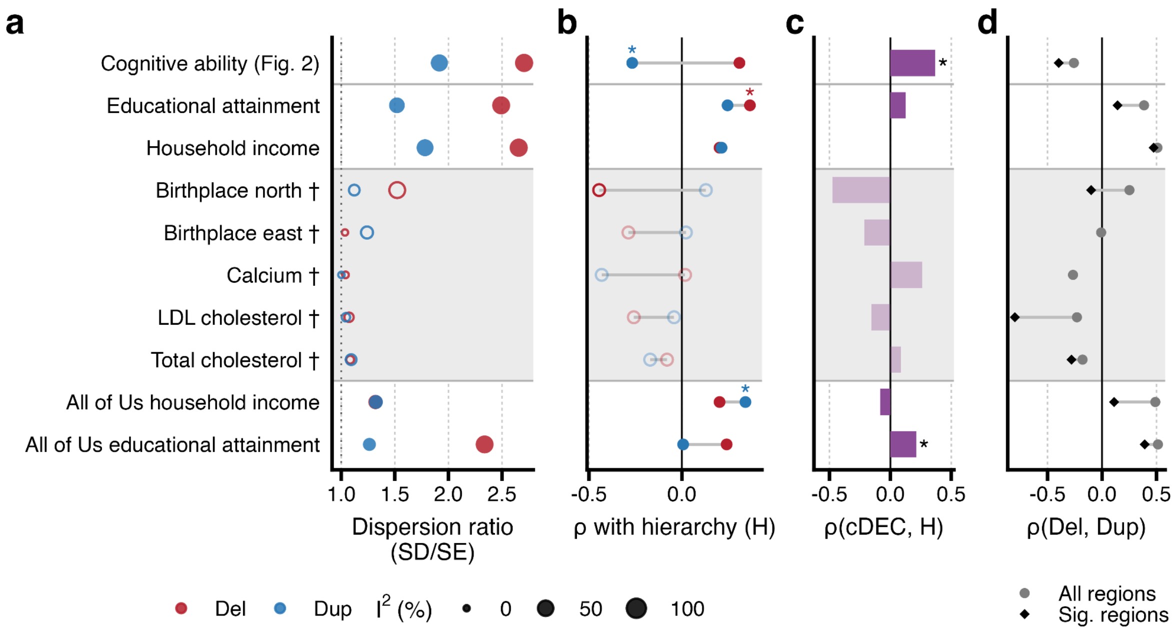

**Supplementary Fig. 9. Phenotypic specificity and generalization: profile summary.**

Legend: Columns **a–d** as in Supplementary Fig. 7. Despite approximately twice the sample size, negative controls (Blood biochemistry measures) and birthplace coordinates showed few regional burden associations after multiple-comparisons correction (significant regions out of 180, deletion/duplication: calcium, 2/0; LDL cholesterol, 3/1; total cholesterol, 10/3; birthplace north, 5/0; birthplace east, 0/0). Birthplace north and east, calcium, LDL and cholesterol showed limited regional heterogeneity (SD/SE ≈ 1; $I^2$ <25%); their cDEC–H values are not interpreted. Asterisk, two-sided p-spin < 0.05 (nominal). Grey bands and daggers (†) identify analyses in which either CNV profile showed limited regional heterogeneity ($I^2$ <25%); open symbols identify the affected profile. Spatial statistics involving that profile, including cDEC–H and deletion–duplication correlations, are shown descriptively and interpreted cautiously (Supplementary Methods 5.1).

Values from **Supplementary Tables 6-8**. See **Supplementary Note 4**.

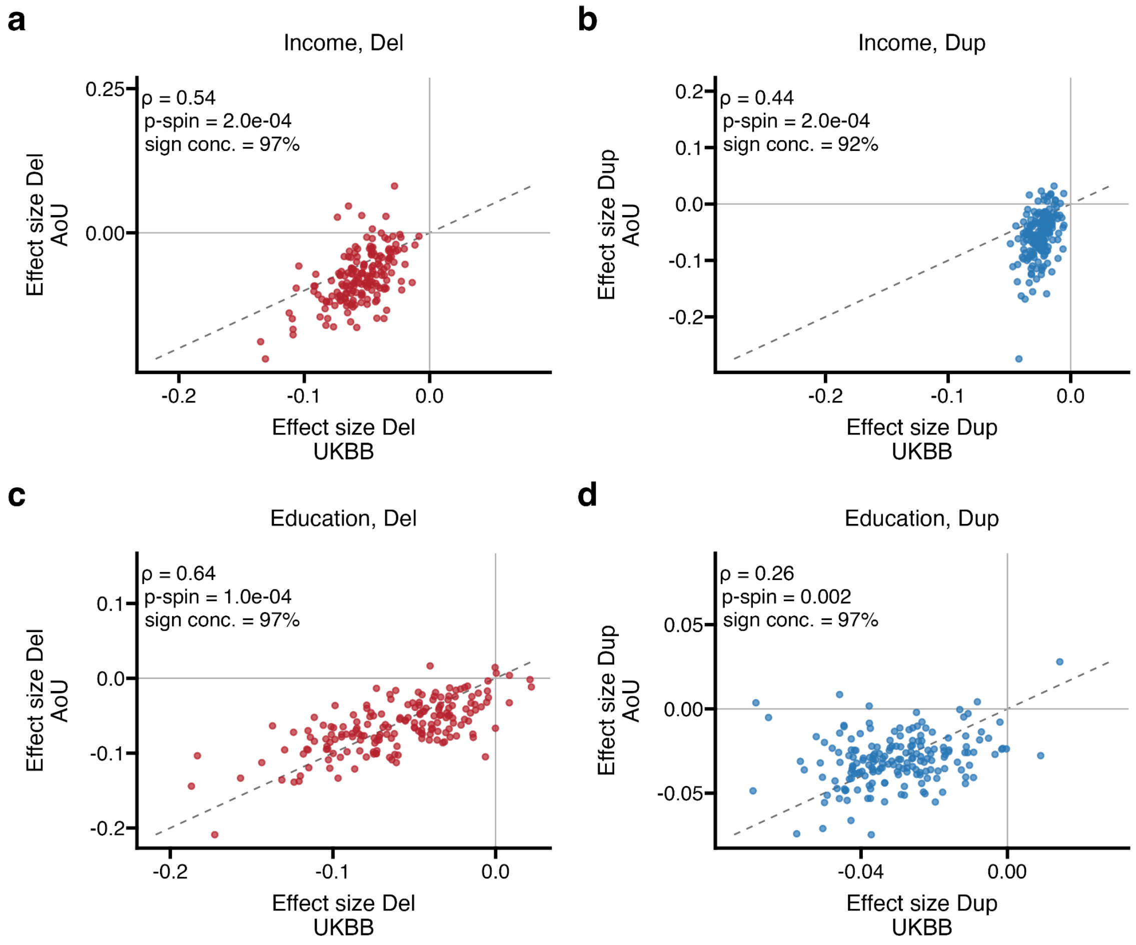

**Supplementary Fig. 10. Cross-cohort generalization in All of Us.**

Legend: UK Biobank versus All of Us (Array∩WGS call set, CNVs ≥ 50 kb), region-wise effect sizes for household income (**a**, deletions; **b**, duplications) and educational attainment (UK Biobank EA versus All of Us advanced degree; **c**, deletions; **d**, duplications). Each point is one region (n = 180); dashed line, identity. Printed: Spearman ρ, two-sided p-spin (10,000 null rotations) and sign concordance (percentage of regions with the same sign). Values from **Supplementary Table 6.** See **Supplementary Note 5**.

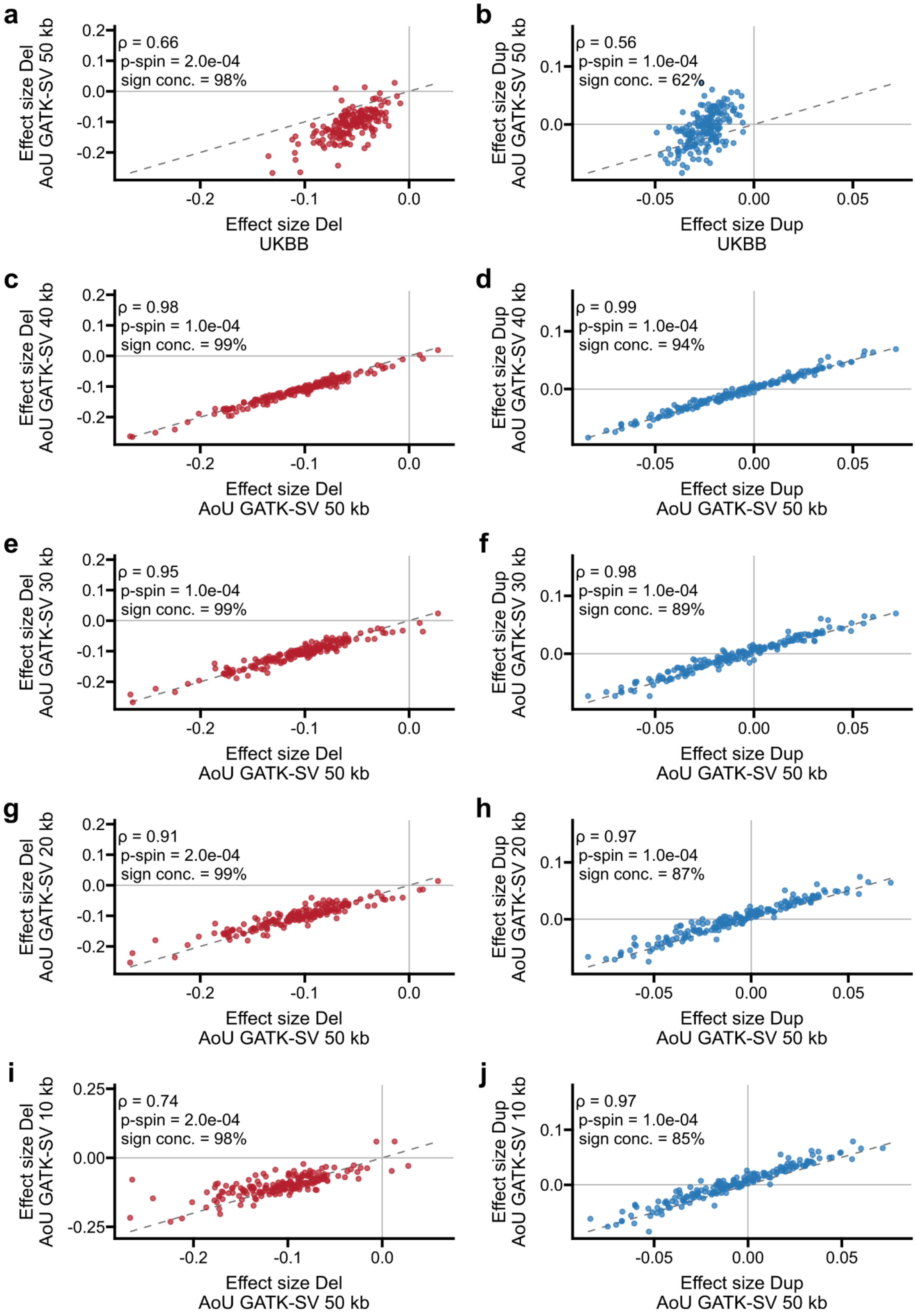

**Supplementary Fig. 11. CNV-calling technology and minimum CNV size in All of Us.**

Legend: Household-income effect-size profiles for deletions (left) and duplications (right). **a,b)** UK Biobank versus All of Us programme-provided GATK-SV calls with minimum CNV size ≥50 kb. **c,d), e,f), g,h) and i,j)** GATK-SV ≥50-kb profiles versus profiles at minimum sizes of 40, 30, 20 and 10 kb, respectively. Each point is a cortical region (n = 180); dashed lines indicate identity. Spearman ρ, two-sided p-spin and sign concordance are shown. Spin tests used 10,000 null rotations. Analysis samples and source-call-set counts are described in **Supplementary Methods 9 and Supplementary Table 17**.

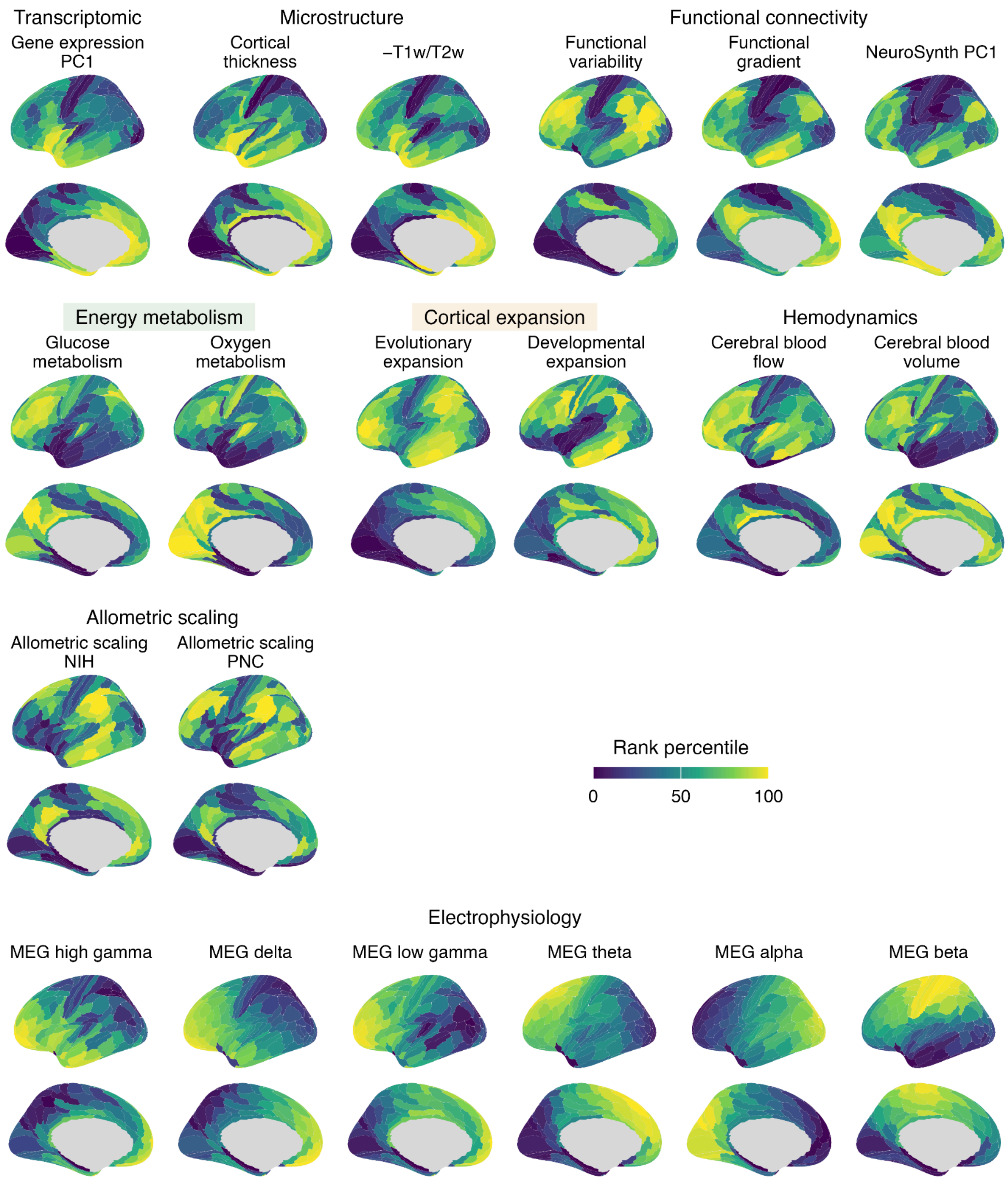

**Supplementary Fig. 12. Spatial decoding with 20 normative cortical maps.**

Legend: The 20 normative cortical maps[20,21]. The maps are grouped by domain: hierarchy and microstructure, energy metabolism, cortical expansion, hemodynamics, allometric scaling, and magnetoencephalography (MEG). Colours show each map's regional rank percentile, from low (0) to high (100). The −T1w/T2w map is oriented as the cortical hierarchy (H, Fig 2.), increasing from sensorimotor to association cortex. Map definitions and original sources are provided in **Supplementary Methods 6**.

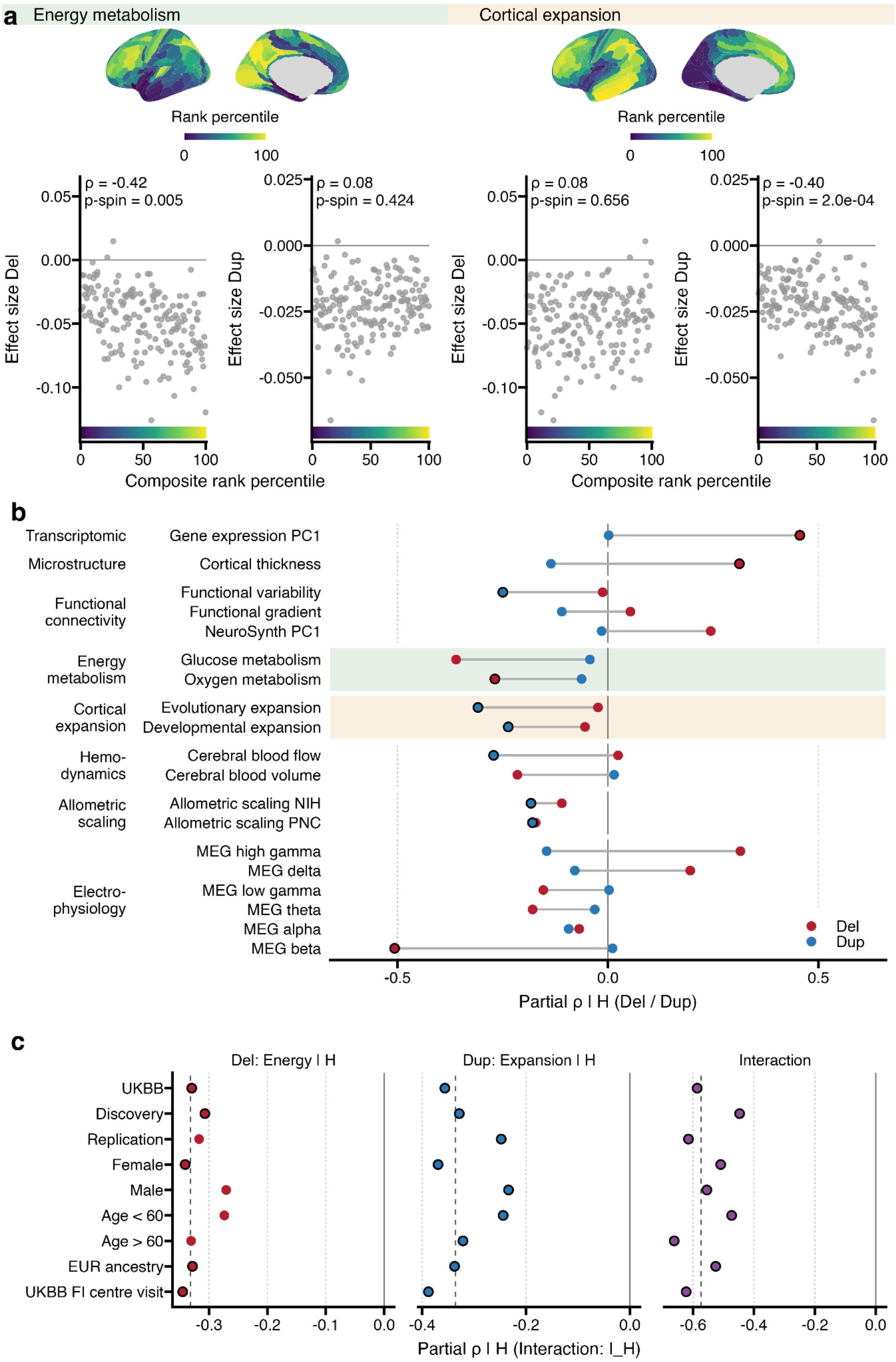

**Supplementary Fig. 13. Energy metabolism and cortical expansion: unadjusted associations, single maps and replication.**

Legend: a) Unadjusted version of Fig. 3c,d: raw deletion and duplication effect sizes versus the composite rank percentile for energy metabolism and cortical expansion (composites formed by averaging the rank-based Z-scores of the two component maps, oxygen and glucose metabolism, and evolutionary and developmental expansion; displayed as rank percentiles); the colour strip under the x-axis repeats the map scale. Spearman ρ and two-sided p-spin. b) Partial Spearman ρ | H of the deletion (red) and duplication (blue) profiles with each of 19 single maps (map and H rotated jointly); −T1w/T2w is omitted because it defines H; dark outline, p-spin < 0.05 (nominal). c) Consistency of the H-adjusted results across UK Biobank and overlapping internal subsets (discovery, replication, female, male, age ≤ 60, age > 60, EUR ancestry, UK Biobank fluid-intelligence centre visit): deletions with energy metabolism, duplications with cortical expansion, and the interaction I_H = [ρ(Del) − ρ(Dup)]energy − [ρ(Del) − ρ(Dup)]expansion (**Supplementary Note 6**); dashed line, full-sample value; dark outline, p-spin < 0.05 (nominal). I_H was not detected in the independent five-cohort sample (**Supplementary Note 6**). Each correlation is computed across n = 180 regions. Spin tests were two-sided and used 10,000 null rotations.

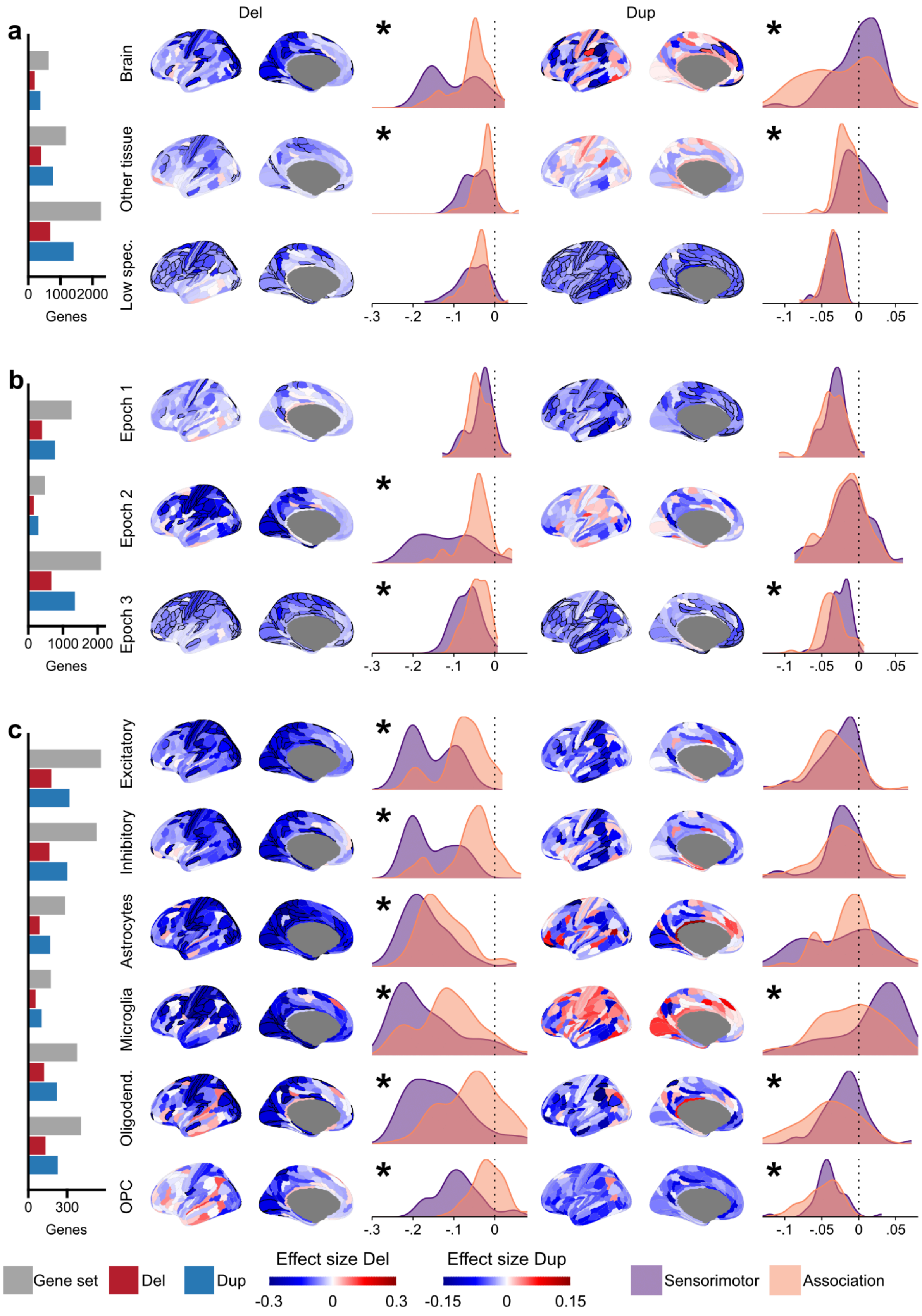

**Supplementary Fig. 14. Tissue specificity, temporal dynamics and cell types.**

Legend: Blocks, **a**) Human Protein Atlas tissue specificity (brain-elevated, other-tissue-elevated, low tissue specificity); **b**) BrainVar peak expression epoch 1 to 3; **c**) adult cell-type markers. Column 1: median regional gene-set size (grey) and median unique deleted and duplicated genes **(Supplementary Table 2)**. Other columns as in Fig. 4: effect-size maps per strata (Tissue specificity (a) and BrainVar epoch (b): $P < 0.05/1{,}080 = 4.63E\text{-}5$; Cell types (c): $P < 0.05/2{,}160 = 2.31E\text{-}5$) and sensorimotor-versus-association densities; asterisk, exact two-sided Kolmogorov–Smirnov test, FDR $q < 0.05$ across the 94 tests in **Supplementary Table 11**.

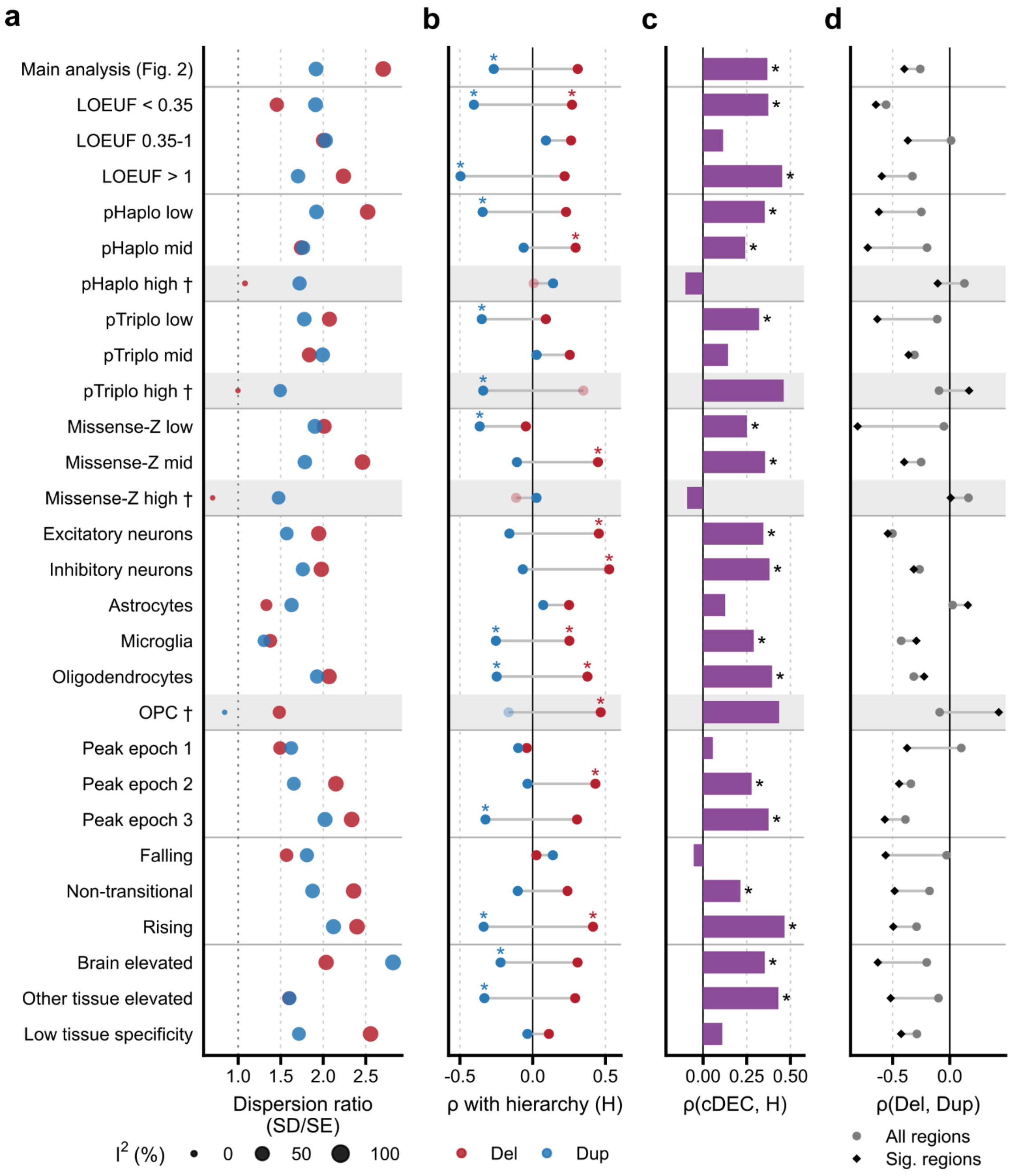

**Supplementary Fig. 15. Biological strata: profile summary.**

Legend: Columns as in Supplementary Fig. 7. Rows: LOEUF (3), pHaplo (3), pTriplo (3), missense Z (3), cell-type markers (6), BrainVar peak epoch (3) and trajectory (3), and Human Protein Atlas tissue specificity (3). Asterisk, two-sided p-spin < 0.05 (nominal). Grey bands and daggers (†) identify strata in which either the deletion or duplication profile showed limited regional heterogeneity ($I^2$ <25%). Spatial statistics involving the flagged profile, including cDEC–H and deletion–duplication correlations, are shown descriptively and interpreted cautiously (**Supplementary Methods 5.1**). Values from **Supplementary Table 7**.

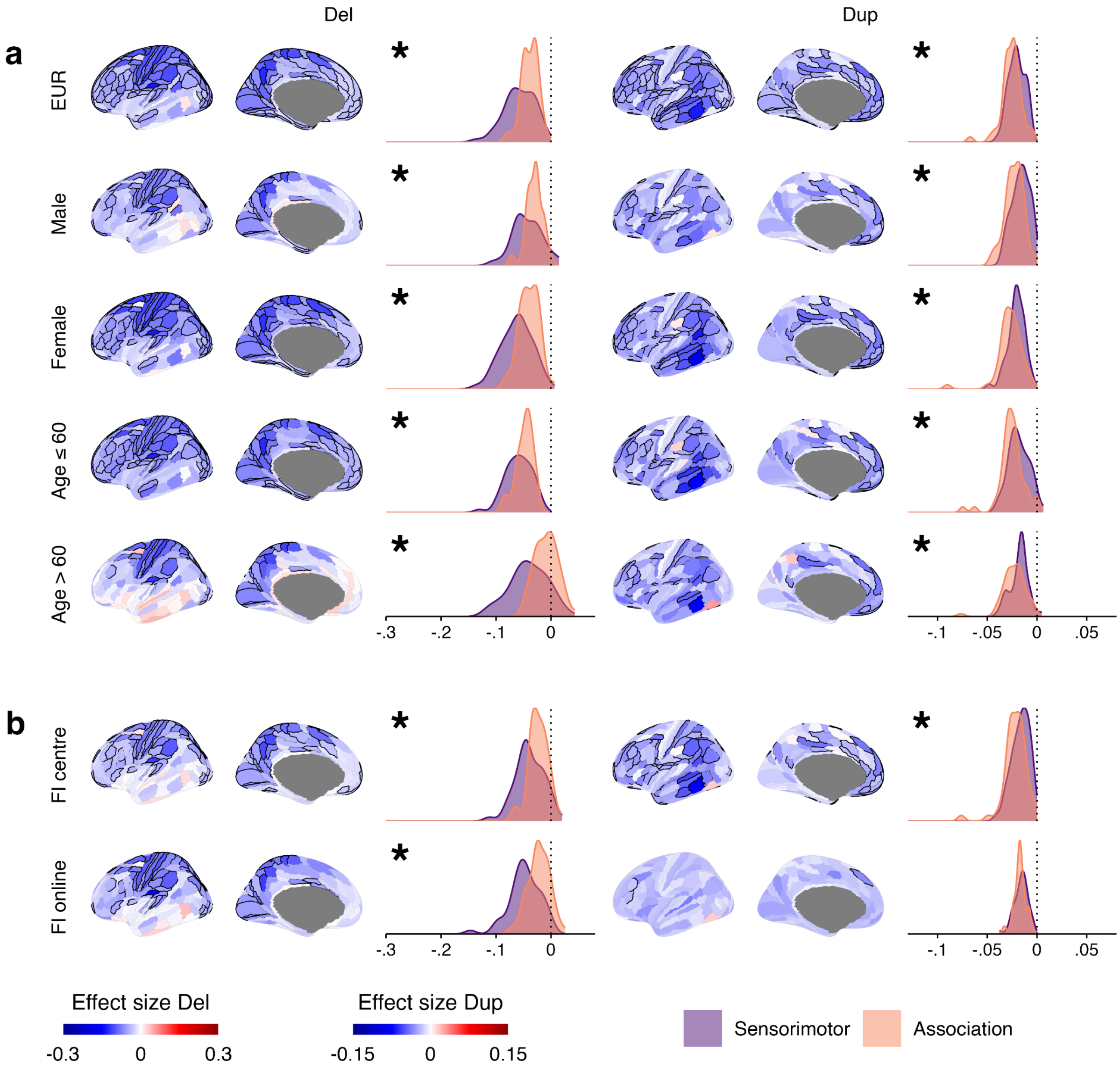

**Supplementary Fig. 16. Impact of age, sex, ancestry and assessment type.**

Legend: Rows: **a**) EUR ancestry, male-only, female-only, age ≤ 60 and age > 60; **b**) fluid intelligence assessed at the assessment centre and online (UK Biobank). Columns: deletion effect-size map (±0.3) and sensorimotor-versus-association density, then duplication (±0.15). Black outline, Bonferroni-significant region (0.05/360). Sensorimotor and association poles:

bottom and top 30% of H (cortical hierarchy; 54 regions each; Supplementary Fig. 8). Asterisk, exact two-sided Kolmogorov–Smirnov test, FDR q < 0.05 across the 94 tests in **Supplementary Table 11**. Dotted line, 0. Density windows: deletions −0.30 to 0.08; duplications −0.13 to 0.08.

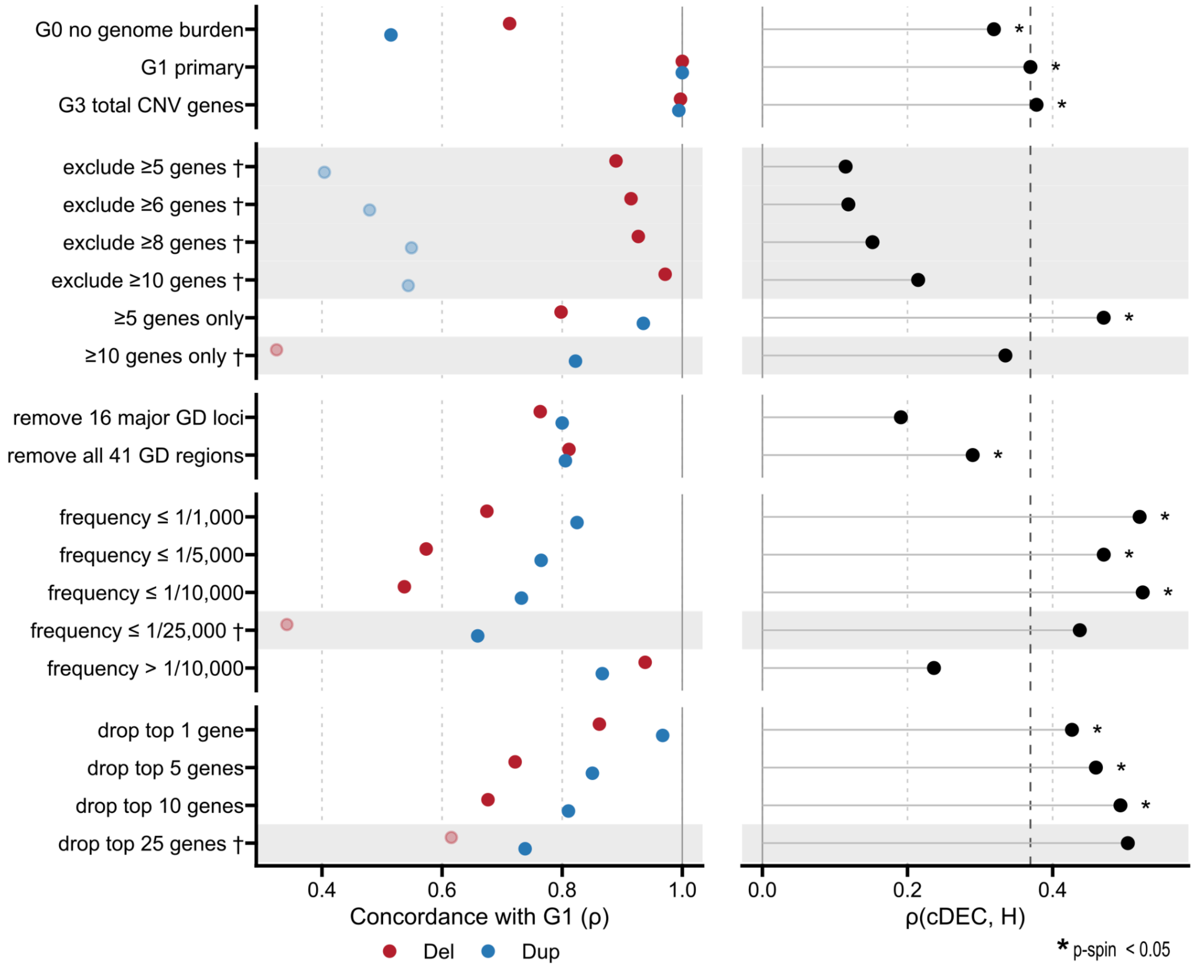

**Supplementary Fig. 17. Robustness to genes, loci, CNV size and frequency.**

Legend: Left: concordance (Spearman ρ) of each specification's deletion (red) and duplication (blue) profile with the matched G1 sensitivity rerun (primary Genome Burden specification refitted within the same robustness framework; not necessarily identical to the Fig. 2 coefficients). Right: ρ(cDEC, H) for each specification; dashed line, G1 value (0.37); asterisk, two-sided p-spin < 0.05 (nominal). Open symbols, profile with $I^2 < 25\%$ (**Supplementary Table 7**); cDEC point open when either CNV profile has $I^2 < 25\%$ ; statistics involving these profiles are interpreted cautiously **(Supplementary Methods 5.1).** Rows: Genome Burden specifications; gene-count caps; removal of the 16 major or all 41 genomic-disorder regions; CNV-frequency caps; removal of the most frequently affected genes. Gene-count caps, frequency caps and locus or gene removals remove CNV events, not participants: carriers of removed CNVs remain in the model with recalculated (zero) burden. Regional effects and retained burden counts are provided in **Supplementary Table 12**; leave-one-chromosome-out and leave-one-locus-out results in **Supplementary Tables 6 and 12**. Spin tests used 10,000 null rotations. See **Supplementary Note 1**.

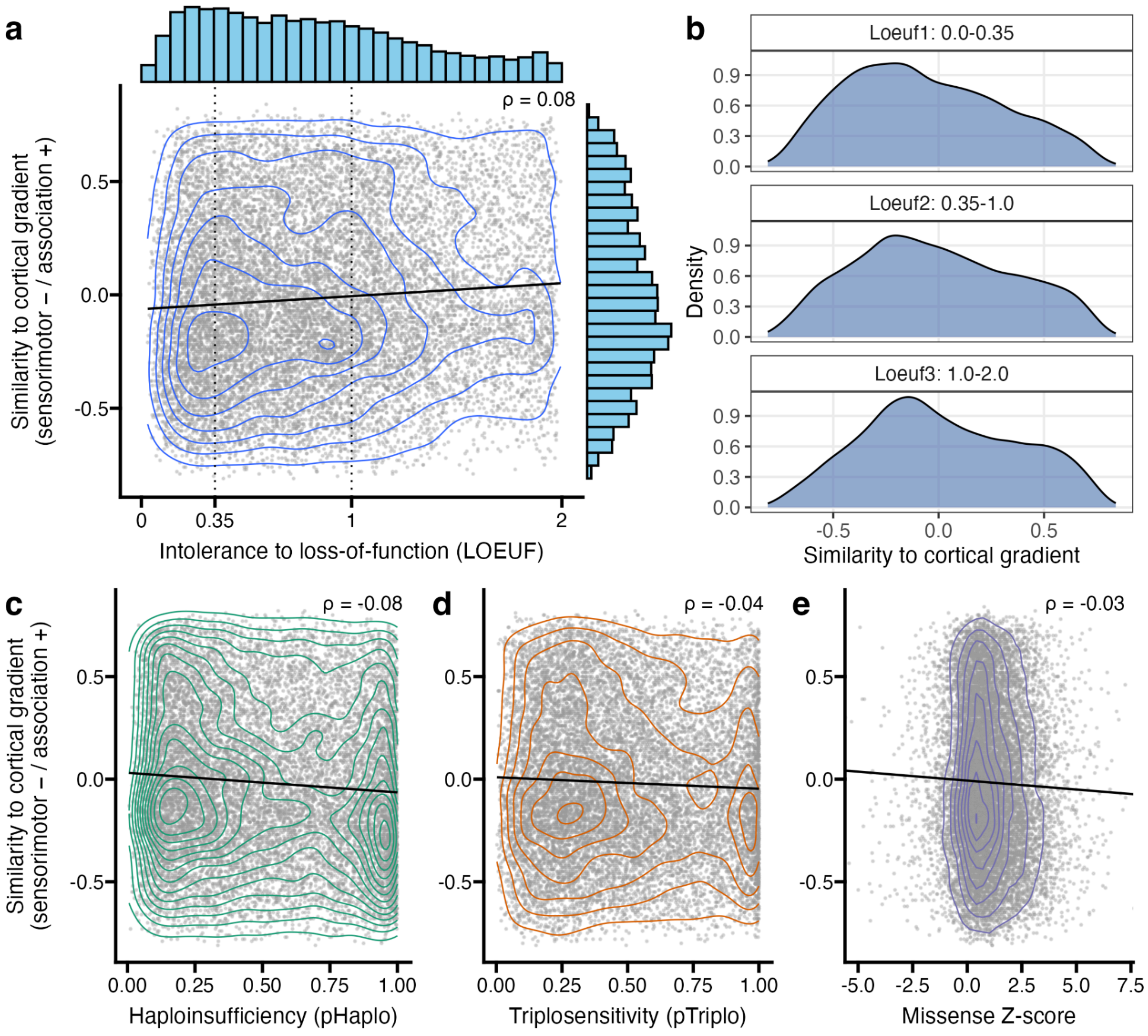

**Supplementary Fig. 18. Correlation between genetic constraint and similarity to the cortical gradient.**

Legend: **a**) Relationship between genetic constraint (LOEUF, x-axis) and similarity of each gene’s cortical expression profile to the sensorimotor–association hierarchy (H = −T1w/T2w, y-axis). Each point represents one gene; contours show two-dimensional density, and marginal histograms show the distributions of both measures. Positive similarity indicates alignment toward association cortex, whereas negative similarity indicates alignment toward sensorimotor cortex. **b)** Distributions of expression-profile similarity to H for genes stratified into three LOEUF categories: 0–0.35, 0.35–1.0 and 1.0–2.0. **c–e)** Corresponding relationships with probability of haploinsufficiency (pHaplo; **c)**, probability of triplosensitivity (pTriplo; **d)** and missense Z-score **(e)**. Spearman’s ρ is shown in each scatter plot. Lower LOEUF and higher pHaplo, pTriplo and missense Z-scores indicate greater constraint or dosage sensitivity.

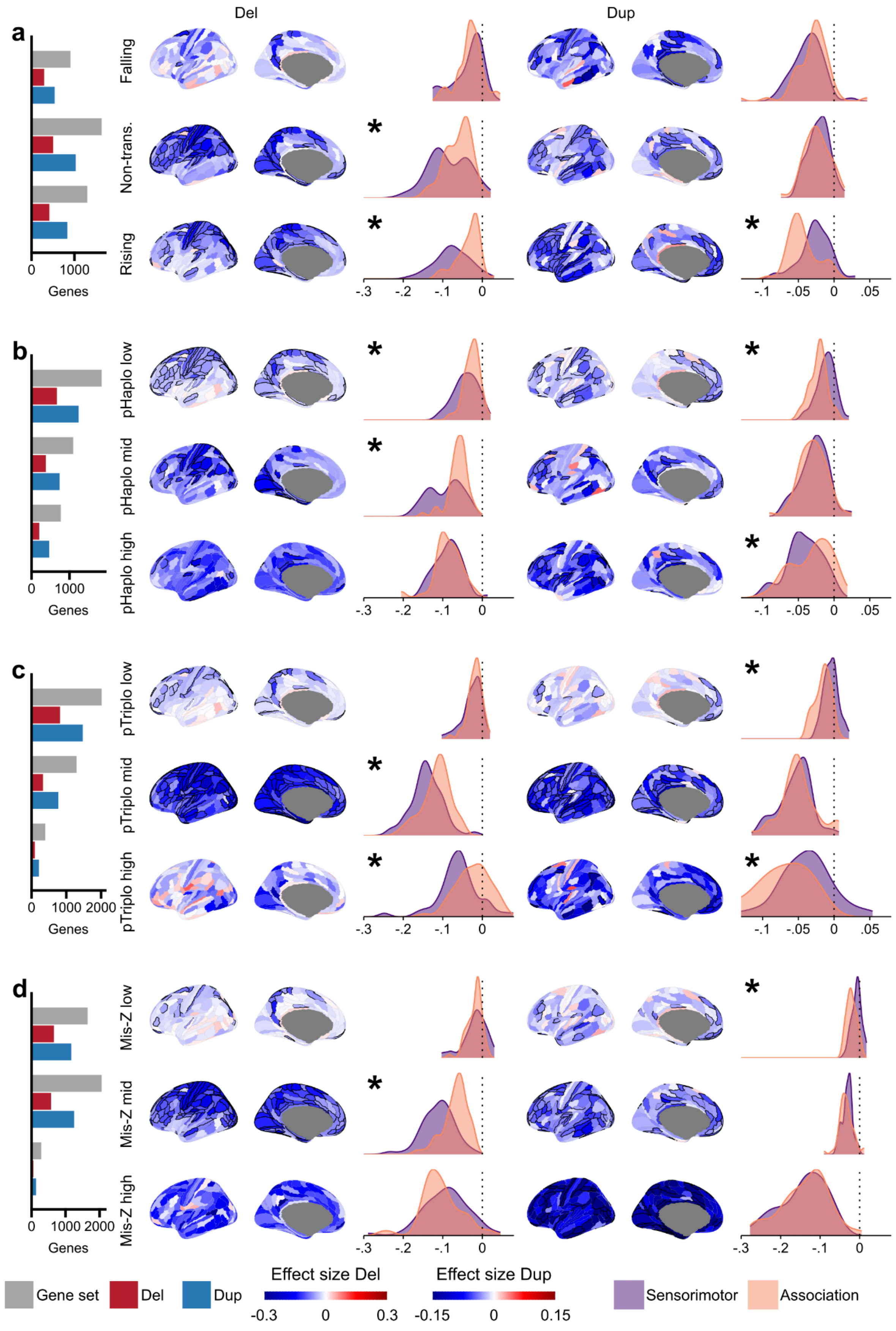

**Supplementary Fig. 19. Temporal trajectories, haploinsufficiency, triplosensitivity and**

**missense constraint.**

Legend: Blocks, **a)** BrainVar temporal trajectory (falling, non-transitional, rising); **b)** pHaplo; **c)** pTriplo; **d)** missense Z-score, each in tertiles. Layout as in Supplementary Fig. 14: effect-size maps per strata (all blocks, three strata: $P < 0.05/1{,}080 = 4.63E-5$) and sensorimotor-versus-association densities; asterisk, exact two-sided Kolmogorov–Smirnov test, FDR $q < 0.05$ across the 94 tests in **Supplementary Table 11**. Density windows: deletions −0.30 to 0.08; duplications −0.13 to 0.08, except the missense rows (−0.30 to 0.08) to show the full high-missense-Z duplication distribution.

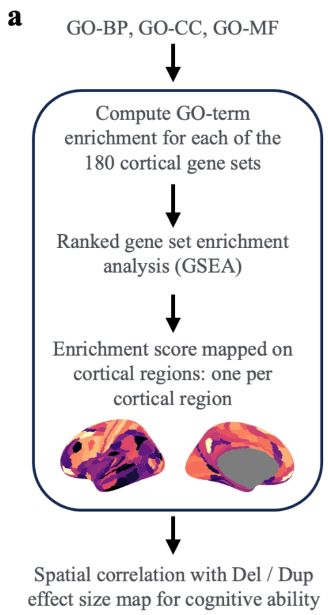

**Supplementary Fig. 20. Gene ontology enrichment across cortical gene sets**.

Legend: Workflow for i) computing gene ontology (GO) enrichments for each of the 180 cortical regions and its gene set using ranked gene set enrichment (fgsea package in R); ii) assembling normalized enrichment scores (NES) by GO-term name and cortical-region identifier to create a cortical profile for each GO term; and iii) correlating each retained profile with the complete deletion and duplication effect-size profiles of cognitive ability.

Enrichment was tested for 2,519 GO terms in three categories: GO-BP, biological process (1,938 terms); GO-CC, cellular component (272); GO-MF, molecular function (309). Nine GO terms showed significant enrichment in more than 5 cortical regions (FDR $q < 0.05$ across all tested term–region combinations); none of these nine profiles correlated significantly with the deletion or duplication effect-size profile (FDR $q < 0.05$ across the nine maps, separately for deletions and duplications). Spatial correlations used Spearman ρ and two-sided spin-permutation tests with 10,000 null rotations (Supplementary Tables 14, 15).

BP: biological process; CC: cellular component; GO: gene ontology; MF: molecular function; NES: normalized enrichment score.

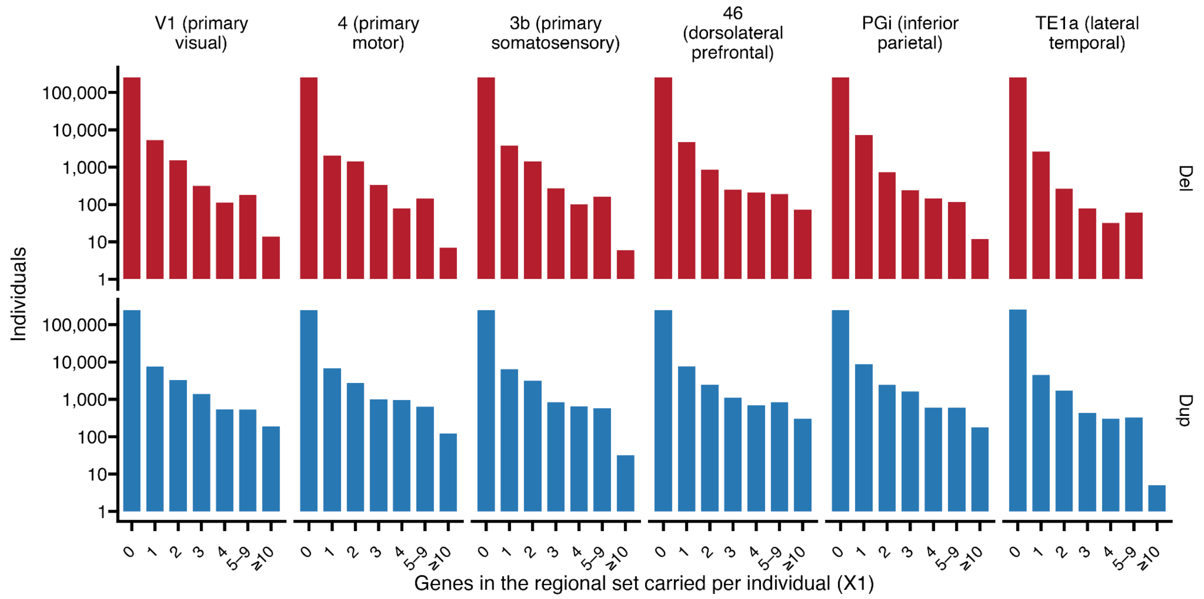

**Supplementary Fig. 21. Distribution of the number of genes hit by CNVs per individual across representative gene sets**.

Legend: Histograms showing the number of individuals (y-axis, logarithmic scale) by within-set gene burden (x-axis; $X_1$, number of genes in the set encompassed by the individual's CNVs; higher values grouped as indicated) for six representative cortical gene sets, separately for deletions (top row) and duplications (bottom row). Most individuals have zero burden within a given set, and relatively few carry multiple deleted or duplicated genes.

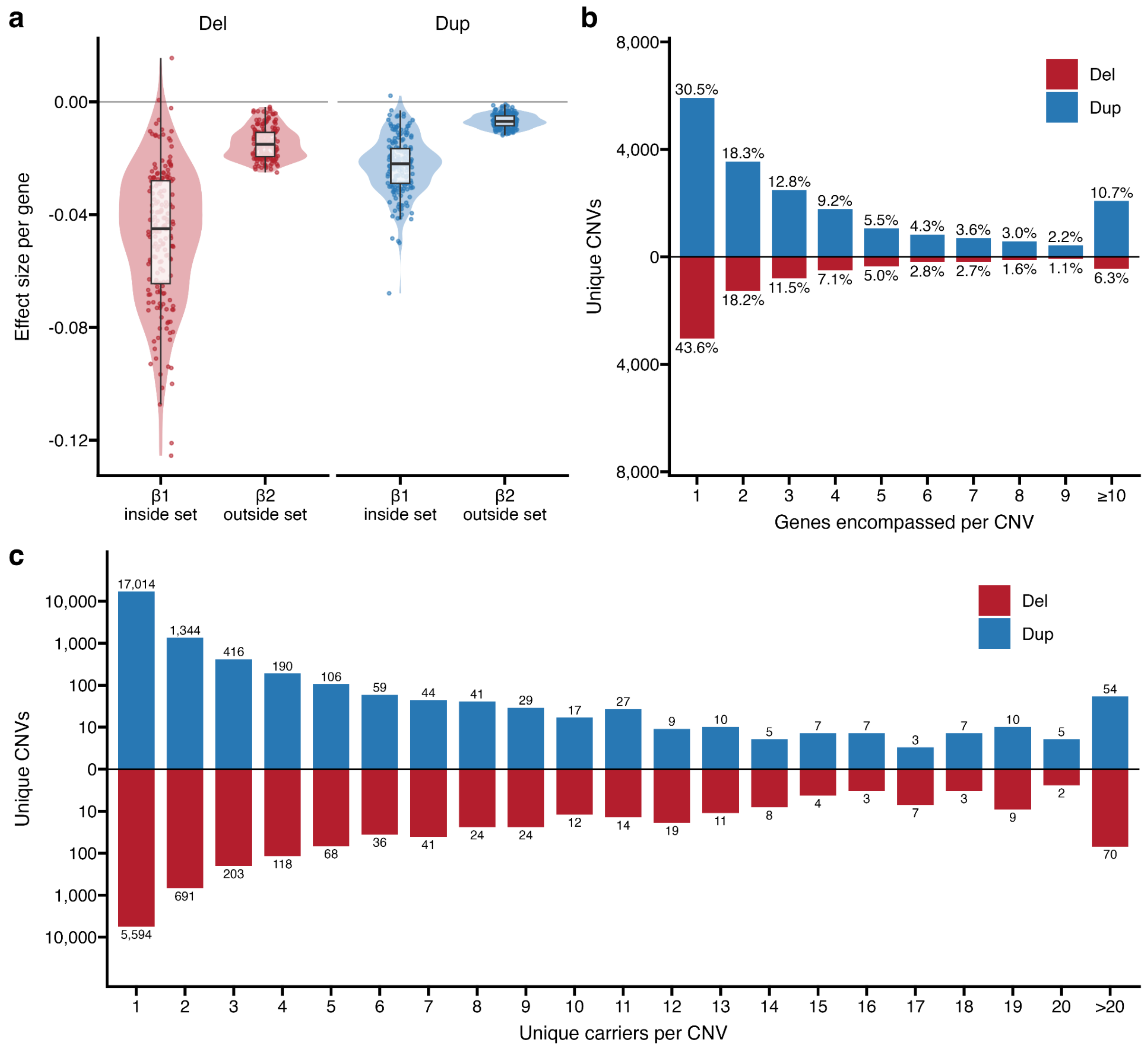

**Supplementary Fig. 22. Burden coefficients and CNV characteristics in the CC-GSBA of cognitive ability.**

Legend: **a**) Violin plots comparing the effect size estimates for genes inside the target gene set (β1) versus genes outside the set (β2) across all 180 cortical gene sets, shown separately for deletions (left) and duplications (right). Inside effects show greater variance than outside-set effects for both CNV types. The center line denotes the median, the box represents the interquartile range (IQR), and data points are shown as a jittered overlay to illustrate the full distribution of the gene sets' effect sizes. **b**) Number of protein-coding genes fully encompassed per distinct CNV. This is a mirror bar chart of deletions (red, below) and duplications (blue, above) in the analysed sample (N = 258,292), restricted to CNVs encompassing at least one gene. Labels give the percentage of distinct CNVs of each type. Deletions (n = 6,961): median 2 genes, mean 3.34; Duplications (n = 19,404): median 3 genes, mean 4.44; One gene: 43.6% of deletions and 30.5% of duplications; Ten or more genes: 6.3% of deletions and 10.7% of duplications. **c**) CNV recurrence. Distinct deletions (red) and duplications (blue), defined by identical coordinates and type, are binned by the number of unique individuals carrying each CNV. Bins run from 1 to 20 carriers, plus more than 20. Labels give the number of distinct CNVs, and the y-axis is pseudo-logarithmic. Most events were ultra-rare: 80.4% of deletions and 87.7% of duplications were carried by a single individual. 1.03% of deletions and 0.30% of duplications were carried by 20 or more individuals.

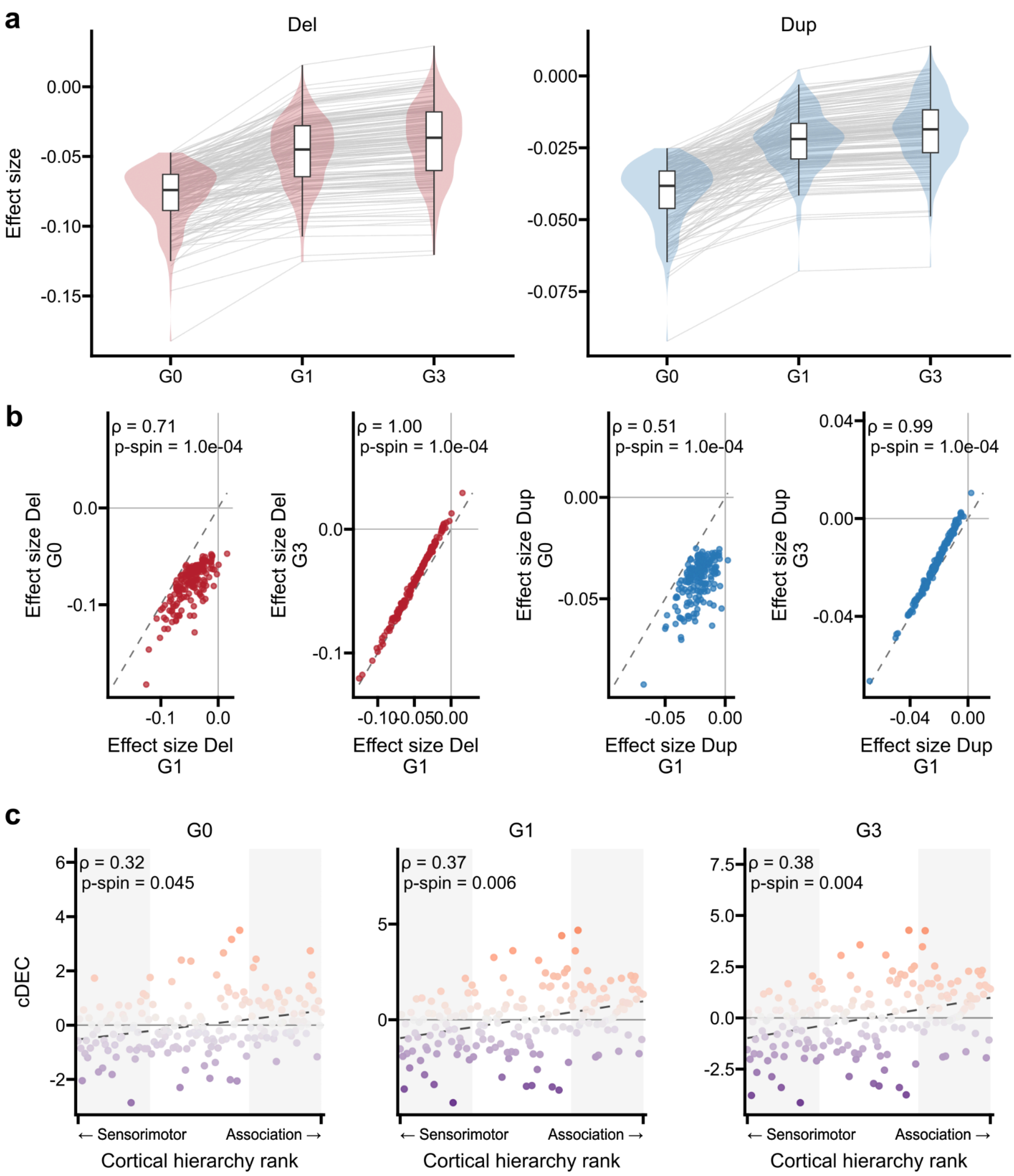

**Supplementary Fig. 23. Genome Burden specifications G0, G1 and G3.**

Legend: **a**) Regional effect sizes without a Genome Burden term (G0), with outside-set genes (G1, primary) and with the total gene count (G3), for deletions and duplications. Grey lines join the same region; violins with box plots show the distribution. **b**) G0 and G3 versus G1 effect sizes per region; dashed line, identity; Spearman ρ and two-sided p-spin from

**Supplementary Table 6**. **c**) cDEC versus cortical hierarchy rank for each specification, from sensorimotor to association cortex; shaded areas, bottom and top 30% of the hierarchy; points coloured by cDEC; dashed line, least-squares fit (visual aid); ρ(cDEC, H) and p-spin from **Supplementary Table 7**. See **Supplementary Note 7**.
